# Conservative route to genome compaction in a miniature annelid

**DOI:** 10.1101/2020.05.07.078311

**Authors:** José M. Martín-Durán, Bruno C. Vellutini, Ferdinand Marlétaz, Viviana Cetrangolo, Nevena Cvetesic, Daniel Thiel, Simon Henriet, Xavier Grau-Bové, Allan M. Carrillo-Baltodano, Wenjia Gu, Alexandra Kerbl, Yamile Marquez, Nicolas Bekkouche, Daniel Chourrout, Jose Luis Gómez-Skarmeta, Manuel Irimia, Boris Lenhard, Katrine Worsaae, Andreas Hejnol

## Abstract

The causes and consequences of genome reduction in animals are unclear, because our understanding of this process mostly relies on lineages with often exceptionally high rates of evolution. Here, we decode the compact 73.8 Mb genome of *Dimorphilus gyrociliatus*, a meiobenthic segmented worm. The *D. gyrociliatus* genome retains traits classically associated with larger and slower-evolving genomes, such as an ordered, intact Hox cluster, a generally conserved developmental toolkit, and traces of ancestral bilaterian linkage. Unlike some other animals with small genomes, the analysis of the *D. gyrociliatus* epigenome revealed canonical features of genome regulation, excluding the presence of operons and *trans*-splicing. Instead, the gene dense *D. gyrociliatus* genome presents a divergent Myc pathway, a key physiological regulator of growth, proliferation, and genome stability in animals. Altogether, our results uncover a conservative route to genome compaction in annelids, reminiscent of that observed in the vertebrate *Takifugu rubripes*.

---

Animals, and eukaryotes generally, exhibit a striking range of genome sizes across species^1^, seemingly uncorrelated with morphological complexity and gene content. This has been deemed the “C-value enigma”^2^. Animal genomes often increase in size due to the expansion of transposable elements (TE) (e.g. in rotifers^3^, chordates^4,5^ and insects^6^) and through chromosome rearrangements and polyploidisation (e.g. in vertebrates^7–9^ and insects^10^), which is usually counterbalanced through TE removal^11^, DNA deletions^12,13^ and rediploidisation^14^. Although the adaptive impact of these changes is complex and probably often influenced by neutral nonadaptive population dynamics^15,16^, genome expansions might also provide new genetic material that can stimulate species radiation^7^ and the evolution of new genome regulatory contexts^17^ and gene architectures^18^. By contrast, the evolutionary drivers of genome compaction are more debated and hypotheses are often based on correlative associations^1^, e.g. with changes in metabolic^19^ and developmental rates^20^, cell and body sizes^1,21^ (as in some arthropods^22,23^, flatworms^22^ and molluscs^24^), and the evolution of radically new lifestyles, such as powered flight in birds and bats^13,25^ and parasitism in some nematodes^26,27^ and orthonectids^28^. However, these correlations often suffer from multiple exceptions, e.g. not all parasites have small genomes^27^ neither does the insect with arguably smallest body size have a compact genome^29^, and thus they probably reflect lineage-specific specialisations instead of general trends in animal evolution. In addition, genomic compaction leading to minimal genome sizes, as in some free-living species of nematodes^30^, tardigrades^31,32^ and appendicularians^5,33^, apparently co-occurs with prominent changes in gene repertoire^34,35^, genome architecture (e.g. loss of macrosynteny^36^) and genome regulation (e.g. *trans*-splicing and operons^37–39^), yet these divergent features are also present in closely related species with larger genomes^5,32,40^. Therefore, it is unclear whether these are genomic changes required for genomic streamlining or lineage specialisations unrelated to genome compaction.

The marine annelid *Dimorphilus gyrociliatus* (O. Schmidt, 1857) (formerly *Dinophilus gyrociliatus*) has been reported to have a C-value (i.e. haploid genome size) of only 0.06–0.07 pg (~59–68 Mb)^41^, the smallest ever reported for an annelid^42^, and a haploid karyotype of 12 chromosomes^43^. *D. gyrociliatus* is a free-living meiobenthic species^44^ whose adults exhibit a strong sexual dimorphism, evident already during embryogenesis (Fig. 1a). The adult females are about 1mm long and display a typical albeit simplified annelid segmental body plan^45^ with only six segments, reduced coelom, and no appendages, parapodia or chaetae (Supp. Note 1). *D. gyrociliatus* males are however only 50 µm long, comprise just a few hundreds of cells, lack a digestive system, but still possess highly specialised sensing and copulatory organs^46^. Despite their miniature size, *D. gyrociliatus* retains ancestral annelid traits, such as a molecularly regionalised nervous system in the female^47,48^ and the typical quartet spiral cleavage^49^ (Fig. 1b). With only a handful of genomes sequenced (Supp. Table 1), annelids have retained ancestral spiralian and bilaterian genomic features^50^. Therefore, *D. gyrociliatus*, with its reduced genome size and small body, emerges as a unique system to investigate the genome architecture and regulatory changes associated with genome compaction and assess the interplay between genomic and morphological miniaturisation.

**Figure 1.**
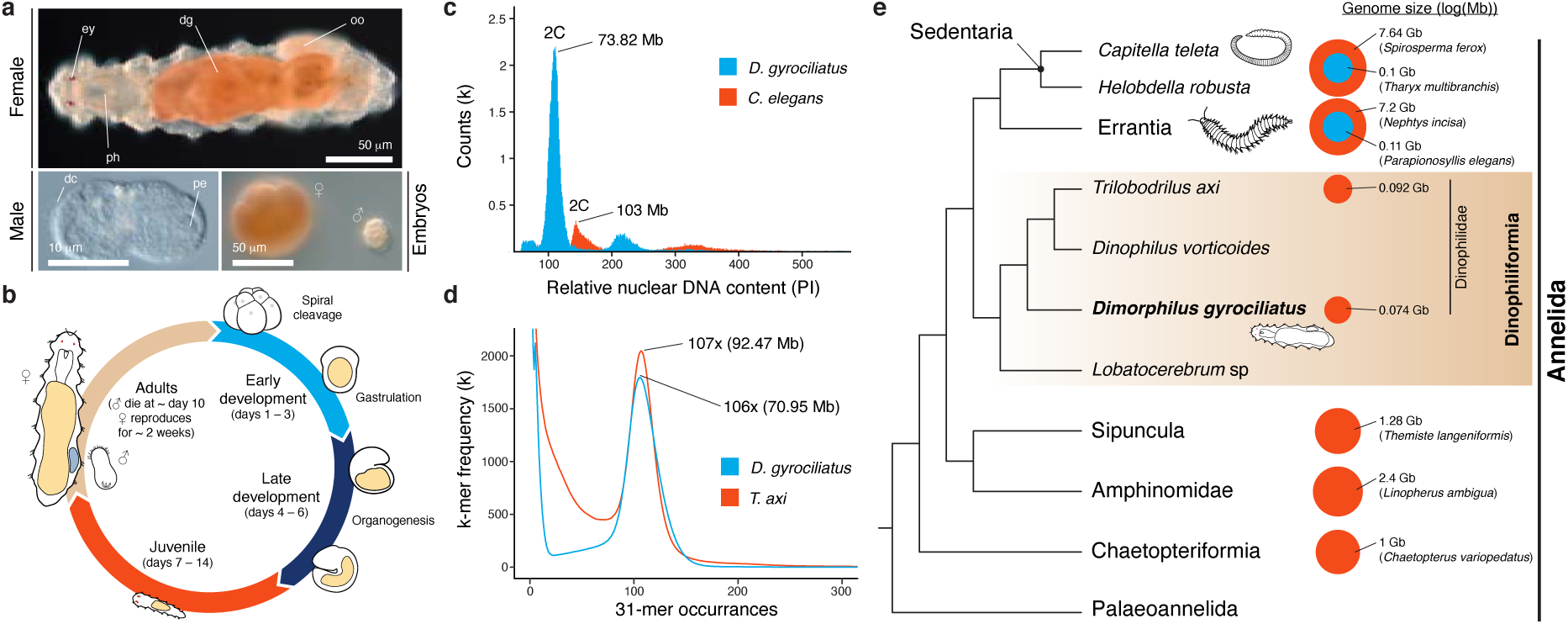
*Dimorphilus gyrociliatus* has the smallest annelid genome. (a) Differential interference contrast images of adults and embryos of *D. gyrociliatus*. The adults are miniature annelid worms with an extreme sexual dimorphism, already apparent during early embryogenesis. (b) The life cycle of *D. gyrociliatus* comprises a 6-days long embryogenesis with a canonical early spiral cleavage programme, followed by a juvenile and an adult, reproductively active stage. (c) Flow cytometry analysis using the nematode *C. elegans* as reference and propidium iodide (PI) nuclear intensity estimates the genome size of *D. gyrociliatus* in 73.82 Mb. (d) k-mer counts estimation the genome size of *D. gyrociliatus* and *T. axi* to be 70.95 Mb and 92.47 Mb, respectively. (e) *D. gyrociliatus* and *T. axi* belong to Dinophiliformia, the sister group to Sedentaria and Errantia, and their genome sizes are the smallest known among annelids. dc, dorsal ciliary field; dg, digestive system; ey, eye; oo, oocyte; pe, penis; ph, pharynx. Drawings are not to scale.

## Results

We performed long-read PacBio sequencing (Extended Data Fig. 1a) to generate a highly contiguous (N50, 2.24 Mb) and complete (95.8% BUSCO genes) ~78 Mb-long haploid assembly, comparable in quality to other published annelid genomes (Extended Data Fig. 1d, e; Supp. Table 1). Flow cytometry measurements and k-mer based analyses estimated the size of *D. gyrociliatus* genome to be 73.82 Mb and 70.95 Mb, respectively (Fig. 1c, d), agreeing with previous estimations^41^. While their simple morphology originally prompted to consider them early-branching annelids^51^ (“Archiannelida”), molecular phylogenies later placed *D. gyrociliatus* either within Sedentaria^52^ or as sister to Errantia+Sedentaria^53^, the two major annelid clades (Supp. Note 2). Gathering an extensive dataset of annelid sequences^54^, we robustly placed *D. gyrociliatus* together with *Trilobodrilus axi, Dinophilus vorticoides* and *Lobatocerebrum* sp. – all miniature annelids – in a clade we name Dinophiliformia that is sister to Errantia+Sedentaria, thus confirming previous hypotheses^53^ (Fig. 1e; Extended Data Fig. 2). Given the generally larger bodies and genome sizes found in annelid lineages outside Dinophiliformia (Fig. 1e), and that *T. axi* also has a compact, 92.47 Mb genome (Fig. 1d), our data suggest genome size reduction and morphological miniaturisation both occurred in the lineage leading to *D. gyrociliatus* and its relatives.

To assess how changes in repeat content contributed to genome reduction in *D. gyrociliatus*, we annotated the complement of transposable elements (TEs), uncovering a much lower percentage (4.87%) than in other annelid genomes (Fig. 2a; Extended Data Fig. 3a, b). Most TEs (91.5%) group in four classes and as in the annelid *Helobdella*^50^, TEs are either old copies or very recent expansions (Fig. 2b). The most abundant TE class is a Ty3-*gypsy*-like LTR retrotransposon that appears to be an annelid-or *D. gyrociliatus*-specific subfamily, and thus we name Dingle (<u>Dinophilidae</u> *<u>G</u>ypsy*-like <u>e</u>lements) (Extended Data Fig. 3c). As in some insect and nematode clades^55^, where LTR retrotransposon *envelope* (*env*) proteins are apparently related to *env* proteins of DNA viruses, Dingle *envelope* (*env*) protein shows similarities with envelope glycoprotein B precursors of cytomegalovirus (CMV) and herpesviridae-1 (HSV-1) (Extended Data Fig. 3d, e). Compared to species with minimal genome sizes, *D. gyrociliatus* TE load is 3–4x lower than in the appendicularian *Oikopleura dioica* and the tardigrade *Ramazzottius varieornatus*, but around 4x larger than in insects with larger, but still compact genomes (~100 Mb) (Supp. Table 5). Therefore, TE depletion contributed to genome compaction in *D. gyrociliatus*, but this does not appear to be the main driving factor, since other small animal genomes show even lower fractions of TEs.

**Figure 2.**
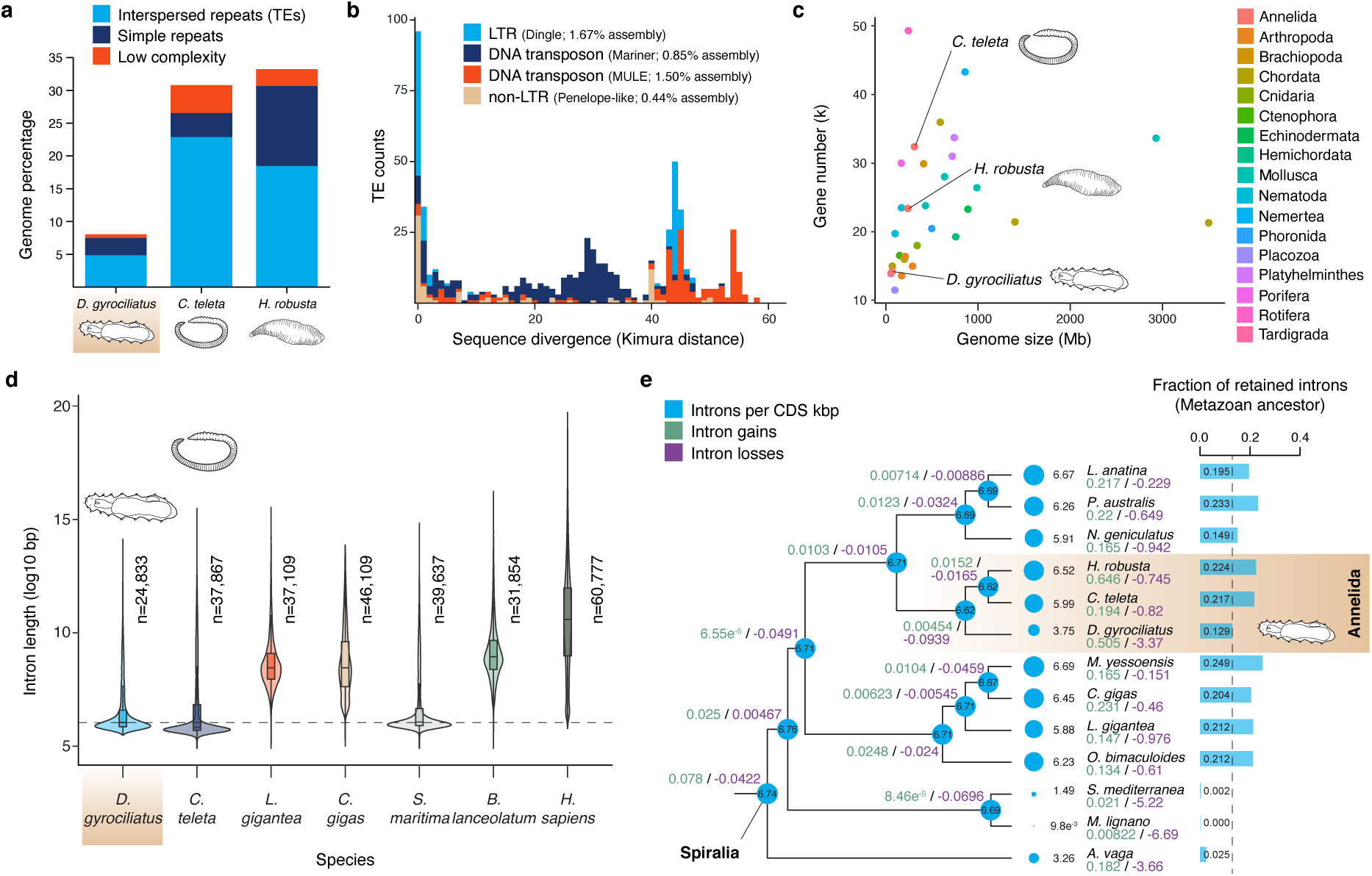
*Dimorphilus gyrociliatus* has a reduced transposable element and intronic landscape. (**a**) Percentage of the genome assigned to transposable elements (TEs) and repeats in three annelid genomes. *D. gyrociliatus* has considerably less TEs and simple repeats than other annelids. (**b**) TE abundance according to sequence divergence (Kimura distance) to family consensus. TE expansions are limited in size and correspond to either very recent bursts or old elements. (**c**) Number of annotated genes in 28 animal genomes plotted against genome size. *D. gyrociliatus* has a reduced gene repertoire compared to other annelids, but comparable to other animals of similar genome size. (**d**) Size distribution of orthologous introns in seven bilaterian species. Intron size is comparable between *D. gyrociliatus* and the annelid *C. teleta* and the centipede *S. maritima*, which are both slow evolving lineages with larger genomes. Dotted horizontal line indicates *D. gyrociliatus* median intron size. (**e**) Rates of intron gain (green), intron loss (red) and introns per kilobase of CDS (blue) in representative spiralian lineages and a consensus phylogeny. *D. gyrociliatus* has lost introns, yet at a much lower rate and preserving many more ancestral animal introns than other fast evolving spiralian lineages, such as flatworms and rotifers.

To explore how changes in gene architecture influenced genome compaction, we employed transcriptomic data and *ab initio* predictions to annotate 14,203 protein coding genes in the *D. gyrociliatus* genome, a smaller gene repertoire than that of other annelids (Fig. 2c; Extended Data Fig. 1b, c; Supp. Table 1). However, the gene number is comparable to free-living species with similar genome sizes, such as *O. dioica*^33^ (~15,000 genes) and *R. varieornatus*^32^ (~14,000 genes). With a gene density (208.86 genes/Mb) double than in the annelids *Capitella teleta* (99.96 genes/Mb) and *Helobdella robusta* (97.5 genes/Mb), *D. gyrociliatus* has shorter intergenic regions and transcripts, but similar exon lengths and even larger UTRs (Extended Data Fig. 4a, b, d–f), suggesting that intron shortening might have contributed to genome compaction. However, although *D. gyrociliatus* shows overall very short introns (median 66 bp) and its splicing is thus more efficient at removing short intron sizes (Extended Data Fig. 4i), introns are not shorter on average than in *C. teleta* (median 57 bp) and even similar to the centipede *Strigamia maritima* (median 67 bp) (Fig. 2d; Extended Data Fig. 4h), both with larger genomes than *D. gyrociliatus*. Instead, *D. gyrociliatus* has fewer introns than other annelids (Fig. 2e) and exhibits an intron density and ancestral intron retention comparable to other animals with small genome sizes, such as *O. dioica, C. elegans*, and *R. varieornatus* (Extended Data Fig. 4j, k). Therefore, gene and intron loss, rather than short intron size – which was likely a pre-existing condition – correlates with genome compaction in *D. gyrociliatus*, unlike in free living nematodes of similar genome size^56^.

To investigate how gene loss shaped the *D. gyrociliatus* genome and morphology, we first reconstructed clusters of orthologous genes employing a dataset of 28 non-redundant proteomes covering major animal groups and estimated gene loss and gain rates. Over 80% of *D. gyrociliatus* genes are assigned to multi-species gene families; the highest percentage in any annelid sequenced so far (Extended Data Fig. 5a). However, 38.9% of the genes in *D. gyrociliatus* are in orthogroups where there is only one *D. gyrociliatus* sequence, and thus *D. gyrociliatus* has the smallest average gene family size among annelids (1.63 genes/orthogroup; Supp. Table 7). Although the rate of gene family loss is greater than in *C. teleta*, an annelid species with a conservatively evolving genome^50^, gene loss in *D. gyrociliatus* is similar to those of the annelids *H. robusta* and *Hydroides elegans*, species with larger genomes (Fig. 3a; Extended Data Fig. 5b). Therefore, our data suggest that reduction of gene family size outweighs complete gene family loss, and thus likely underpins the reduced total gene number of *D. gyrociliatus*, as also observed in certain *Caenorhabditis* species of small genome size^56,57^ Consistent with the streamlining of its gene repertoire, we detected only nine expanded gene families in *D. gyrociliatus* (but 73 and 42 in *C. teleta* and *H. robusta*, respectively), most of them corresponding to locally duplicated genes implicated in immune responses (Extended Data Fig. 5c–e). In addition, *D. gyrociliatus* shows canonical repertoires of gene families expanded in other annelids, such as G protein-coupled receptors (GPCRs) and epithelial sodium channels (ENaCs)^50^ (Extended Data Fig. 6a, b; Supp. Table 8). The GPCR complement of genomes is dynamic and often linked to specific (neuro)physiological adaptations, as seen in lineages with miniature genomes that have experienced either losses (e.g. *O. dioica* lacks Class C, glutamate receptors) or expansions (e.g. *C. elegans*^58^ and *R. varieornatus*^59^ expanded Class A, rhodopsin receptors) (Extended Data Fig. 6b). Thus, the conserved GPCR repertoire and the canonical neuropeptide complement (Extended Data Fig. 6c) further support that *D. gyrociliatus* nervous system is functionally equivalent to, though morphologically smaller than that of larger annelids^47,48^.

**Figure 3.**
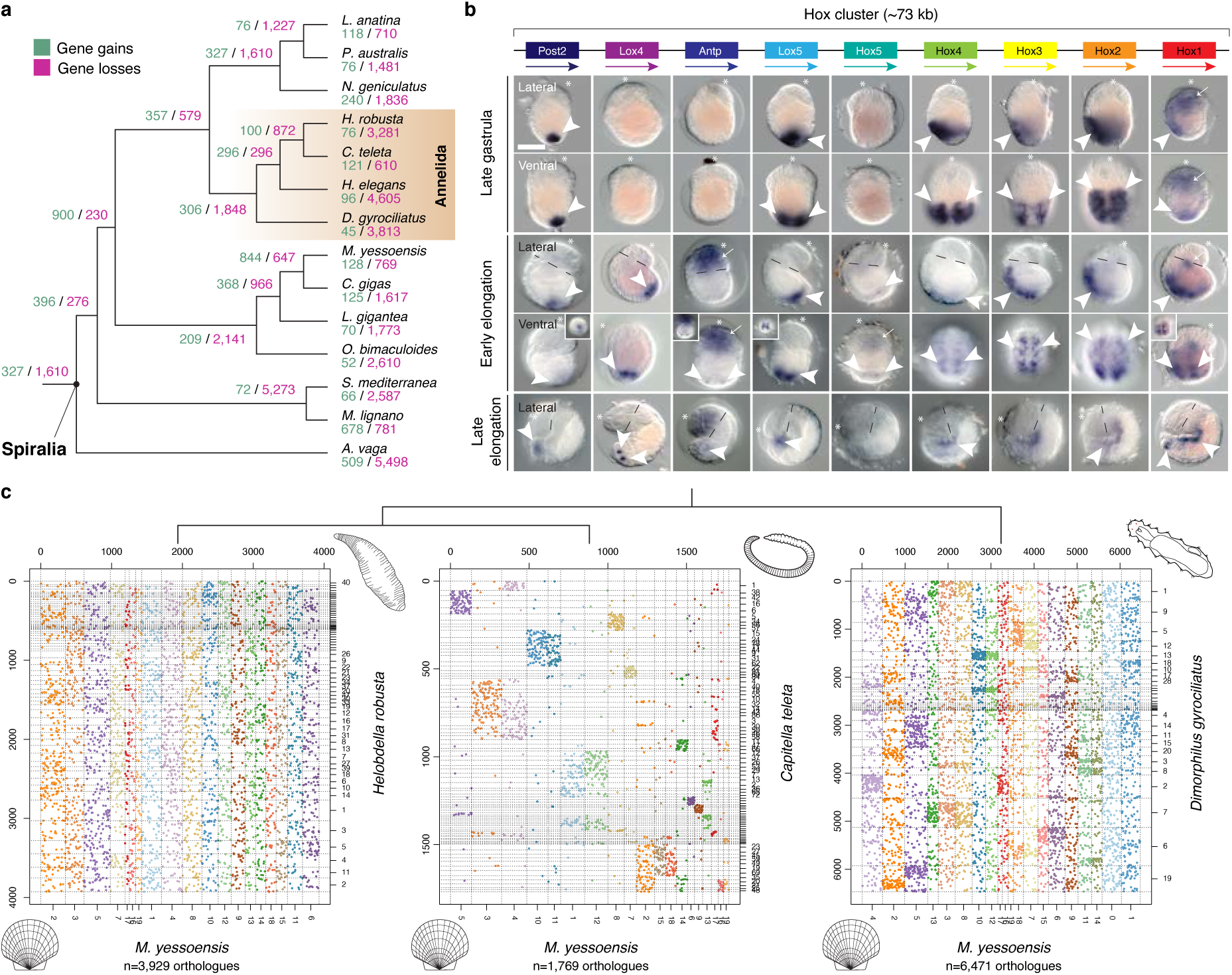
*Dimorphilus gyrociliatus* has retained a conserved developmental toolkit and ancestral linkeage blocks. (**a**) Number of gene family gains (green) and losses (red) in representative spiralian lineages under a consensus tree topology. Gene loss in *D. gyrociliatus* is similar or lower than that observed in other fast evolving spiralian lineages. (**b**) *D. gyrociliatus* has a conserved Hox complement, organised in a compact cluster (top). Whole mount *in situ* hybridisation during embryogenesis reveals that Hox genes exhibit staggered anteroposterior domains of expression, but not temporal collinear expression domains (arrowheads) along the trunk region, with *Hox1, Hox5* and *Antp* further exhibiting anterior head expression domains (arrows). Dotted lines in lateral views of early and late elongation timepoints demarcate the head-trunk boundary. Scale bar, 50 µm. (**c**) Oxford dot plots of orthologous genes between the scallop *M. yessoensis* and three annelid genomes. Orthologous genes are coloured according to their position in *M. yessoensis* linkage groups. The presence of an organised Hox cluster correlates with the preservation of some macrosyntenic blocks (areas of higher density of shared orthologs) in *D. gyrociliatus*, which are lost in the fast-evolving *H. robusta*.

Despite its miniature body plan, *D. gyrociliatus* has an overall conserved developmental toolkit, at the level of both transcription factors and signalling pathways (Extended Data Fig. 5f, g). *D. gyrociliatus*, and Dinophilidae generally, exhibit a limited repertoire of certain extracellular signalling molecules (e.g. Wnt and TGF-b ligands) and lacks *bona fide* FGF and VEGF ligands (Extended Data Fig. 5g–i). However, these simplifications do not affect the receptor repertoire (Extended Data Fig. 5j). Unlike appendicularians^60^, tardigrades^32^ and nematodes^32^ with compact genomes, *D. gyrociliatus* exhibits a compact, ordered Hox cluster, only lacking *post1* (Fig. 3b; Extended Data Fig. 7a, b). In other annelids^61,62^, *post1* is separate from the main Hox cluster, and as in brachiopods^63^, it is expressed in chaetoblasts^62^, supporting the homology of this novel cell-type^63^. Remarkably, the distantly related *H. robusta* and *D. gyrociliatus* both lack chaetae, *post1*, and FGF ligand (also expressed in annelid chaetoblasts; Extended Data Fig. 5k–r), suggesting that the secondary loss of chaetae followed convergent routes of gene loss in different annelid species.

To investigate whether the clustered Hox genes of *D. gyrociliatus* exhibit temporal collinearity, we first performed comparative transcriptomics at four different stages of the *D. gyrociliatus* female life cycle (Extended Data Fig. 8a, b). Genome-wide expression dynamics revealed five main clusters of co-regulated genes (Extended Data Fig. 8c), corresponding to major developmental events, such as cell proliferation in early development or during adult growth (clusters 5 and 4, respectively), sex differentiation (cluster 2), nervous system maturation during late embryogenesis and post-embryogenesis (cluster 1) and increased metabolism after hatching (cluster 3). While there is a gradual increase in gene upregulation as embryogenesis proceeds, which stabilises in the juvenile to adult transition (Extended Data Fig. 8d–f), all Hox genes but *Hox5, Antp* and *post2* are expressed during early embryogenesis (days 1–3; Extended Data Fig. 7c). Employing whole mount *in situ* hybridisation, we identified late gastrula (~3 days after egg deposition) as the earliest stage at which most Hox genes become simultaneously transcribed, including *post2* (Fig. 3b), altogether suggesting that *D. gyrociliatus* Hox genes lack temporal collinearity. Different from other annelid species^64–66^, *D. gyrociliatus* embryogenesis is slow, taking about six to seven days from egg laying to hatching (Fig. 1b), and thus it is unlikely that Hox temporal collinearity is compressed to span a short and quick early morphogenesis. During body elongation and segment formation, Hox genes are expressed in staggered anteroposterior domains along the developing trunk, in patterns resembling those of *C. teleta*^62^, further supporting that *D. gyrociliatus* retains the ancestral annelid molecular body patterning (Fig. 3b; Extended Data Fig. 7d). Therefore, *D. gyrociliatus* Hox genes show only staggered expression domains along the anteroposterior axis (Extended Data Fig. 7e), providing a compelling case where temporal collinearity is not driving Hox cluster compaction and maintenance^67^.

Animal groups with reduced genome sizes show altered gene orders, as exemplified by their disorganised Hox clusters^60,68^ and the loss of conserved gene linkage blocks that represent the ancestral chromosomal organisation^36,50^. In *O. dioica*, this loss has been related to the loss of the classical Non-Homologous End Joining (c-NHEJ) double strand DNA break (DSB) repair pathway^69^. In addition to an ordered Hox cluster, *D. gyrociliatus* shows residual conservation of ancestral linkage blocks, which appear eroded but still visible (Fig 3c). These blocks are almost intact in *C. teleta* but completely lost in *H. robusta* (Fig. 3c; Extended Data Fig. 7f). Moreover, *D. gyrociliatus* has a conserved DSB repertoire (Supp. Table 9), with the exception of BRCA1, which is however also absent in other invertebrates capable of homologous recombination, such as *Drosophila melanogaster*^70^. Therefore, mutation prone DSB repair mechanisms that can increase DNA loss do not underpin genomic compaction in *D. gyrociliatus*, which occurred without drastic genome architecture rearrangements.

Changes in genome size have been positively correlated to differences in cell and body size in a range of animal groups^1,21–24^. Given the miniature body size and the compact genome of *D. gyrociliatus*, we thus hypothesised that the molecular mechanisms controlling cell and organ growth might exhibit critical divergences in this lineage, should these two traits be connected. To test this hypothesis, we used genome wide KEGG annotation (Supp. File 4) to reconstruct signalling pathways known to be involved in the control of cell growth and proliferation (cyclin/CDKs^71^, PI3K/Akt/mTOR^72^) and organ size (Hippo pathway^73^) in metazoans (Fig. 4a). *D. gyrociliatus* shows orthologs of all core components of these pathways (Supp. Table 10), with the exception of PRR5 – an mTOR complex 2 interactor that is however dispensable for complex integrity and/or kinase activity^74^ – and a clear ortholog of p21/p27/p57 kinases, general inhibitors of cyclin-CDK complexes among other roles^75^. Besides, the Myc transduction pathway, which regulates growth and proliferation^76^ and sits downstream of the Hippo and PI3K/Akt/mTOR pathways^73,77^, lacks the regulators *mad* (in *D. gyrociliatus*) and *mnt* (in all Dinophilidae), a condition also shared with the appendicularian *O. dioica* (Fig. 4b; Supp. Table 11). In Dinophilidae, MYC additionally has a W135 point mutation in the broadly conserved MYC box II (MBII) transactivation domain that has been shown to impair MYC function in human cells, in particular its ability to repress growth arrest genes^78^ (Fig. 4c). Myc downregulation in vertebrates and flies causes hypoplasia^79^, which could explain the miniature size of dinophilids, and slows down DNA replication^80^, which could act as a selective pressure favouring smaller genomes. Although the full extent of these genomic changes is hard to evaluate given the poor understanding of cell and organ growth in annelids, our data provide a substrate for studying whether there is a mechanistic link between genome size reduction and organism miniaturisation in *D. gyrociliatus*.

**Figure 4.**
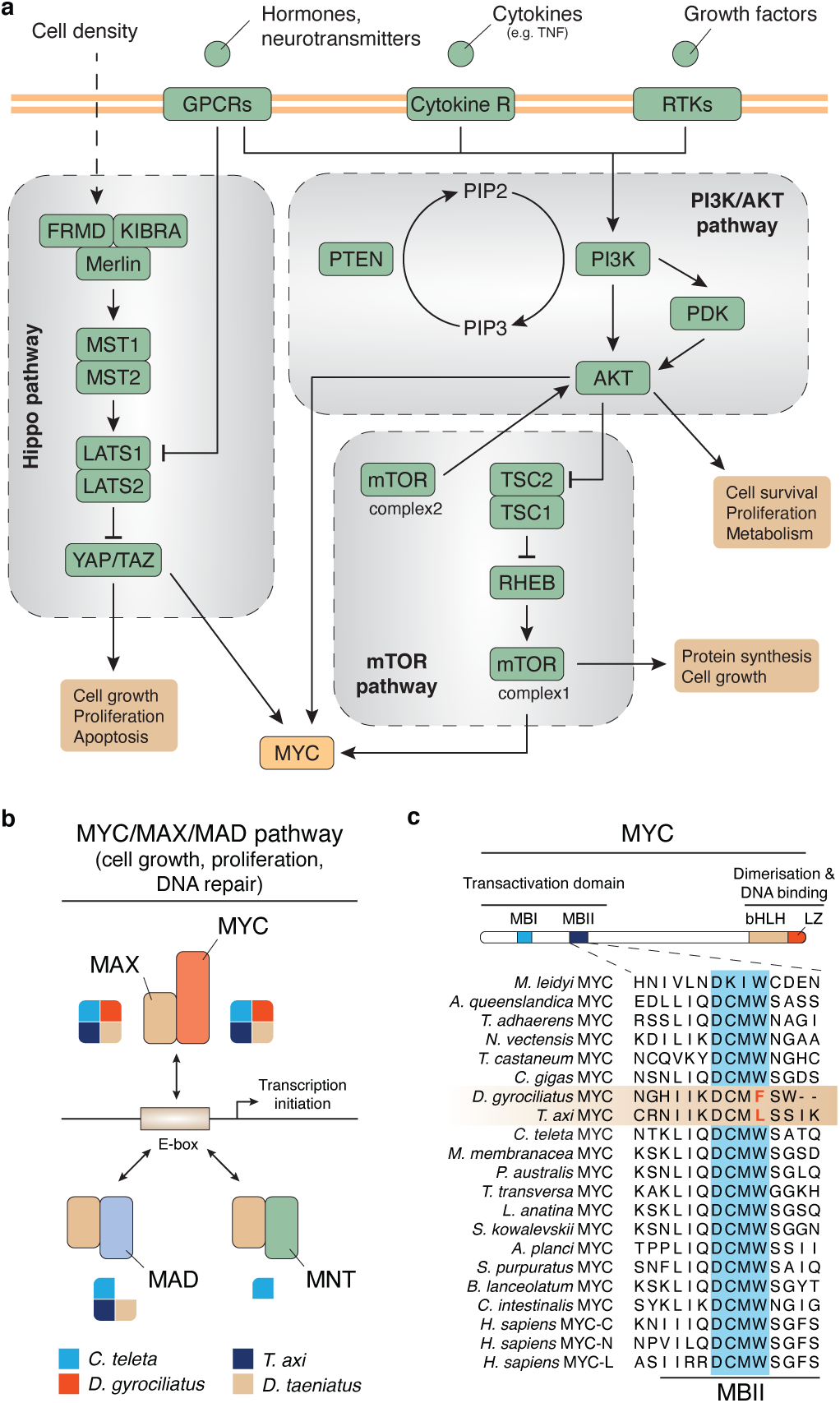
*Dimorphilus gyrociliatus* exhibits a divergent MYC pathway. (**a**) Schematic representation of signalling pathways involved in cell growth/proliferation and organ size in animals. *D. gyrociliatus* shows conserved Hippo and PI3K/Akt/mTOR pathways (green boxes), but also divergences in the MYC pathway (orange box), one of the downstream regulators. See main text and Supplementary Table 11 for a complete list of genes. (**b**) Schematic representation of the MYC/MAX/MAD pathway and the interactions between the main protein partners. *D. gyrociliatus* lacks *bona fide* MAD and MNT proteins (the latter also absent in other members of Dinophilidae). (**e**) Multiple protein alignment of the MBII repressor domain of MYC, highlighting how Dinophilidae exhibit point mutations in the critical tryptophan (W) residue.

To investigate how compaction affected genome regulation, we first used ATAC-seq to identify ~10,000 reproducible open chromatin regions in adult *D. gyrociliatus* females (Extended Data Fig. 9a–d). Open chromatin regions are short in *D. gyrociliatus* and mostly found in promoters (Fig. 5a, b), consistent with its small genome size and small intergenic regions. Despite the generally short intron size in *D. gyrociliatus*, 944 ATAC-seq peaks were in intronic regions significantly larger than non-regulatory introns (Fig. 5c). We recovered a canonical regulatory profile (Fig. 5d), which together with the lack of putative spliced leaders in 5’ UTRs (Extended Data Fig. 4g), suggests that *trans*-splicing and operons do not occur in *D. gyrociliatus*, similar to other annelids^81^. The CTCF DNA-binding motif was the most abundant in active regulatory regions, located mostly in promoters and as single motifs (Fig. 5e; Extended Data Fig. 9e–h). Unlike nematodes with compact genomes^82^, which lack CTCF, the *D. gyrociliatus* genome encodes for a CTCF ortholog (Supp. Fig. 8). However, localisation of CTCF DNA-binding motifs, for the most part close to transcriptional start sites, instead of in intergenic regions, suggests that CTCF might play a role in regulating gene expression in *D. gyrociliatus* rather than in chromatin architecture as seen in vertebrates^83^. Thus, our data indicate that *D. gyrociliatus* has retained conserved genomic regulatory features (e.g. lack of operons and trans-splicing, presence of CTCF) but streamlined regulatory regions and potentially lost distal intergenic *cis*-regulatory elements with genome compaction.

**Figure 5.**
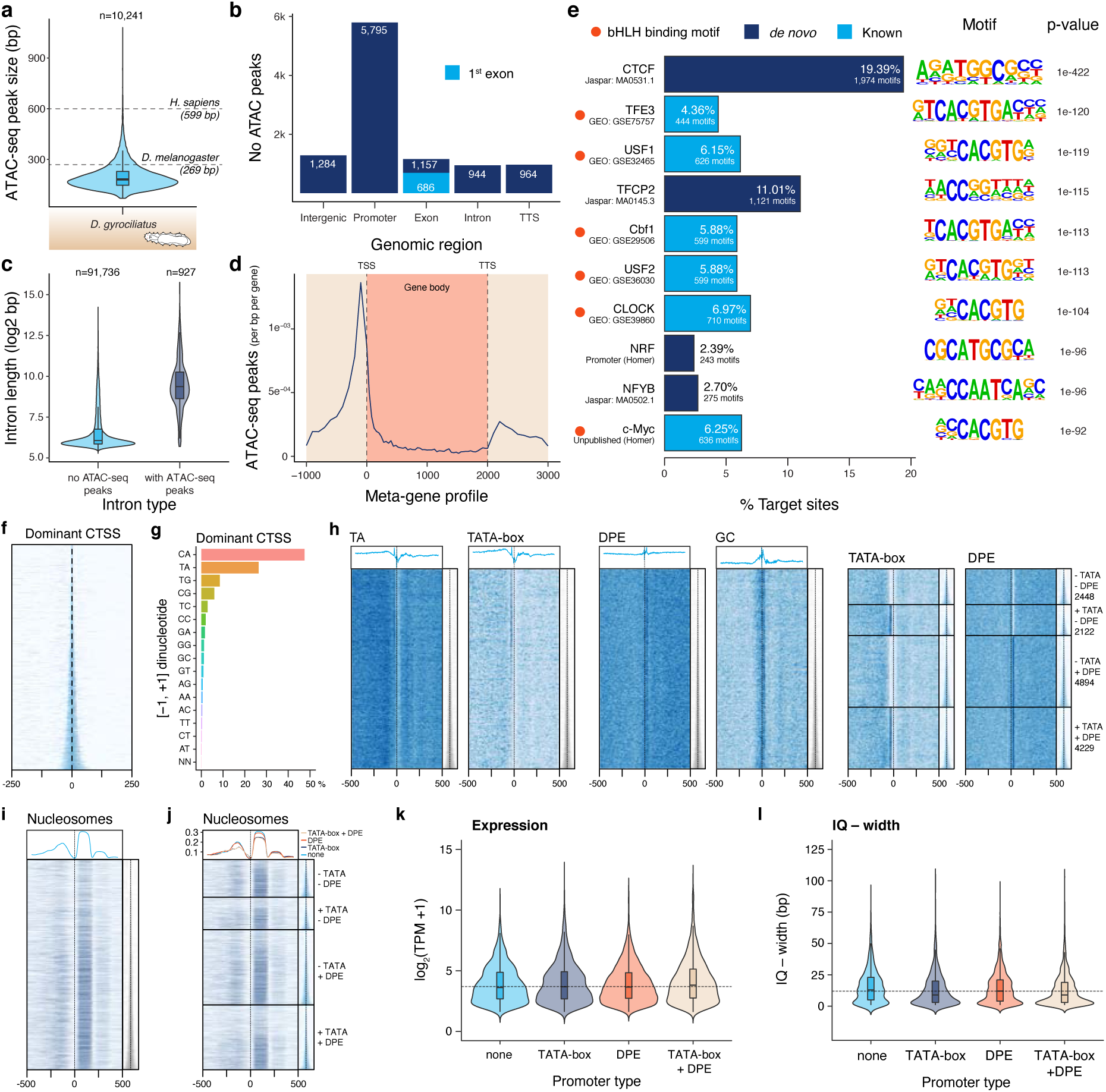
The regulatory genomic landscape of *Dimorphilus gyrociliatus*. (**a**) Violin plot depicting ATAC-seq peak size distribution in *D. gyrociliatus* compared to the median values in the fly *D. melanogaster* and humans. The open chromatin regions are shorter in *D. gyrociliatus* than in other animal genomes. (**b**) Distribution of ATAC-seq peaks according to genomic feature. Most of the open chromatin regions are found in promoters, intergenic regions and (first) introns. (**c**) Violin plots of size distributions in introns with and without ATAC-seq peaks. The presence/absence of open chromatin regions in introns correlates positively with size. (**d**) Metagene profile of ATAC-seq signal. All gene lengths are adjusted to 2kb. (**e**) Top ten most-significant motifs identified in *D. gyrociliatus* ATAC-seq peaks. The most abundant motif in open chromatin regions corresponds to CTCF. (**f**) Tag-clusters centred on the dominant CAGE-supported TSS (CTSS) are usually narrow (based on interquantile range q0.1-q0.9) and (**g**) retain the canonical metazoan polymerase II initiation pyrimidine (C, T)/purine (A, G) dinucleotides. (**h**) Most (11,245 out of 13,693) of the CTSS have a TATA-box and/or a downstream promoter element (DPE). (**i**) Nucleosomes are consistently located after the CTSS, regardless of the promoter type (**j**). (**k, l**) While genes with a TATA-box tend to be slightly narrower on average, there are not major differences in expression levels between genes with different promoter elements.

Since most regulatory information is restricted to promoter regions (<1 kb upstream the transcription start site, TSS), we applied CAGE-seq to characterise promoter architecture (Extended Data Fig. 10a). Promoters are narrow (<150 bp) in *D. gyrociliatus* and use pyrimidine-purine dinucleotides as preferred initiators (Fig. 5f, g; Extended Data Fig. 10e). Upstream TA and downstream GC enrichment, respectively, revealed the presence of TATA-box and downstream promoter elements (DPE) in *D. gyrociliatus*, with TATA-box generally associated with short promoters (Fig. 5h; Extended Data Fig. 10f). Similar to vertebrates^84^, strength of nucleosome positioning correlates with promoter broadness in *D. gyrociliatus* (Fig. 5i) and thus narrow TATA-box dependent promoters have lower +1 nucleosome occupancy than wide non-TATA-box promoters (Fig. 5j). As in other eukaryotes, TATA-box containing *D. gyrociliatus* promoters have somewhat higher expression levels, while promoters with DPE motif have no particular features, indicating this element might be non-functional (Fig. 5k, l). Therefore, the general *D. gyrociliatus* promoter architecture resembles that of other bilaterians (Extended Data Fig. 10g), further supporting that genomic compaction did not alter genome regulation.

## Discussion

Our study demonstrates that genome compaction and morphological miniaturisation are specificities of *D. gyrociliatus* (Fig. 1e), grounded in a nested phylogenetic position within Annelida, TE depletion, intergenic region shortening, intron loss and streamlining of the gene complement and genome regulatory landscape (Fig. 2a, e; Fig. 3a; Fig. 5a, f). Traditionally, morphological miniaturisation in *D. gyrociliatus* and Dinophiliformia has been considered a case of progenesis (underdevelopment)^45,52^, yet the exact underlying mechanisms are unknown. As in other animal lineages^34,35,85^, our data support that morphological change might be partially explained by gene loss in *D. gyrociliatus* (Fig. 6a), as we identified a reduced repertoire of extracellular signalling ligands and the loss of developmental genes related to missing organs, such as chaetae (*post1* and FGF ligand) and mesodermal derivatives like coeloms (VEGF ligand). However, *cis*-regulation of gene expression is mostly restricted to the proximal regions in *Dimorphilus* (Fig. 5b). Therefore, our study suggests that coordinated distal gene regulation, which is an animal innovation^86^ whose emergence has been associated with the evolution of sophisticated gene regulatory landscapes and morphological diversification^87,88^, is also limited in *D. gyrociliatus*.

**Figure 6.**
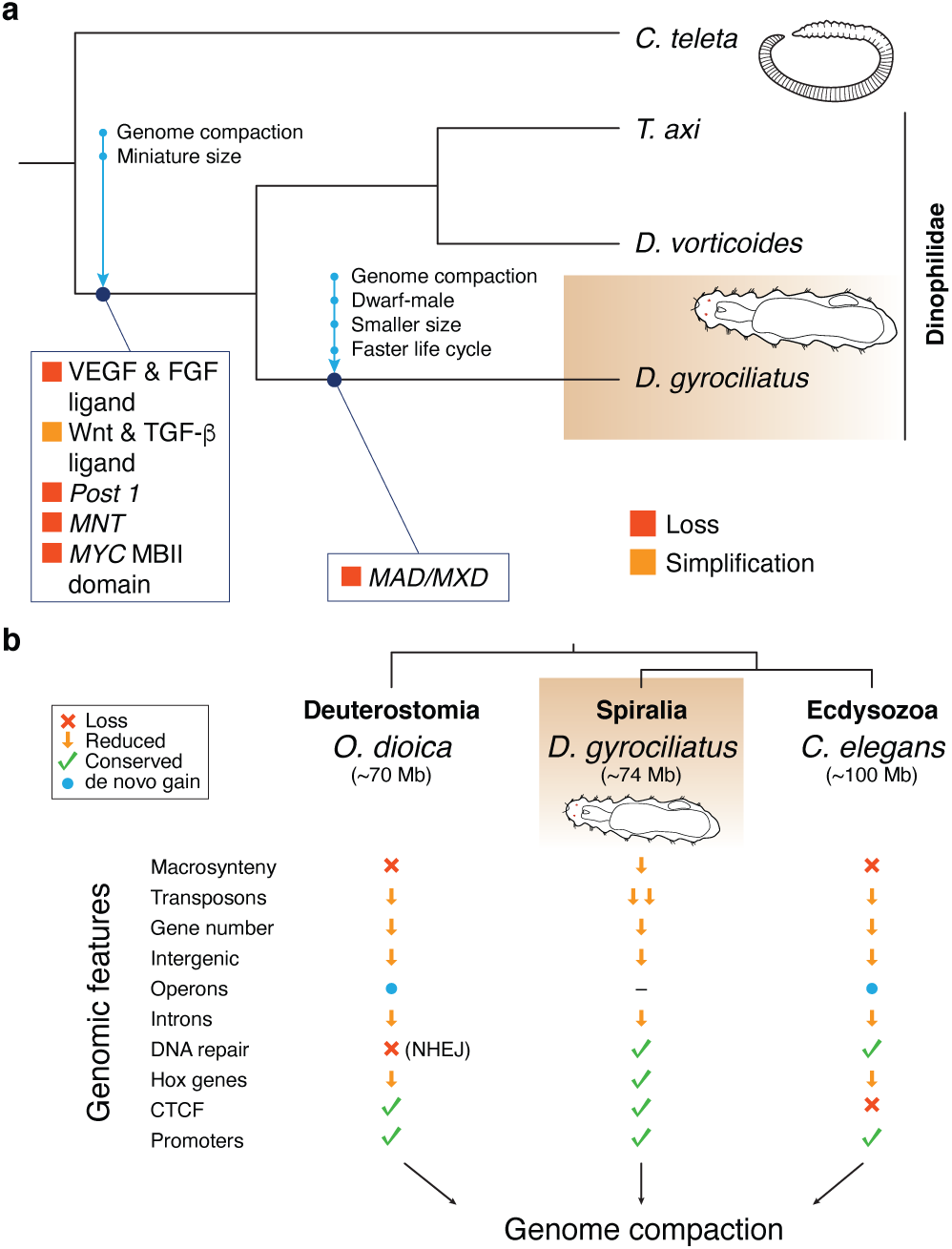
A novel conservative route to genome compaction in *Dimorphilus gyrociliatus*. (**a**) Schematic diagram of the genomic changes which occurred during genome compaction and morphological miniaturisation in *D. gyrociliatus* and Dinophilidae. (**b**) *D. gyrociliatus* genome represents a more conservative evolutionary pathway to genome compaction compared to the more drastic genomic changes experienced by other bilaterian lineages with compact genomes, such as *O. dioica* and *C. elegans*.

Unlike in other cases of genomic compaction^5,30–33,36–39^, but similar to what has been reported for the teleost fish *Takifugu rubripes*^89,90^, our work provides compelling evidence that genome miniaturisation did not trigger drastic changes in genome architecture and regulation in *D. gyrociliatus* (Fig. 3c; Fig. 5c, e, h; Fig. 6b). Therefore, the genomic features observed in appendicularians, tardigrades and some nematodes are lineage specificities that might have eventually facilitated genome compaction, but that are not always associated with genome size reduction, thus questioning the assumed causal link between fast evolving genomic traits and genome compaction. Altogether, our study characterises an alternative, more conservative route to genome compaction, and furthermore provides an exciting new system and genomic resources to investigate the evolutionary plasticity and function of core cellular mechanisms in animals.

## Methods

### Genome sequencing and assembly

Adult females of *D. gyrociliatus* were used to isolate genomic DNA (gDNA) following standard guanidium isothiocyanate protocol and RNase A treatment. Library was prepared using Pacific Biosciences 20 kb library preparation protocol and size-selected using BluePippin with 5 kb cut-off. The library was sequenced on a Pacific Bioscience RS II instrument using P6-C4 chemistry at the Norwegian Sequencing Centre (NSC). An Illumina library of median insert size of 298 bp was sequenced in 101 bases paired end mode on an Illumina HiSeq 2500 instrument at GeneCore (EMBL). All raw sequence data associated to this project are available under the project with primary accession PRJEB37657 in the European Nucleotide Archive (ENA).

PacBio reads were filtered with SMRTAnalysis v.2.3.0.140936 and assembled with PBcR v.8.3rc2^91,92^ using default options, except for k-mer=14 and asmMerSize=14. Four rounds of decontamination using Blobtools v.0.9.16^93^ were applied, removing contigs with similarity to bacteria, algae, fungi or unicellular eukaryotes. A consensus assembly was generated with Quiver and improved with Pilon v.1.16^94^ using the Illumina paired end reads previously filtered for adapters with cutadapt v.1.4.2^95^. We used HaploMerger2 v.20151124^96,97^ to reconstruct a high-quality haploid reference assembly, which we further scaffolded with SSPACE-LongRead v.1.1^98^.

We collected hundreds of adult individuals of *Trilobodrilus axi* Remane, 1925 at the intertidal beach of Königshafen, Sylt (Germany)^44^ and extracted gDNA as described above to prepare a TruSeq v3 Illumina library that was sequenced in 101 bases paired end mode on a full lane of an Illumina HiSeq 2500 instrument at GeneCore (EMBL). Before assembly, we removed adapters and low quality regions with cutadapt v.1.4.2^95^ and Trimmomatic v.0.35^99^, error correction with SPAdes v.3.6.2^100^ and deduplication with Super_Deduper v.2.0. Cleaned reads were assembled with Platanus v.1.2.4^101^ and contigs with similarity to proteobacteria were identified with Blobtools v.0.9.16^93^. After removal of bacterial contigs, we generated the final assembly with Velvet v.1.2.10^102^.

We used BUSCO v2 pipeline^103^ to validate the completeness of the genome assemblies. Out of the 978 metazoan BUSCO genes, 930 were complete (95.1%), 7 were fragmented (0.7%) and 41 were missing (4.2%) (Extended Data Figure 1e) in the *D. gyrociliatus* genome assembly. Only 27 (2.8%) of the BUSCO genes were complete and duplicated. BUSCO analysis on the *T. axi* genome resulted in 835 complete (85.4%), 27 complete and duplicated (2.8%), 75 fragmented (7.7%) and 68 missing (6.9%) (Extended Data Figure 1e). Finally, we used KAT v.2.4.2^104^ to estimate the completeness and copy number variation of the assemblies (Supplementary Figure 1).

### Genome size measurements

For flow cytometry measures, adult *D. gyrociliatus* females and *C. elegans* worms (reference) were starved for 3–4 days before analysis. *D. gyrociliatus* and *C. elegans* were chopped with a razor blade in General-Purpose Buffer^105^ and the resulting suspension of nuclei was filtered through a 30 µm nylon mesh and stained with propidium iodide (Sigma; 1 mg/mL) on ice. We used a flow cytometer Partex CyFlow Space fitted with a Cobalt Samba green laser (532 nm, 100 mW) to analyse the samples, performing three independent runs with at least 5,000 nuclei per run. For k-mer-based measures, we used the raw Illumina paired-end reads of *D. gyrociliatus* and *T. axi*. We removed adapters using cutadapt^95^ v.1.4.271, quality trimmed the reads using Trimmomatic^99^ v.0.3575, performed error-correction using SPAdes^100^ v.3.6.276 and removed duplicated reads using Super-Deduper v.2.0. We identified and removed contaminant reads using BlobTools v.1.1.1, and normalised read coverage to 100x in both datasets using BBNorm from BBTools suite v38.86 to mitigate the effects of a strong GC content bias in *D. gyrociliatus* and reduce the impact of highly abundant repeats in *T. axi*. We used Jellyfish v.2.2.386^106^ to count and generate a histogram of canonical 31-mers, and GenomeScope 2.0^107,108^ to estimate the genome size and heterozygosity (Fig. 1d, Supplementary Figure 2). We also used Smudgeplot^107^ to estimate ploidy and analyse the genome structure (Supplementary Figure 3).

### Transcriptome sequencing and assembly

A publicly available dataset (Sequence Read Archive, accession number SRX2030658) was used to generate a *de novo* transcriptome assembly as previously described^47^. Redundant contigs were removed using *cd-hit-est* program with default parameters of CD-HIT^109^ and CAP3^110^. Additionally, we used that dataset to generate a genome-guided assembly using Bowtie2^111^ and Trinity v2.1.1^112^. Supplementary Table 2 shows standard statistics for the *de novo* and genome guided assemblies calculated with Transrate^113^. Transcriptome completeness was evaluated with BUSCO v2^103^.

### Stage-specific RNA-seq

Two biological replicates of four developmental stages of *D. gyrociliatus* (early embryo, 1–3 days old; late embryo, 4–6 days old; juvenile females, 7–9 days old; and adult females, 20–23 days old) were used to isolate total RNA with TRI Reagent® Solution (Applied Biosystems) following manufacturer’s recommendations and generate Illumina short-reads on a NextSeq 500 High platform in 75 base paired end reads mode and a ~270 bp library mean insert size at GeneCore (EMBL). We pseudo-aligned reads to *D. gyrociliatus* filtered gene models with Kallisto v0.44.0^114^, and followed the standard workflow of DESeq2^115^ to estimate counts, calculate size factors, estimate the data dispersion, and perform a gene-level differential expression analysis between consecutive stages (Supplementary Data Files 1–3). Datasets were first corrected for low count and high dispersion values using the *apeglm* log fold change shrinkage estimator^116^, and then compared using Wald tests between contrasts. For clustering and visualization, we homogenized the variance across expression ranks by applying a variance stabilizing transformation to the DESeq2 datasets. We used the *pheatmap* package to create heatmaps^117^, the package *EnhancedVolcano* for volcano plots^118^, and *ggplot2* for the remaining plots^119^. To characterize and identify enriched gene ontology terms, we used the package *clusterProfiler*^120^. All analyses performed in R^121^ using the RStudio Desktop^122^.

### Phylogenetic analysis

Annelid transcriptomes (Supplementary Data File 4) were downloaded from SRA and assembled using Trinity v2.5.1^112^ with the Trimmomatic^99^ read trimming option. Transcriptomes were then translated using Transdecoder v5.0.2^112^ after searching for similarity against the metazoan Swissprot database. Predicted proteins were searched using HMMER^123^ for 1,148 single-copy phylogenetic markers previously described^124^ using reciprocal BLAST to discard possible paralogues and character supermatrix was assembled as described before^124^. From this initial dataset, we selected the 264 genes with lowest saturation, yielding a concatenate alignment of 71,508 positions (as the analysis of the full dataset with site-heterogeneous models was not computationally tractable). Phylogenetic analyses were performed on the concatenated alignment using IQTREE^125^ with a C60 mixture model, a LG matrix to account for transition rates within each profile, the FreeRate heterogeneity model (R4) to describe across sites evolution rates and an optimisation of amino acid frequencies using maximum likelihood (FO). Support values were drawn from 1,000 ultrafast bootstraps with NNI optimisation. We also carried out Bayesian reconstruction using a site-heterogeneous CAT+GTR+Gamma model running two chains for more than a thousand iterations. We reached reasonable convergence for one of the datasets (bpdiff > 0.19).

### Annotation of repeats and transposable elements

We used RepeatModeler v.1.0.4^126^ and RepeatMasker “open-4.0”^127^ to generate an automated annotation of transposable elements (TEs) and repeats (Supplementary Table 3). We performed a BLAST analysis using the TE sequences recovered with RepeatModeler and PFAM sequence collections corresponding to entries RVT_1 (PF00078) (Supplementary Table 4) to uncover non-LTR retrotransposons and Helitrons represented by only a few copies. Using MITE Digger^128^, we identified MITEs whose Terminal Inverted Repeats (TIRs) matched *Mariner* transposons in the *D. gyrociliatus* genome. *D. gyrociliatus* DNA transposons belong to *Mariner* and *Mutator-Like Elements* (*MULE*) based on the amino-acid signature of their transposases. To establish the gene arrangement in LTR retrotransposons, we performed 6-frame translations of most intact copies, identified as such by having two identical LTRs and being flanked by short direct repeats created by Target Site Duplication (TSD). LTR retrotransposons were further compared to other elements of the *Ty3/gypsy* clade using a set of protein sequences comprising the reverse transcriptase domain and the integrase core domain. The phylogeny of *Ty3/gypsy* was established with a collection of sequences from the Gypsy database^129^, including hits obtained with TBLASTN (databases NR and TSA) using *D. gyrociliatus* sequences as queries. To look for distant homologues of the protein found downstream the integrase in LTR retrotransposons, we submitted a multiple sequence alignment of ten peptide sequences (corrected to the original coding frame when recovered from disrupted genes) to HHPred (database PDB_mmCIF70_28_Dec). Using MODELLER, the three best hits (Probability > 99, E-value < e^−29^) were used to model the 3D structure of the *Dingle-1* envelope.

### Gene prediction and functional annotation

The predicted set of core eukaryotic genes generated by CEGMA^130^ was used to train and run AUGUSTUS v.3.2.1^131^. The predicted proteomes of the annelids *Capitella teleta* and *Helobdella robusta* were aligned to the *D. gyrociliatus* genome using EXONERATE v.2.2.0^132^, and PASA v.2.0.2^133^ was used to align the transcriptome to the genome with BLAT and GMAP aligners^134,135^. EvidenceModeler v.1.1.1^136^ was used to generate weighted consensus gene predictions, giving a weight of 1 to *ab initio* gene predictions and spliced protein alignments, and a weight of 10 to the PASA transcript assemblies. EvidenceModeler output was used to refine PASA gene models and generate alternative splice variants. Predictions with BLAST hit against transposons and/or with an overlap equal or greater than 90% on masked regions were removed. The final prediction set contains 14,203 coding-protein loci that generate 17,409 different peptides. We used ORFik^137^ to refine transcription start sites (TSS) with Cap Analyses Gene Expression (CAGE)-seq data. Functional annotation for the 17,409 different transcripts was performed with Trinotate v.3.0. We retrieved a functional annotation for 13,437 gene models (77.18%).

### Gene structure evolution

We compared genome-wide values of gene structure parameters among *D. gyrociliatus, C. teleta, H. robusta, D. melanogaster, C. elegans* and *O. dioica*. For *C. teleta, H. robusta, D. melanogaster*, and *C. elegans* (Supplementary Table 6). To identify splice leader sequences in *D. gyrociliatus*, we predicted protein coding sequences in the *de novo* assembled transcriptome with Transdecoder v.5.5.0^112^ and used the scripts *nr_ORFs_gff3.pl* (from Transdecoder) and *gff3_file_UTR_seq_extractor.pl* (from PASA) to extract the non-redundant 5’ UTR sequences of protein coding transcripts. We used these sequences and Jellyfish v.2.2.3^106^ to identify over-represented 22-mer and 50-mer sequences that would correspond to the splice leader.

### Intron evolution analysis

We compared distributions of intron lengths between *D. gyrociliatus, Homo sapiens, Capitella teleta, Crassostrea gigas, Lottia gigantea, Strigamia maritima* and *Branchiostoma lanceolatum* (Supplementary Table 6) using only introns in genes with orthologs across the seven species (as defined by OrthoFinder; see below) and orthogroups with less than four paralogs per species. To identify conserved and novel *D. gyrociliatus* introns, we aligned each *D. gyrociliatus* protein against each annotated protein isoform of each orthologous gene of the abovementioned six species and added the intron positions into the alignments^138^. To identify high-confidence conserved intron positions, we required that a given *D. gyrociliatus* intron position was found at the exact position of the alignment and with the same phase (0, 1 or 2) in at least four out of six other species. To define high-confidence non-conserved (likely novel) introns, we required that a *D. gyrociliatus* intron position did not match an intron position with the same phase within 25 alignment positions in any of the other six species. To assess the impact of intron length on splicing efficiency on *D. gyrociliatus, S. maritima* and *H. sapiens*, we used RNA-seq-based quantifications of intron retention as previous described^139^ and implemented by *vast-tools*^140^. Only introns that had sufficient read coverage^139,140^ were used to calculate average PIR.

To quantify intron gain and loss in *D. gyrociliatus* we generated a database of homologous introns from 28 metazoan genomes (Supplementary Table 6), obtaining one-to-one orthologous genes using BUSCO v3^103^ (prot mode and 1e-04 evalue) and the OrthoDB v9^141^ dataset of 978 single-copy animal orthologs. We aligned the predicted peptides using MAFFT v7.310 G-INS-i algorithm^142^ and used Malin^143^ to identify conserved intron sites and infer their conservation status in ancestral nodes. We estimated the rates of intron gain and loss in each node with Malin’s built-in model maximum-likelihood optimization procedure. We used this model to estimate the posterior probabilities of intron presence, gain and loss in extant and ancestral nodes. For each node, we calculated the intron density expressed as introns per kbp of coding sequence (introns/CDS kbp), as follows:

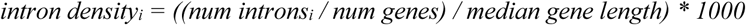

where *num introns*_*i*_ is the number of introns present in a given node (extant or ancestral, corrected by missing sites), *num genes* = 978 (number of alignments of one-to-one orthologs) and *median gene length* = 682.5 bp (as obtained from the lengths of the seed proteins curated in the OrthoDB v9 Metazoa dataset; ‘ancestral’ FASTA file). We used the same strategy to obtain the rates of intron gain and loss per node in terms of introns/CDS kbp. In addition, we inferred the uncertainty of the estimated intron gains, losses, and presence values with Malin and 1000 bootstrap iterations. To visualize the evolution of intron content, we used the *ape* library v5.0^144^ from the R statistical package v3.6 (R Core Team, 2017).

### Gene family evolution analyses

We used OrthoFinder v.2.2.7^145^ with default values to reconstruct clusters of orthologous genes between *D. gyrociliatus* and 27 other animal proteomes (Supplementary Table 6). OrthoFinder gene families were used to infer gene family gains and losses at different nodes using the ETE 3 library^146^. Gene expansions were computed for each species using a hypergeometric test against the median gene number per species for a given family. We used the functionally annotated gene sets of *D. gyrociliatus, C. teleta* and *H. robusta* to identify their repertoires of transcription factors, ligands and receptors. If a gene was not in the annotated *D. gyrociliatus* genome assembly, we performed manual search via BLAST on the *de novo* and genome guided transcriptome. For *T. axi* and *D. vorticoides*, gene identification was conducted on the assembled transcriptome via manual BLAST searches. To reconstruct KEGG pathways via KEGG Mapper^147^, we used the functionally annotations obtained from Trinotate to extract KEGG IDs. GPCR sequences in *D. gyrociliatus* and other animals (Supplementary Table 7) were retrieved using HMMER v3.2.1^123,148^ (e-value cutoff < 0.01) with Pfam profiles of class A (PF00001), class B (PF00002), class C (PF00003) and class F (PF01534) GPCRs (according to GRAFS classification). Sequences from each class were tested for false positives from other classes (including cAMP slime-mold class E GPCRs, PF05462). Phylogenetic analysis of GPCRs were performed as described elsewhere^149^. Neuropeptide candidates (Supplementary Data File 5) were retrieved by a combination of BLAST-searches (e-value cutoff < 0.1) and the use of a customised script^149^ to detect cleavage patterns on precursors.

### Orthology assignment

Multiple protein alignments were constructed with MAFFT v.7^142^, poorly aligned regions were either removed by hand or with gBlocks^150^. Maximum likelihood trees were constructed with FastTree 2^151^ using default parameters and visualised with FigTree.

### Gene expression analyses

*D. gyrociliatus* embryos were collected in their egg clusters and manually dissected. The embryonic eggshell was digested in a solution of 1% sodium thioglycolate (Sigma-Aldrich, T0632,) and 0.05% protease (Sigma-Aldrich, P5147) in sea water, pH 8, for 30 min at room temperature (RT), followed by relaxation in MgCl_2_ and fixation. Whole mount *in situ* hybridization (WMISH) was performed as described elsewhere^47^. Images were taken with a Zeiss Axiocam HRc connected to a Zeiss Axioscope Ax10 using bright-field Nomarski optics. *C. teleta* embryos were fixed and WMISH was performed as previously described^152^. *Capitella teleta* orthologs *Ct-fgf8/17/18/24*^153^ (Protein ID: 218971), *Ct-pvf1*^154^ (Protein ID: 153454) and *Ct-pvf2*^154^ (Protein ID: 220370) were mined from the publicly available genome^50^. WMISH samples were imaged on a Leica DMRA2 compound microscope coupled with a QIClick camera. Animals stained for F-actin were fixed for 30 min at RT, incubated with 1:100 BODIPY FL-Phallacidin (Life Technologies, cat #B607) and imaged with an IXplore SpinSR (Olympus). *Capitella* WMISH images were rendered using Helicon Focus (HelSoft). Contrast and brightness of images were edited with Photoshop (Adobe Systems, Inc.) when needed.

### Macrosynteny analysis

Single-copy orthologues obtained using the mutual best hit approach (MBH) were used to generate Oxford synteny plots comparing sequentially indexed orthologue positions as previously described^155^. Plotting order was determine by hierarchical clustering of the shared orthologue content using the complete linkage method^156^.

### Assay for Transposase-Accessible Chromatin using sequencing (ATAC-seq)

ATAC-seq libraries were performed as described elsewhere^157^, using 50,000-70,000 cells (~18 adult females). Cell dissociation and lysis was improved by disaggregating the tissue with a syringe in lysis buffer. Transposed DNA fragments were amplified by 16 cycles of PCR. Two biological replicates were sequenced in an Illumina NextSeq500 in rapid paired end mode and 75 base read length. Adapter contamination was removed with cutadapt v.1.2.1^95^ and cleaned reads were aligned to the unmasked genome with bowtie2^111^. Peaks were called with MACS2 v.2.1.1.20160309^158^ with the options --nomodel --extsize 70 --shift -30 --call-summits --keep-dup 1. Irreproducible discovery rates (IDR) were calculated with IDRCode^159^. A final set of 10,241 consistent peaks (IDR ≤ 0.05) was used for *de novo* motif enrichment analysis using HOMER^160^, with default parameters, except-size given (Supplementary Data File 6).

### Cap Analysis Gene Expression (CAGE)-seq

Total RNA from adult *D. gyrociliatus* was isolated using Trizol followed by RNeasy RNA clean-up protocol (Qiagen). CAGE libraries were prepared for two biological replicates (barcodes ATG and TAC) using the latest nAnT-iCAGE protocol^161^. The libraries were sequenced in single-end 50 base mode (Genomic Facility, MRC LMS). Demultiplexed CAGE reads (47 bp) were mapped to the *D. gyrociliatus* genome assembly using Bowtie2^162^ and resulting Bam files were imported into R using the standard CAGEr package (version 1.20.0) and G-correction workflow^163^. Normalization was performed using a referent power-law distribution^164^ and CAGE-derived TSSs that passed the threshold of 1 TPM were clustered together using distance-based clustering (Supplementary Data File 7). Genomic locations of tag clusters were determined using the ChIPseeker package and gene model annotations, where promoters were defined to include 500 bp upstream and 100 bp downstream of the annotated transcript start site. Visualization of motifs, sequence patterns, or reads coverage was performed using Heatmaps and seqPattern Bioconductor packages.

## Author contributions

J.M.M.-D., B.C.V., K.W. and A.H. conceived the study. J.M.M.-D., B.C.V. and A.H. designed experiments and analyses. J.M.M.-D. and V.C. performed collections and extractions. J.M.M.-D. and B.C.V. generated the genome and transcriptome assemblies. B.C.V. analysed the RNA-seq data. F.M. performed phylogenetic analyses and gene family evolution studies. V.C., A.K., N.B. and A.M.C.-B. performed gene expression analyses. W.G. performed flow cytometry analyses. N.C. and B.L. performed and analysed CAGE-seq. J.M.M.-D. and J.L.G.-S. performed and analysed ATAC-seq. J.M.M.-D., S.H., D.T., D.C., M.I., X.G.-B. and Y.M. performed computational analyses. All authors contributed to interpretation of the results, and J.M.M.-D., K.W. and A.H. wrote the manuscript.

## Data availability

All new raw sequence data associated to this project are available under project with primary accession PRJEB37657 in the European Nucleotide Archive (ENA). Genome annotation files and additional datasets are available in https://github.com/ChemaMD/DimorphilusGenome.

## Code availability

All custom code used in this study is freely available in https://github.com/fmarletaz/comp_genomics.

## Acknowledgements

We thank all members of the Hejnol lab for their continuous support, as well as the staff at the Norwegian Sequencing Centre (NSC) and GeneCore (EMBL) for their technical support. We also thank Rafa Domínguez Acemel, Marta Silva Magri and Sandra Jimenez Gancedo for their help with establishing ATAC-seq in *D. gyrociliatus*, Juan Tena-Aguilar for his help with ATAC-seq bioinformatic analyses, and Marina Ezcurra for providing live *C. elegans* worms for flow cytometry analyses. Finally, we thank Alexandre de Mendoza and Christopher Laumer for a critical read to this manuscript, as well as Mark Blaxter and two other anonymous reviewers for their comments and improvements. This study was supported by Sars Centre core budget and the European Research Council (ERC) Grant Agreement No. 648861 to A.H. J.M.M.-D. was additionally supported by the ERC Grant Agreement No. 801669, and B.C.V. by an EMBO Long-Term Fellowship (ALTF 74-2018). J.L.G.-S. received funding from the ERC (Grant Agreement No. 740041) and the Spanish Ministerio de Economía y Competitividad (Grant No. BFU2016-74961-P) and the institutional grant Unidad de Excelencia María de Maeztu (MDM-2016-0687).

---

**Extended Data Figure 1.**
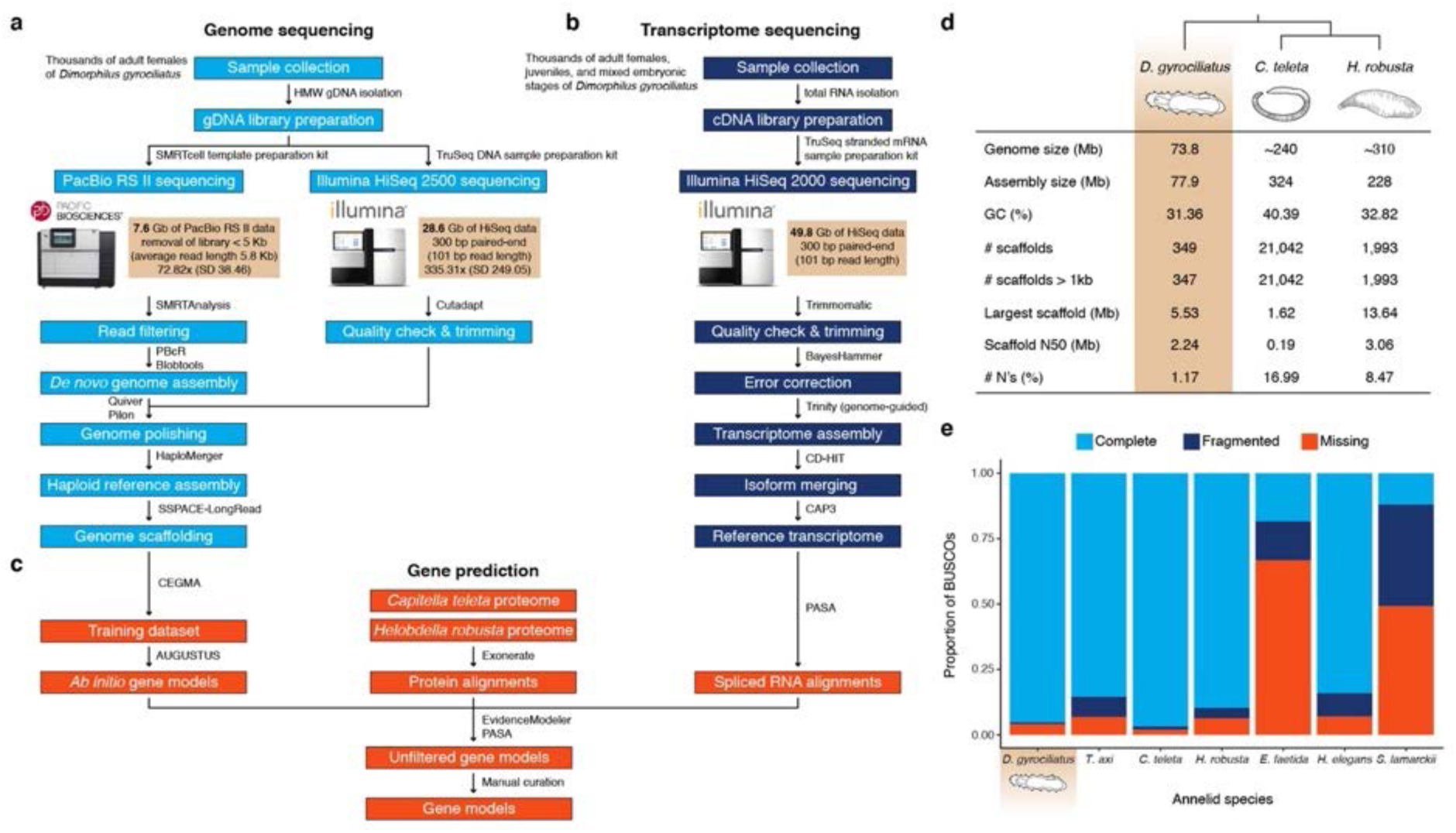
Sequencing approach and assembly statistics. (**a**–**c**) Diagram of the approach taken to sequence and assemble *Dimorphilus gyrociliatus* genome and transcriptome, and to annotate coding genes in the genome. (**d**) Comparison of genome assembly statistics between *D. gyrociliatus* and the annelids *C. teleta* and *H. robusta. D. gyrociliatus* genome is smaller than one third of *C. teleta*’s genome, and the assembly is contained in only ~350 scaffolds, with an N50 of 2.24 Mb, the second-best contiguity value for an annelid genome assembly to date. (**e**) Genome completeness, as indicated by metazoan BUSCO repertoire, in genome assemblies of different annelid lineages. *D. gyrociliatus* completeness is comparable to *C. teleta*, the most conservative annelid genome sequence to date.

**Extended Data Fig. 2.**
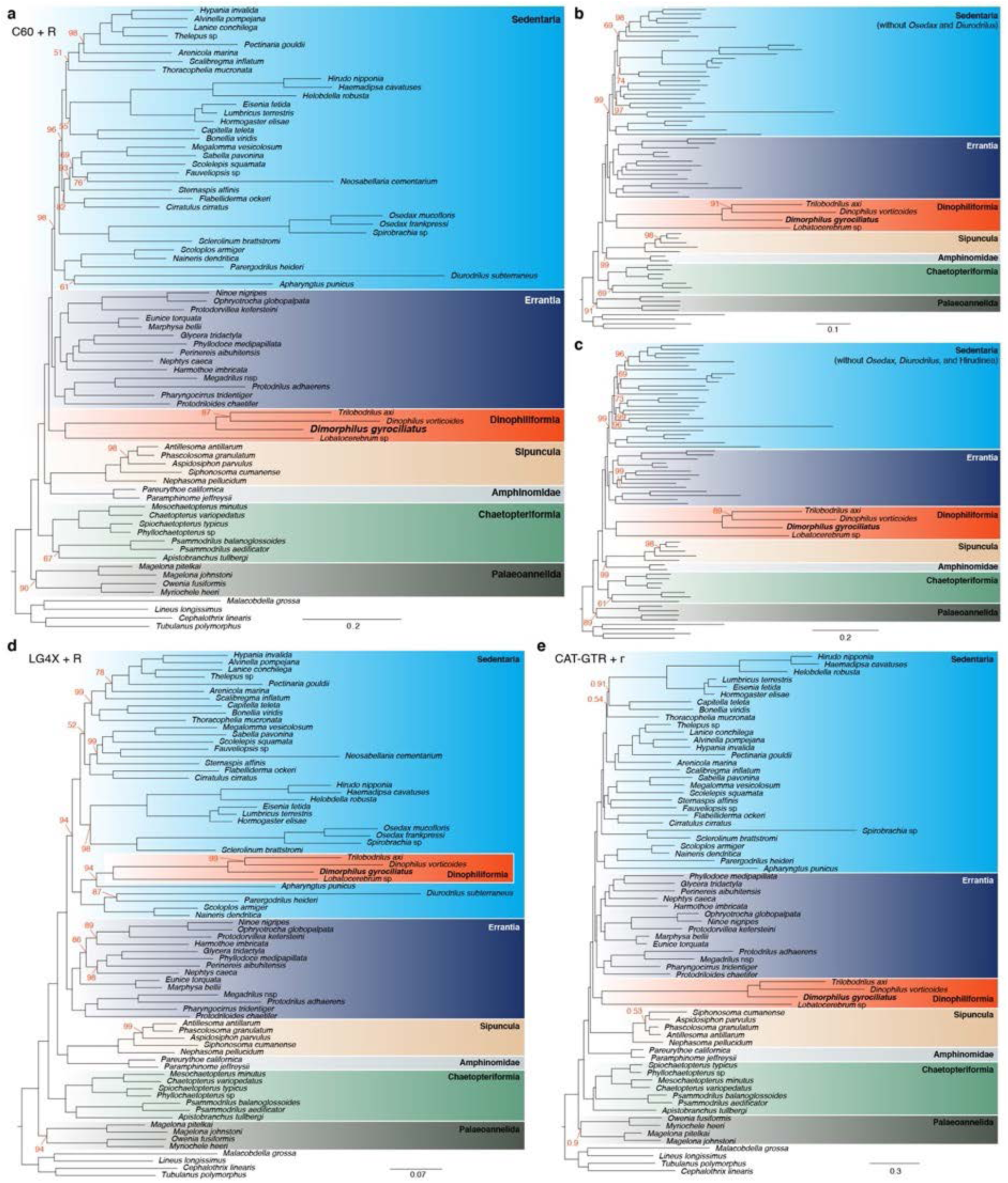
*Dimorphilus gyrociliatus* phylogenetic position. (**a**–**c**) Maximum likelihood phylogenetic tree using the site heterogeneous model of protein evolution C60 + R and the entire annelid dataset (**a**), excluding the fast evolving *Osedax* and *Diurodrilus* lineages (**b**), and additionally excluding Hirudinea (**c**). In all cases, *D. gyrociliatus* forms with *Trilobodrilus axi, Dinophilus vorticoides* and *Lobatocerebrum* sp. the clade Dinophiliformia, being this robustly placed as sister to Sedentaria + Errantia. (**d**) Maximum likelihood tree using the site homogeneous model of protein evolution LG4X + R and the entire annelid dataset. This condition recapitulates Dinophiliformia, but places this group inside Sedentaria, related to other fast evolving sedentarian lineages. (**e**) Bayesian phylogenetic tree using the site heterogeneous model of protein evolution CAT-GTR + G and excluding long branch lineages (*Osedax, Diurodrilus* and Hirudinea) recapitulates the maximum likelihood tree with the site heterogenous model and the same dataset. In all trees, only values other than 100 bootstrap or 1 posterior probability are shown.

**Extended Data Fig. 3.**
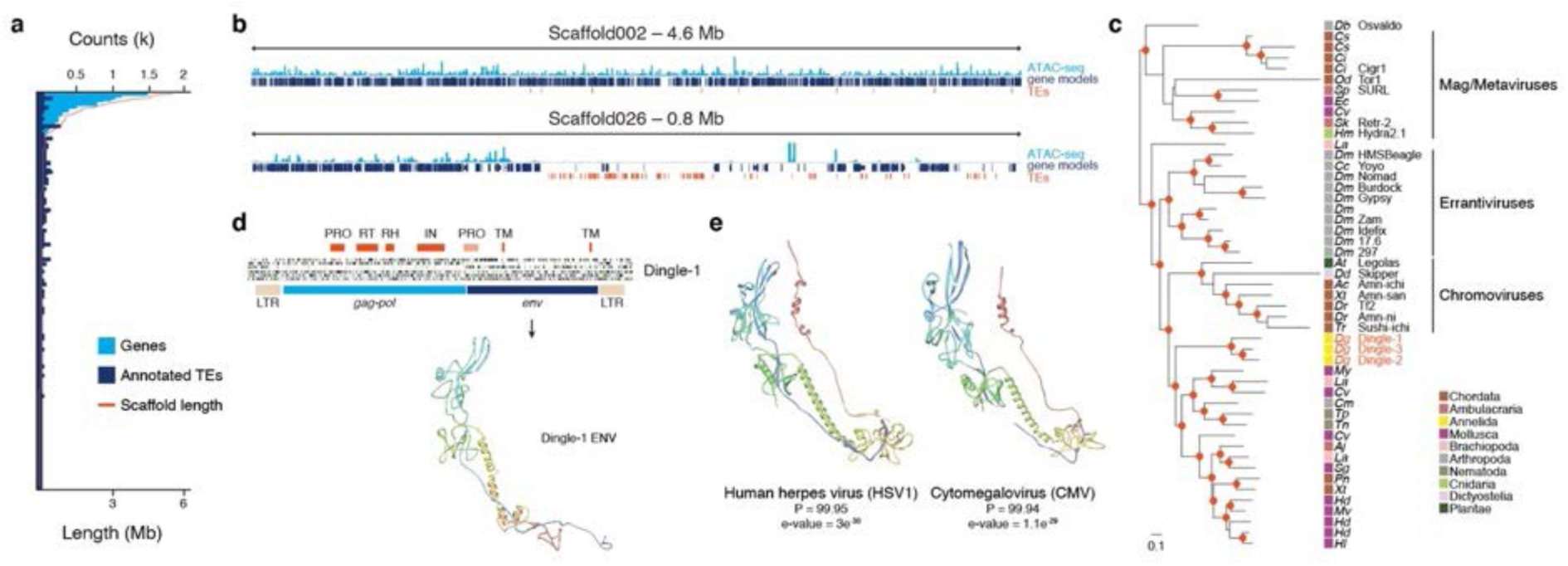
The transposable element repertoire of *Dimorphilus gyrociliatus*. (**a**) Graph showing the number of genes and transposable elements (TEs) per scaffold. (**b**) Diagram of Scaffold002 and Scaffold026 illustrating how transposable elements (TEs, in red) often concentrate in gene-free (dark blue boxes) and closed chromatin (as indicated by ATAC-seq signal; light blue) islands. (**c**) Maximum likelihood phylogeny of the *pol* gene showing that Dingle is a new family of *Ty3/gypsy* LTR element. The scale bar shows the number of substitutions per site and red dots are bootstrap values > 0.7. (**d**) Genetic organization of Dingle, showing protein domains (top red boxes), 6-frame translations (green lines, ATG; black lines, stop codons) and the predicted protein structure of ENV, which shows resemblance to that of human herpes viruses (**e**). In (**c**), Ac, *Anolis carolinensis*; Aj, *Apostichopus japonicus*; At, *Arabidopsis thaliana*; Cc, *Ceratitis capitata*; Ci, *Ciona intestinalis*; Cm, *Callosobruchus maculatus*; Cs, *Ciona savignyi*; Cv ; *Crassostrea virginica*; Db, *Drosophila buzzati*; Dd, *Dictyostelium discoideum*; Dg, *Dimorphilus gyrociliatus*; Dm, *Drosophila melanogaster*; Dr, *Danio rerio*; Ec, *Elliptio complanata*; Hd, *Haliotis discus hannai*; Hl, *Haliotis laevigata*; Hm, *Hydra_magnipapillata*; La, *Lingula anatina*; Mv, *Mimachlamys varia*; My, *Mizuhopecten yessoensis*; Od, *Oikopleura dioica*; Pn, *Pundamilia nyererei*; Sg, *Saccostrea glomerata*; Sk, *Saccoglossus kowalevskii*; Sp, *Strongylocentrotus purpuratus*; Tn, *Trichinella nelson*; Tp, *Trichinella pseudospiralis*; Tr, *Takifugu rubripes*; Xt, *Xenopus tropicalis*.

**Extended Data Fig. 4.**
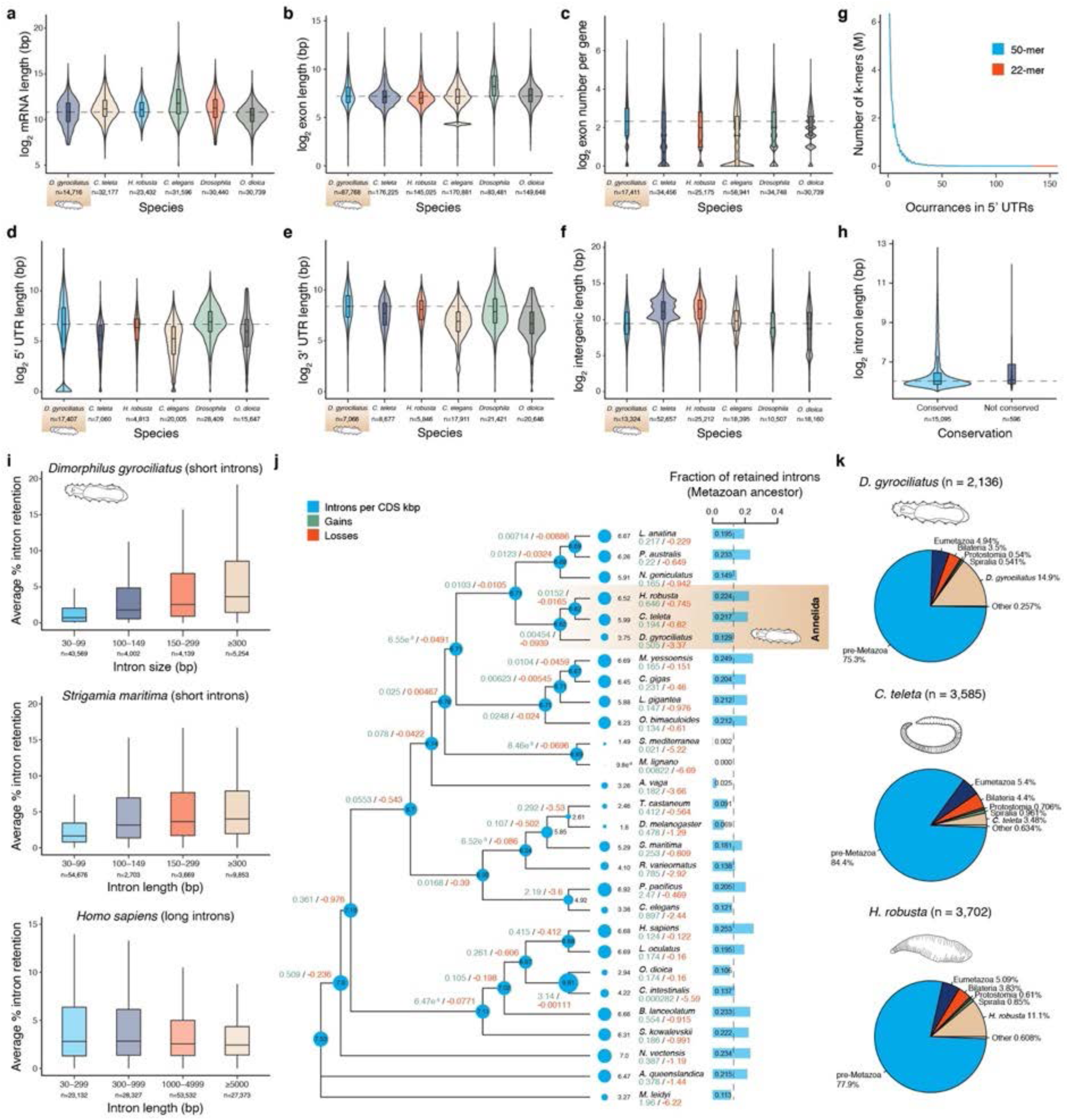
Comparative analyses of gene structure features in the *Dimorphilus gyrociliatus* genome. (**a**–**f**) Violin plots showing the genome-wide distribution of mRNA and exon lengths, exon numbers per gene, and the lengths of 5’ UTR, 3’UTR and intergenic regions in *D. gyrociliatus*, the annelids *C. teleta* and *H. robusta*, and the bilaterians with compact genomes *C. elegans, D. melanogaster* and *O. dioica*. (**g**) The distribution of occurrences of 22-mer and 50-mer in RNA-seq-based 5’ UTR regions of *D. gyrociliatus* does not indicate the presence of overrepresented sequences that could act as splice leaders. (**h**) Violin plot showing the distribution of intron sizes between conserved and non-conserved introns in *D. gyrociliatus*. (**i**) The percentage of intron retention according to intron size demonstrates that the splicing machinery in *D. gyrociliatus* is adapted to short introns, as it occurs in the centipede *S. maritima* (also with short introns) and inversely to what is observed in *H. sapiens*, a species with longer introns. (**j**) Metazoan-wide analysis of intron density, intron gain and intron loss rates per lineage and their ancestors. Intron density (blue circles) are indicated at each node and terminal tip of the phylogram. Net intron gains and losses are indicated below the species name, together with the fraction of introns conserved in each extant genome, among the ones inferred to have been present at the last metazoan common ancestor. *D. gyrociliatus* shows rates of intron loss and retention of ancestral introns similar to other animal lineages with much larger genomes. (**k**) Inferred origin of the intron sites in *D. gyrociliatus* and the annelid *C. teleta* and *H. robusta*, expressed as the sum of gain probabilities at their respective ancestral nodes.

**Extended Data Fig. 5.**
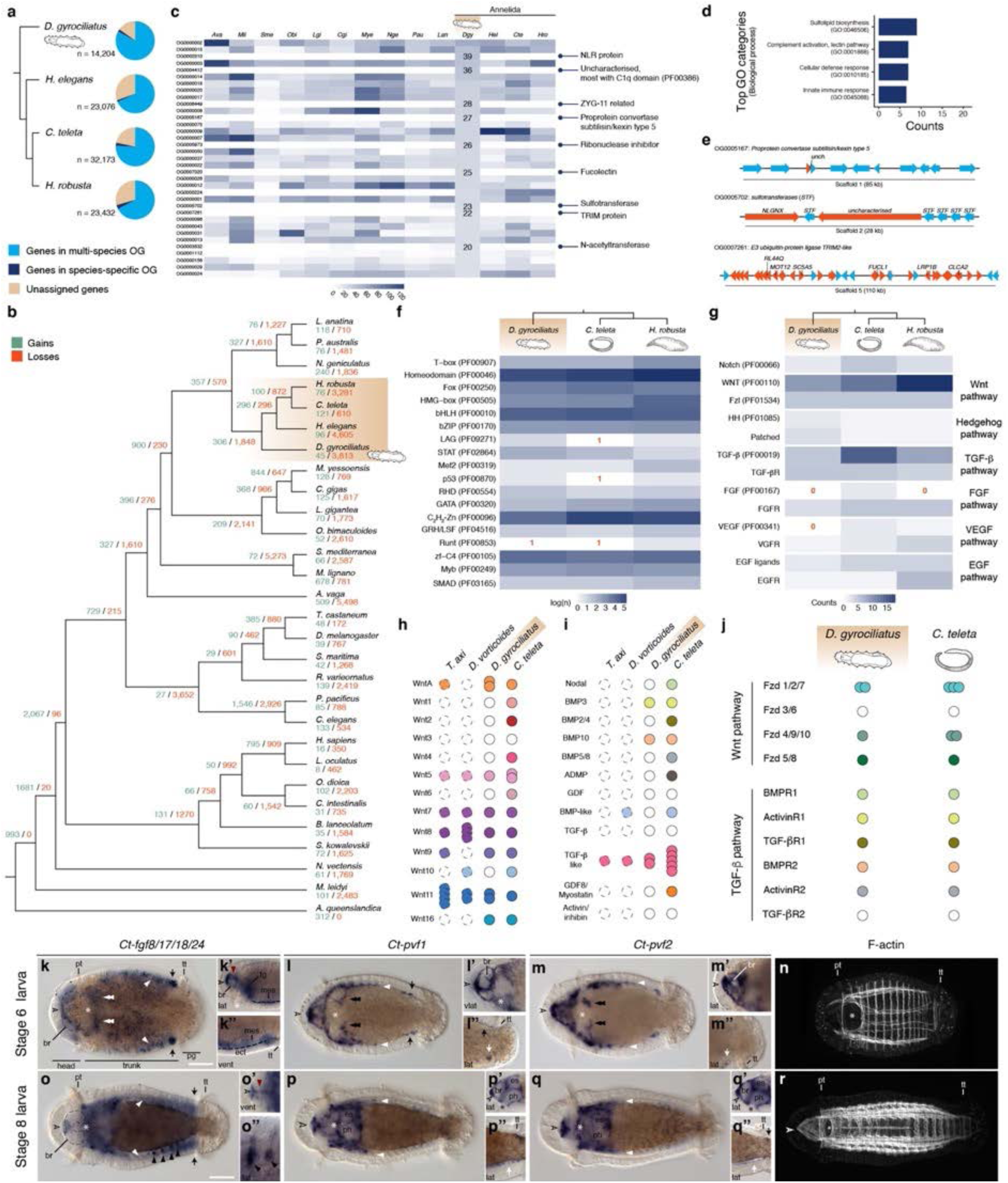
Expansions and gene losses in the genome of *Dimorphilus gyrociliatus*. (**a**) Orthology analyses of G-protein coupled receptors (GPCRs) for each class. The magenta asterisks highlight *D. gyrociliatus* receptors, and the annotations are given based on the *D. melanogaster* orthology. In (**a**) 5HT, serotonin; Ado, Adenosin; Akh, adipokinetic hormone; AstA, allatostatin A; AstC, allatostatin C; Boss, bride of sevenless; Capa, capability; Ccap, crustacean cardioactive peptide; CCHa, CCHamide; Cck, cholecystokinin; CNMa, CNMamide; Crz, corazonin; Dh, diuretic hormone; Dop, dopamine; Ec, ecdysteroid; ETH, ecdysis triggering hormone; FMRFa, FMRFamide; Fz, Frizzled; Glut, glutamate; Lrrc, leucine rich repeat containing; mAch, muscarinic acetylcholine; Mthl, methuselah; mtt, mangetout; Myos, myosuppressin; NpF, neuropeptide F; Oct, octopamine; Oct-R-mb, Octopamin receptor in mushroom bodies; Pdf, pigment dispersing factor; R, receptor; Rh, rhodopsin; RYa, RYamide; Sexp, sex peptide; SIFa, SIFamide; Smo, Smoothened; sNpF, short neuropeptide F; stan, starry night; Tre, trapped in endoderm; Tyr, tyramine. (**b**) Phylogram with the number of GPCRs per class in representative animal species. Contrary to other animals with compact genomes and miniaturised morphologies, such as tardigrades, nematodes and appendicularians, *D. gyrociliatus* has a conserved GPCR repertoire. (**c**) PSI-BLAST cluster map of *D. gyrociliatus* pro- neuropeptides, each dot corresponding to one sequence, their colour corresponds to the legend in upper left corner. Connections are based on e-values < 1e-7 (see upper right corner). In (**c**), a, amide; ast, allatostatin; crz, corazonin; ct, calcitonin; dh, diuretic hormone; elh, ecdysis triggering hormone; ep, excitatory peptide; glyho-a, glycoprotein hormone alpha; glyho-b, glycoprotein hormone beta; gnrh, gonadotropin releasing hormone; ilp, insulin like peptide; myom, myomodulin; np-F, neuropeptide F; np-Y, neuropeptide Y; npl, neuropeptide-like; pdf, pigment dispersing factor; pedpep, pedal peptide; scap, short cardioactive peptide.

**Extended Data Fig. 6.**
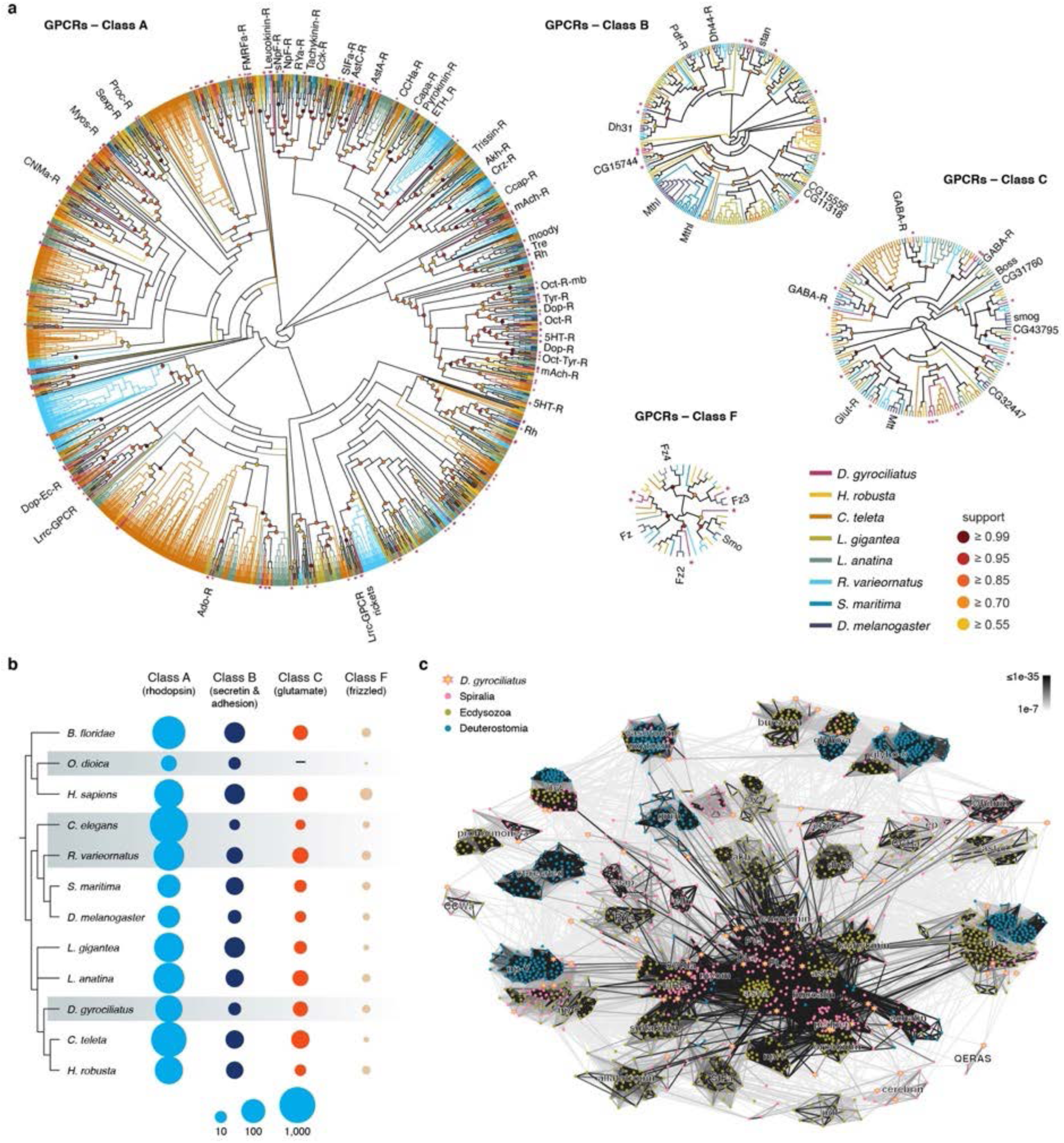
The GPCR and neuropeptide repertoire of *Dimorphilus gyrociliatus*. (**a**) Orthology analyses of G-protein coupled receptors (GPCRs) for each class. The magenta asterisks highlight *D. gyrociliatus* receptors, and the annotations are given based on the *D. melanogaster* orthology. In (**a**) 5HT, serotonin; Ado, Adenosin; Akh, adipokinetic hormone; AstA, allatostatin A; AstC, allatostatin C; Boss, bride of sevenless; Capa, capability; Ccap, crustacean cardioactive peptide; CCHa, CCHamide; Cck, cholecystokinin; CNMa, CNMamide; Crz, corazonin; Dh, diuretic hormone; Dop, dopamine; Ec, ecdysteroid; ETH, ecdysis triggering hormone; FMRFa, FMRFamide; Fz, Frizzled; Glut, glutamate; Lrrc, leucine rich repeat containing; mAch, muscarinic acetylcholine; Mthl, methuselah; mtt, mangetout; Myos, myosuppressin; NpF, neuropeptide F; Oct, octopamine; Oct- R-mb, Octopamin receptor in mushroom bodies; Pdf, pigment dispersing factor; R, receptor; Rh, rhodopsin; RYa, RYamide; Sexp, sex peptide; SIFa, SIFamide; Smo, Smoothened; sNpF, short neuropeptide F; stan, starry night; Tre, trapped in endoderm; Tyr, tyramine. (**b**) Phylogram with the number of GPCRs per class in representative animal species. Contrary to other animals with compact genomes and miniaturised morphologies, such as tardigrades, nematodes and appendicularians, *D. gyrociliatus* has a conserved GPCR repertoire. (**c**) PSI-BLAST cluster map of *D. gyrociliatus* pro-neuropeptides, each dot corresponding to one sequence, their colour corresponds to the legend in upper left corner. Connections are based on e-values < 1e-7 (see upper right corner). In (**c**), a, amide; ast, allatostatin; crz, corazonin; ct, calcitonin; dh, diuretic hormone; elh, ecdysis triggering hormone; ep, excitatory peptide; glyho-a, glycoprotein hormone alpha; glyho-b, glycoprotein hormone beta; gnrh, gonadotropin releasing hormone; ilp, insulin like peptide; myom, myomodulin; np-F, neuropeptide F; np-Y, neuropeptide Y; npl, neuropeptide-like; pdf, pigment dispersing factor; pedpep, pedal peptide; scap, short cardioactive peptide.

**Extended Data Fig. 7.**
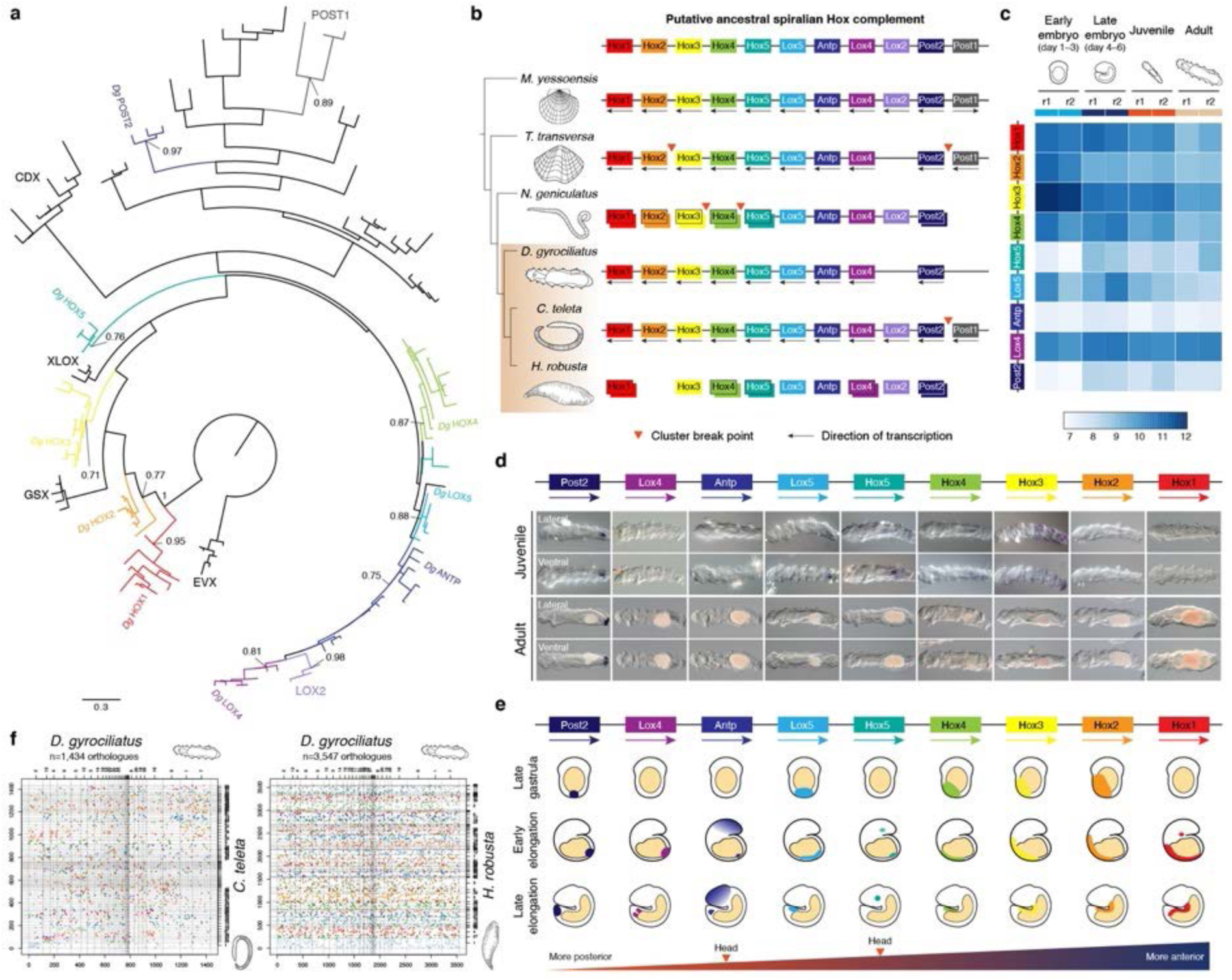
The Hox cluster of *Dimorphilus gyrociliatus*. (**a**) Maximum likelihood tree of Hox and ParaHox genes, with Evx proteins as outgroup, to assign orthology relationships of *Dimorphilus* Hox genes (indicated in the tree). Only bootstrap values for main orthogroups are shown. (**b**) Schematic representation of Hox complements and Hox genomic organisations in representative spiralian species, with the putative ancestral spiralian Hox cluster on the top. Each Hox orthologous group is coloured differently. (**c**) Heatmap of gene expression values of *Dimorphilus* Hox genes during the life cycle. (**d**) Whole mount in situ hybridisation of Hox genes in *Dimorphilus* juveniles and adults. Only *Post2* and *Hox3* show conspicuous expression domains in the hindgut and posterior ectoderm of the juvenile, respectively. In adults, we only detect expression of *Post2* in the hindgut. (**e**) Schematic summary of Hox gene expression in relation to the Hox genomic organisation during *Dimorphilus* embryonic development. Hox genes exhibit an anteroposterior spatial collinearity along *Dimorphilus* trunk, with *Antp* and *Hox5* being additionally expressed in head domains. However, Hox genes do not exhibit temporal collinearity, as all but *Hox5, Antp*, and *Lox5* become expressed by the end of gastrulation. (**f**) Oxford dot plots of orthologous genes between *D. gyrociliatus, C. teleta* and *H. robusta*. Macrosyntenic relationships are little conserved between annelid worms, indicating lineage-independent large-scale genomic reorganisations.

**Extended Data Fig. 8.**
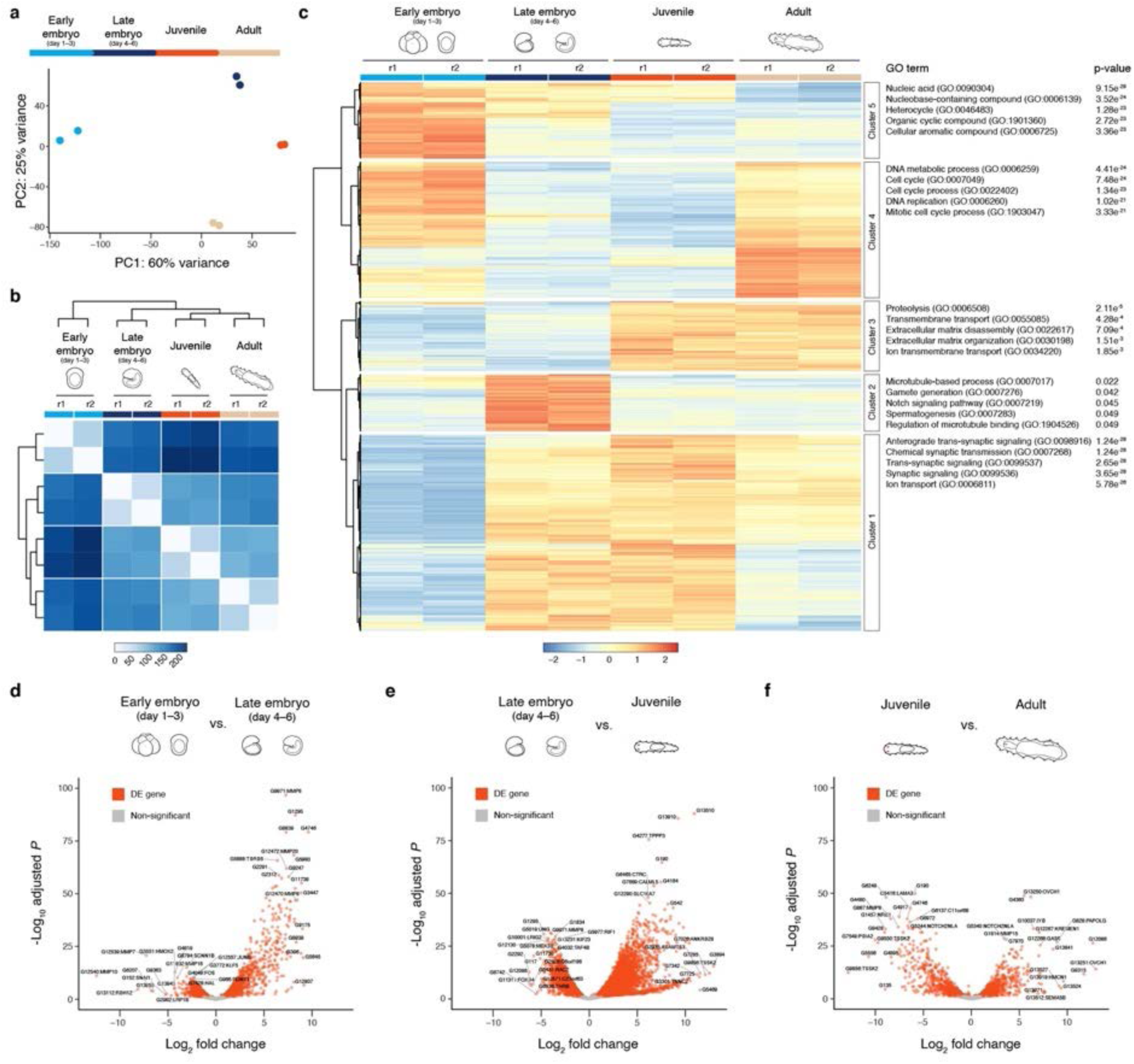
Differential expression analyses during the life cycle of *Dimorphilus gyrociliatus*. (**a**) Principal component analysis of the stage-specific RNA-seq samples using the top eight thousand most-variable genes. The raw count data was transformed to homogenize the variance and normalised using the variance stabilising method from DESeq2. (**b**) Euclidean distances between the variance stabilized normalized counts of the stage-specific RNA-seq samples. (**c**) Expression patterns for the top three thousand differentially expressed genes. Variance stabilised normalised counts were scaled around the mean value of the row to highlight changes in expression between developmental stages. Gene ontology terms associated with each cluster of expression profile are shown on the right. (**d**–**f**) Differentially expressed genes from pairwise Wald tests between stage-specific RNA-seq samples. The top 18 genes with lowest p-adjusted values and highest log fold change are labelled. Considering gene expression changes significant if the adjusted p-value < 0.05, we identified 8,341 differentially expressed genes (4,543 up and 3,798 down) for “late embryo vs early embryo”; 1,870 genes (938 up and 932 down) for “juvenile vs late embryo”; and 3,746 genes (1,827 up and 1,919 down) for “adult vs juvenile”.

**Extended Data Fig. 9.**
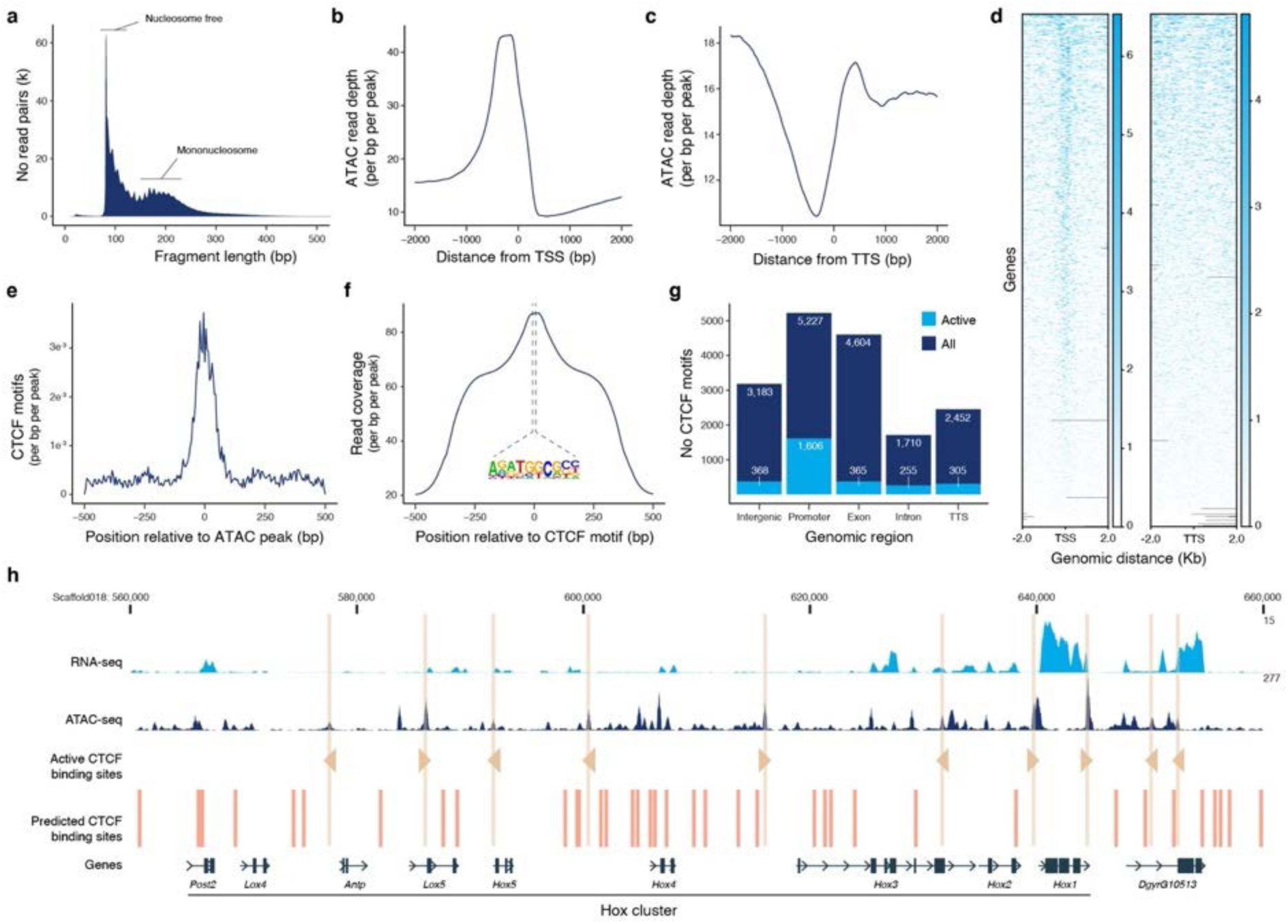
CTCF-binding motifs are the most abundant in open chromatin regions in *Dimorphilus gyrociliatus*. (**a**) Insert size distribution of ATAC-seq samples in *D. gyrociliatus*. (**b, c**) Averaged ATAC-seq read depth around transcription start sites (TSS) and transcription termination sites (TTS). (**d**) Heatmaps of ATAC-seq read coverage around TSS (left) and TTS (right) of each annotated gene. (**e**) Averaged location of CTCF motifs in ATAC-seq peaks. (**f**) Aggregate ATAC-seq read coverage centred around CTCF motifs. (**g**) Number of CTCF motifs according to genomic feature. Most CTCF binding motifs in open chromatin regions (i.e. “active”) are in promoters. (**h**) Genome browser snapshot showing the distribution of CTCF binding motifs in the Hox cluster of *D. gyrociliatus* as example of the general pattern observed genome wide. Most often, there is only one CTCF motif in an open chromatin region, and there is no clear directional arrangement between consecutive or neighbouring active CTCF binding sites.

**Extended Data Fig. 10.**
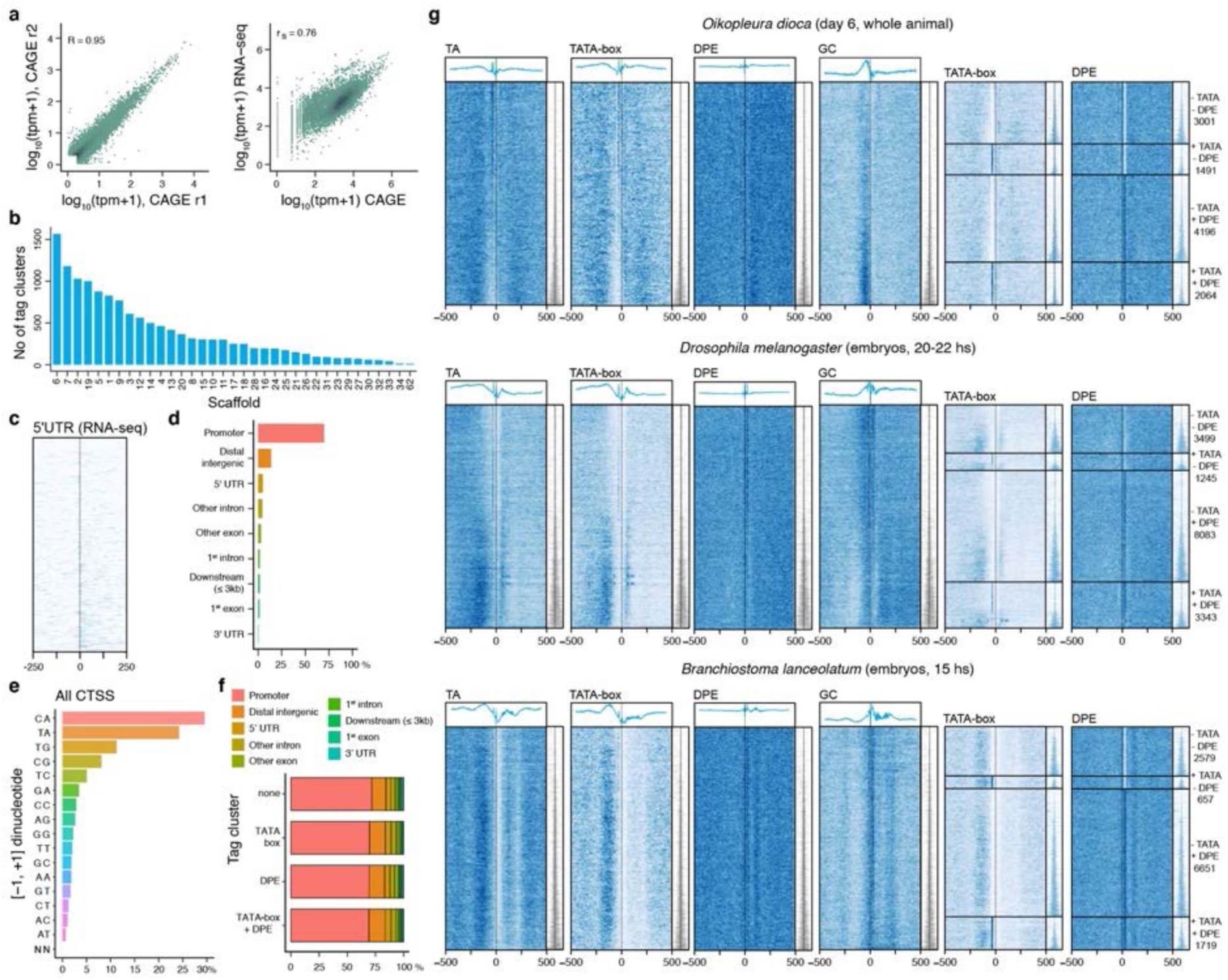
General features and comparative aspects of CAGE-seq derived promoters in *Dimorphilus gyrociliatus*. (**a**) Pearson’s correlation at the CAGE-supported transcription start site (CTSS) level between CAGE-seq biological replicates (left panel) and Spearman correlation between gene-counts derived from RNA- or CAGE-seq (right panel, merged biological replicates). (**b**) Distribution of number of tag clusters/promoters across scaffolds. (**c**) Heatmap of tag-cluster coverage ordered by tag-cluster IQ-width from narrow (top) to broad (bottom) centred on the first nucleotide of 5’ UTRs determined by RNA-seq. (**d**) Genomic locations of dominant CTSS. (**e**) Dinucleotide composition of all CTSSs identified in *Dimorphilus* CAGE-seq libraries. (**f**) Genomic locations of tag clusters identified to contain a TATA-box or downstream promoter element (DPE). (**g**) Sequence patterns in CAGE-seq derived promoters in the appendicularian *O. dioica* (genome size ~70 Mb), the fly *D. melanogaster* (genome size ~140 Mb) and the lancelet *B. lanceolatum* (genome size ~550 Mb). All heatmaps are centred on dominant TSSs and ordered by the tag-cluster/promoter IQ-width from narrower (top) to broader (bottom). IQ-widths are shown as tag cluster coverage in the same order as on the heatmaps (right, in grey or blue). Heatmaps (left to right) represent TA dinucleotide patterns, TATA-box or DPE density (promoter regions are scanned using a minimum of the 80th percentile match to the TATA-box or DPE position weight matrix (PWM)) or GC dinucleotide patterns. Relative signal metaplot is shown above each heatmap. Promoters are divided according to TATA-box or DPE content at −30 or + 30 position relative to the dominant TSS, and a heatmap of TATA-box or DPE density across promoter categories is shown.

