## Supplementary Material Summary for "Conservative route to genome compaction in a miniature annelid" (from another version of this preprint: version 1, 2020-05-08)

**Index:**

- **Supplementary Methods**
- **Supplementary Figures 1–4.** Gene orthology assignments.
- **Supplementary Tables 1–9.**
- **Supplementary Notes 1 and 2.**
- **Supplementary References**
- **Supplementary Data File 1.** Automatic annotation of transposable elements and repeats (gff format file).
- **Supplementary Data File 2.** Automatic annotation of transposable elements and repeats (gff format file).
- **Supplementary Data File 3.** Filtered gene models (gtf format file).
- **Supplementary Data File 4.** Functional annotation of *D. gyrociliatus* filtered gene models (excel spreadsheet).
- **Supplementary Data File 5.** Node-specific rates of intron gains and losses (excel spreadsheet).
- **Supplementary Data File 6.** List of *D. gyrociliatus* neuropeptide precursors (docx format file).
- **Supplementary Data File 7.** Functional annotation *C. teleta* gene models (excel spreadsheet).
- **Supplementary Data File 8.** Functional annotation *H. robusta* gene models (excel spreadsheet).
- **Supplementary Data File 9.** IDR reproducible ATAC-seq peaks (bed format file).
- **Supplementary Data File 10.** Motif annotation of ATAC-seq peaks (txt format file).
- **Supplementary Data File 11.** Raw CAGE-seq tag clusters, replicate 1, plus strand (bedGraph format file).
- **Supplementary Data File 12.** Raw CAGE-seq tag clusters, replicate 1, minus strand (bedGraph format file).
- **Supplementary Data File 13.** Raw CAGE-seq tag clusters, replicate 2, plus strand (bedGraph format file).
- **Supplementary Data File 14.** Raw CAGE-seq tag clusters, replicate 2, minus strand (bedGraph format file).
- **Supplementary Data File 15.** Processed CAGE-seq tag clusters (txt format file).

**Supplementary Methods**

1. **Genome sequencing and assembly**
   1.1. *Material and genomic DNA extraction*

Thousands of adult females of *Dimorphilus gyrociliatus* (O. Schmidt, 1857) (see Supp. Note 1 for more information on the culture and species) were starved for 3–4 days, washed thoroughly in sea water (~28ppm salinity), and snap frozen in liquid nitrogen. Genomic DNA (gDNA) was extracted with guanidium isothiocyanate, purified with a standard phenol chloroform isoamyl alcohol protocol and precipitated with ethanol. After this initial gDNA extraction, samples were treated with 20 mg/mL of RNase A for 1 h at 37 ºC. Before library preparation, RNase A treated gDNA was further purified with phenol chloroform isoamyl alcohol extraction and precipitated with ethanol.

1.2. *Library preparation, sequencing and assembling*

Library was prepared using Pacific Biosciences 20 kb library preparation protocol and size-selected using BluePippin with 5 kb cut-off. The library was sequenced on a Pacific Bioscience RS II instrument using P6-C4 chemistry at the Norwegian Sequencing Centre (NSC). A total of 8 SMRT cells were used for sequencing, four with 240 min and four with 360 min movie time. Sequencing produced a total of 487,071 reads, with an N50 length of 5,842 bp, representing a total of 7,642.72 Mb. In addition, an Illumina library of median insert size of 298 bp was sequenced in 101 bp paired end mode on a full lane of an Illumina HiSeq 2500 instrument at GeneCore (EMBL). Sequencing produced 141,662,088 read pairs that amount to a total of 28,615.74 Mb. All raw sequence data associated to this project is available under project with primary accession PRJEB37657 in the European Nucleotide Archive (ENA).

PacBio reads were filtered with SMRTAnalysis v.2.3.0.140936 and assembled with PBcR v.8.3rc2^67,68^ using default options, except for k-mer=14 and asmMerSize=14 (Extended Data Figure 1a). Four rounds of decontamination using Blobtools v.0.9.16^69^ were applied to the initial assembly. For each round, contaminant PacBio subreads were removed and a new assembly with PBcR using the same initial settings was conducted. Out of the different rounds of decontamination, the final assembly had the maximum proportion of bases (83.73%) annotated as belonging to animals. Thereafter, a consensus assembly was generated with Quiver under the SMRTAnalysis portal and improved with Pilon v.1.16^70^ using the Illumina paired end reads, which were previously filtered for adapters with cutadapt v.1.4.2^71^. We used HaploMerger2 v.20151124^72,73^ to reconstruct a high-quality haploid reference assembly, which we further scaffolded with SSPACE-LongRead v.1.1^74^.

1.3. Trilobodrilus axi *genome sequencing and assembly*

We collected hundreds of *Trilobodrilus axi* Remane, 1925 individuals and performed a gDNA extraction as described above for *D. gyrociliatus*. The gDNA was used to prepare a TruSeq v3 Illumina library that was sequenced in 101 bp paired end mode on a full lane of an Illumina HiSeq 2500 instrument at GeneCore (EMBL). We removed adapters and low quality regions with cutadapt v.1.4.2^71^ and trimmomatic v.0.35^75^ respectively, and performed error correction of the reads with SPAdes v.3.6.2^76^. After deduplication of PCR duplicates with Super_Deduper v.2.0 (https://dstreett.github.io/Super-Deduper/), reads were assembled with Platanus v.1.2.4^77^ and contigs with similarity to proteobacteria were removed with Blobtools v.0.9.16^69^.

1.4. *Assembly statistics, read coverage, and validation*

We used Quast v.3.1^78^ to calculate the assembly statistics of the decontaminated, scaffolded haploid assembly of *D. gyrociliatus* in relation to published annelid genomes (Supplementary Table 1). The decontaminated PacBio and Illumina reads were re-aligned to the final assembly with BWA-MEM v.0.7.12-r1044^79^ and the read coverage was calculated with SAM tools^80^. The average PacBio coverage was 72.82x (SD 38.46) and the average Illumina coverage was 335.31x (SD 249.05). To validate the completeness of the genome assembly, we run the BUSCO v2 pipeline^81^ for genomic data using the metazoan-specific dataset. Out of the 978 BUSCO genes, 930 were complete (95.1%), 7 were fragmented (0.7%) and 41 were missing (4.2%) (Extended Data Figure 1e). We next used BLAT^82^ to align the assembled transcriptome to the genome and the script Isoblat v.0.3^83^ to calculate coverage statistics. 92% of transcripts aligned to the genome, with 96.9% of bases of the transcriptome covered. In total, 93.53% of the transcripts aligned to a single scaffold, and there was on average 1.1 scaffolds per mapped transcript. BUSCO analysis on the *T. axi* genome resulted in 835 complete (85.4%), 75 fragmented (7.7%) and 68 missing (6.9%) (Extended Data Figure 1e).

1.5. *Polymorphism estimation*

To estimate the heterozygosity of the genome, we used GATK and followed its best practices pipeline^84,85^. First, we aligned the Illumina paired end reads to the reference haploid assembly with the BWA-MEM v.0.7.12-r1044 algorithm^79^. Picard tools v.2.0.1 (http://broadinstitute.github.io/picard) were used to mark duplicates before realignment and SNP calling with GATK^86^. After running HaplotypeCaller, the raw variants were hard filtered with VariantFiltration using default values. A total of 29,100 SNPs passed the filter. To calculate the SNP rate, we used the GATK’s function CallableLoci to calculate the number of eligible sites, which amount to 29,059,634 bases. These two values together made a heterozygous SNP rate of 0.001 per site.

2. **Genome size measurements**
   2.1. *Flow cytometry*

Hundreds of adult females of *D. gyrociliatus* were collected and starved for 3–4 days before flow cytometry analysis. For calibration and genome size comparison, we used hundreds of *C. elegans* worms, which were similarly starved. *Dimorphilus* and *C. elegans* worms were transferred separately into a petri dish, washed well in their respective media to remove any contaminant, and finely chopped with a razor blade in 2 ml of General-Purpose Buffer^87^ to generate a suspension of nuclei. This suspension was filtered through a 30 µm nylon mesh and stained with propidium iodide (Sigma; 1 mg/mL) on ice. We used a flow cytometer Partex CyFlow Space fitted with a Cobalt Samba green laser (532nm, 100mW) to analyse the samples. For each sample we made three independent runs analysing at least 1,000 nuclei.

2.2. *K-mer based genome size estimation*

We pre-processed the raw Illumina paired-end reads by (1) trimming adapters using cutadapt v.1.4.2^71^, (2) discarding contaminant reads by aligning them to the genome using BWA-MEM v.0.7.12-r1044^79^, (3) correcting sequencing errors using SPAdes v.3.6.2^76^ with the argument “–only-error-correction”, and (4) deduplicating reads using Super-Deduper v.2.0 (https://dstreett.github.io/Super-Deduper/). We then used Jellyfish v.2.2.3^88^ to count the 31-mer frequency in the processed data which contained 126,850,236 paired-end 101bp reads. We estimated the genome size by dividing the total number of bases by the sequencing depth. To calculate the depth, we identified the peak coverage of unique 31-mers (139x; Figure 1d), which corresponds to the homozygous peak, and estimated the haploid genome size of *D. gyrociliatus* in 64,794,005 bases or 64.8 Mbp following the formula described elsewhere^89^. We applied the same pipeline to Illumina data from the closely related dinophilid *T. axi* and found a haploid genome size of 86,688,406 or 86.7 Mbp.

3. **Transcriptome sequencing and assembly**
   3.1. *RNA preparation and sequencing*

Adult specimens of *D. gyrociliatus*, juveniles and mixed developmental stages were collected and pooled together for total RNA extraction. Samples were flash frozen on liquid nitrogen before storage at -80ºC. Total RNA was isolated with TRI Reagent® Solution (Applied Biosystems) following manufacturer’s recommendations. Total RNA was directionally sequenced at GeneCore (EMBL) on an Illumina HiSeq2000. Sequencing generated a total of 246,574,111 paired end reads of 101 bp length and 300 bp insert size. Sequencing data is available in the Sequence Read Archive, accession number SRX2030658.

3.2. *De novo and genome-guided transcriptome assembly*

We first removed low-quality read pairs (<28) and read pairs contaminated with Illumina sequencing adapters using Trimmomatic^75^. About 89% (218,359,922) read pairs passed this initial quality filter. Only 8% (17,612,544) of these reads had sequencing errors, which were detected and corrected in 19,336,941 bases by BayesHammer^90^. To create a *de novo* transcriptome assembly, we used the corrected reads as input for Trinity v2.1.1^91^ using the *--normalize_reads* parameter to reduce memory requirements and assembly runtime while maintaining relevant data. To perform a genome-guided transcriptome assembly, we first normalized the corrected reads to 50x coverage using Trinity v2.1.1’s utility script *insilico_read_normalization.pl*^91^. This reduced the data to 13,141,727 read pairs. We then aligned the normalized reads to *D. gyrociliatus* genome assembly (v1) using Bowtie2^92^, sorted and indexed the alignments to BAM using samtools^80^, and finally loaded the BAM file as the input for Trinity’s genome-guided assembly mode (v2.1.1). The initial *de novo* and genome-guided assemblies had 125,427 and 134,853 transcripts, respectively. To remove redundant contigs, we selected representative sequences using *cd-hit-est* program with default parameters of CD-HIT^93^. This reduced the number of transcripts to 91,489 (*de novo*) and 78,132 (genome-guided), about 2/3 of the initial assembly sizes. We further reduced the number of transcripts by assembling 7,919 fragments into 3,622 transcripts for the *de novo* assembly and 6,888 fragments into 2,887 transcripts for the genome-guided assembly using CAP3^94^.

The resulting *de novo* transcriptome assembly of *D. gyrociliatus* contains 87,192 transcripts of which 49,945 correspond to unigenes (Trinity genes + CAP3 contigs; Supplementary Table 2). The genome-guided assembly contains 74,131 transcripts and 39,432 unigenes (Supplementary Table 2). The standard statistics shown in Supplementary Table 2 were calculated using Transrate^95^. To evaluate how well the reads were assembled, we used Bowtie2 to map the input reads to the contigs. We found the overall alignment rate was about 98% and 90% for the *de novo* and genome-guided assembly, respectively, indicating that the input reads are well represented in the transcriptomes. The high quality of the assembly was corroborated by Nx statistics, such as the E90N50 (N50 for the top highly expressed transcripts representing 90% of the expression data) of 4.0kb in the *de novo* assembly and 4.2kb in the genome-guided assembly, calculated using Trinity’s *contig_ExN50_statistic.pl* script. Finally, we evaluated the completeness of both assemblies using BUSCO v2^81^ and found a high percentage of metazoan genes (96%) represented in the *D. gyrociliatus* transcriptome.

3.3. *Stage-specific RNA-seq*

We collected four developmental stages of *D. gyrociliatus*: early embryo (1–3 days old), late embryo (4–6 days old), juvenile (7–9 days old), and adult (20–23 days old), with two biological replicates per sample. For each replicate, we collected and transferred a batch of adult *D. gyrociliatus* females from the main culture to a clean petri dish and let them lay eggs for three days. On the third day the adult females were removed from the dish. We collected and snap froze early and late embryos on the third and sixth day of development, respectively, the hatchling juveniles around the ninth day, and finally the adults twenty-three days after egg lay. Total RNA was extracted as described above and sequenced at GeneCore (EMBL) on a single Illumina lane using the NextSeq 500 High with 75 bp paired end reads and ~270 bp mean insert size.

To quantify transcript abundances, we pseudo-aligned the stage-specific RNA-Seq reads to *D. gyrociliatus* filtered gene models with Kallisto v0.44.0^96^, using the *--genomebam* parameter to project the transcript alignments to genomic coordinates, and *--bias* option to correct sequence-specific biases. We imported Kallisto estimated counts, calculated size factors, and estimated the data dispersion using the standard workflow of the DESeq2^97^, and performed a gene-level differential expression analysis to compare “late embryo vs early embryo”, “juvenile vs late embryo”, and “adult vs juvenile”. Datasets were first corrected for low count and high dispersion values using the *apeglm* log fold change shrinkage estimator^98^, and then compared using Wald tests between contrasts. Considering gene expression changes significant if the adjusted p-value < 0.05, we identified 8,341 differentially expressed genes (4,543 up and 3,798 down) for “late embryo vs early embryo”; 1,870 genes (938 up and 932 down) for “juvenile vs late embryo”; and 3,746 genes (1,827 up and 1,919 down) for “adult vs juvenile”.

For clustering and visualization (i.e., principal component analysis and heatmaps), we homogenized the variance across expression ranks by applying a variance stabilizing transformation to the DESeq2 datasets. We used the *pheatmap* package to create heatmaps^99^, the package *EnhancedVolcano* for volcano plots^100^, and *ggplot2* for the remaining plots^101^. To characterize and identify enriched gene ontology terms, we used the package *clusterProfiler*^102^. All analyses performed in R^103^ using the RStudio Desktop^104^.

4. **Phylogenetic analysis**

Annelid transcriptomes (Supplementary Data file 1) were downloaded from SRA and assembled using Trinity (v2.5.1)^105^ with the Trimmomatic^75^ read trimming option activated. Transcriptomes were then translated using Transdecoder (v5.0.2)^105^ after searching for similarity against the metazoan Swissprot database. Predicted proteins were searched using HMMER^106^ for 1,148 single-copy phylogenetic markers previously described^107^ using reciprocal BLAST to discard possible paralogues. Sequences for each marker were aligned with MSAProbs (v0.9.7)^108^ and trimmed to remove mis-translated segments using HMMClean2 (kindly provided by Hervé Philippe). Misaligned domains were pruned with BGME^109^. From this initial dataset, we selected the 264 genes with lowest saturation, yielding a concatenate alignment of 71,508 positions (as the analysis of the full dataset with site-heterogeneous models was not computationally tractable). Phylogenetic analyses were performed on the concatenated alignment using IQTREE^110^ with a C60 mixture model, a LG matrix to account for transition rates within each profile, the FreeRate heterogeneity model (R4) to describe across sites evolution rates and an optimisation of amino acid frequencies using maximum likelihood (FO). Support values were drawn from 1,000 ultrafast bootstraps with NNI optimisation. We also carried out Bayesian reconstruction using a site-heterogeneous CAT+GTR+Gamma model running two chains for more than a thousand iterations. We reached reasonable convergence for one of the datasets (200119-trfl.bpdiff, bpdiff > 0.19).

5. **Annotation of repeats and transposable elements (TEs)**

We used a combination of approaches to annotate repeats in the *D. gyrociliatus* genome. We constructed a *de novo* repeat library with RepeatModeler v.1.0.4^111^, which was used with RepeatMasker “open-4.0”^112^ to generate an automated annotation that identified a small fraction of the genome (11.13%) as derived from Transposable Elements (TEs) and repeats (Supplementary Table 3). The vast majority of annotated TEs correspond to DNA transposons, LTR retrotransposons and Penelope-like Elements. Among repeats, a small fraction of the genome (1.3%) corresponds to protein domains unrelated to TEs, possibly linked to the recent expansion of few gene families. The general feature format (gff) file with the automatic annotation of TEs and repetitive sequences is in Supplementary Data File 2.

In order to uncover low copy repeats that could have escaped the automated annotation, we performed a BLAST analysis using the TE sequences recovered with RepeatModeler and PFAM sequence collections corresponding to entries RVT_1 (PF00078) (Supplementary Table 4). This approach permitted to identify non-LTR retrotransposons and Helitrons, represented by only few copies. Using MITE Digger^113^, we identified MITEs whose Terminal Inverted Repeats (TIRs) matched *Mariner* transposons in the *D. gyrociliatus* genome. The rRNA, snRNA and tRNA genes identified with BLAST and tRNA-scan^114^ represent less than 0.1 % of the genome. The remaining portion of repeats (1.7 % of the genome) could not be identified and correspond to mostly non-coding sequences.

We used structural and phylogenetic criteria to classify *D. gyrociliatus* TEs. The DNA transposons belong to *Mariner* and *Mutator-Like Elements* (*MULE*), based on the amino-acid signature of their transposases. To establish the gene arrangement in LTR retrotransposons, we performed 6-frame translations of most intact copies, identified as such by having two identical LTRs and being flanked by short direct repeats created by Target Site Duplication (TSD). LTR retrotransposons were further compared to other elements of the *Ty3/gypsy* clade, using a collection of protein sequences comprising the reverse transcriptase domain and the integrase core domain. The phylogeny of *Ty3/gypsy* was established with a collection of sequences that includes well-annotated elements described in the Gypsy database^115^. It also includes hits obtained with TBLASTN (databases NR and TSA) using *D. gyrociliatus* sequences as queries.

To look for distant homologues of the protein found downstream the integrase in LTR retrotransposons, we submitted a multiple sequence alignment of ten peptide sequences (corrected to the original coding frame when recovered from disrupted genes) to HHPred (database PDB_mmCIF70_28_Dec). Using MODELLER, the three best hits (Probability > 99, E-value < e^-29^) were used to model the 3D structure of the *Dingle-1* envelope.

6. **Gene prediction and functional annotation**
   6.1. *Gene prediction*

To annotate protein-coding genes, the predicted set of 246 core eukaryotic genes generated by CEGMA^116^ was first used to train AUGUSTUS v.3.2.1^117^ to generate a set of *ab initio* predicted proteins. Thereafter, the predicted proteomes of the annelids *Capitella teleta* and *Helobdella robusta* were aligned to the *D. gyrociliatus* genome using EXONERATE v.2.2.0 algorithm^118^, and PASA v.2.0.2^119^ was used to align the transcriptome to the genome with BLAT and GMAP aligners^82,120^. These three independent sources were inputted into EvidenceModeler v.1.1.1^121^ to generate weighted consensus gene predictions. *Ab initio* gene predictions and spliced protein alignments were given a weight of 1, whereas PASA transcript assemblies were weighted 10. EvidenceModeler output was used to refine PASA gene models and generate alternative splice variants. This initial set of models was filtered to remove predictions with BLAST hit against transposons and/or with an overlap equal or greater than 90% on masked regions. The final prediction set contains 14,203 coding-protein loci that generate 17,409 different peptides. Additionally, 1,787 of these genes have evidence of alternative splicing as identified by PASA. Finally, we used ORFik^122^ to refine transcription start sites (TSS) with Cap Analyses Gene Expression (CAGE)-seq data. The gene transfer format (gtf) file with all filtered gene models is available in Supplementary Data file 3.

6.2. *Functional annotation*

Protein homologies for the 17,409 different gene models were annotated with BLAST v.2.2.31+^123^ on the UniProt/SwissProt database provided by Trinotate v.3.0. We used HMMER v.2.3.2^124^ to identify protein domains using the PFAM-A database provided by Trinotate v.3.0. Signal peptides and transmembrane regions were predicted with signalP v.4.1^125^ and tmHMM v.2.0c^126^ respectively. These annotations were integrated into a database with Trinotate v.3.0, which retrieved Gene Onthology (GO)^127^, eggNOG^128^ and KEGG^129^ terms for each gene model. We retrieved a functional annotation for 13,437 gene models (77.18%). The functional annotation report is provided in Supplementary Data file 4.

7. **Gene structure evolution analysis**
   7.1. *Genome-wide gene structure analysis*

To identify evolutionary changes in gene structure, we compared genome-wide values of gene structure parameters (mRNA length, intergenic region length, 5’ UTR length, 3’ UTR length, exon length, and exon number per gene) between genomes of *D. gyrociliatus* and the annelids *C. teleta* and *H. robusta*, as well as *D. melanogaster*, *C. elegans*, and *O. dioica* genomes (the three best sequenced compact bilaterian genomes). For *C. teleta*, *H. robusta*, *D. melanogaster*, and *C. elegans*, we used the genome assemblies and gff3 annotation files in Ensembl Metazoa on 17.02.17. For *O. dioica*, we used the genome assembly and gff3 annotation file in OikoBase^130^ on 15.02.17.

7.2. *Trans-splicing*

To identify spliced leader sequences in *D. gyrociliatus*, we predicted protein coding sequences in the *de novo* assembled transcriptome with Transdecoder v.5.5.0^105^ and used the scripts *nr_ORFs_gff3.pl* (from Transdecoder) and *gff3_file_UTR_seq_extractor.pl* (from PASA) to extract the non-redundant 5’ UTR sequences of protein coding transcripts. We used these sequences and Jellyfish v.2.2.3^88^ to identify over-represented 22-mer and 50-mer sequences that would correspond to the spliced leader.

7.3. *Intron evolution analysis*

For a first focused comparative analysis of intron features, six species (plus *D. gyrociliatus*) were selected due to their phylogenetic position and known paucity of intron evolution. Genome annotations were downloaded from Ensembl (*Homo sapiens*, version 88), Ensembl Metazoa (*Capitella teleta* v1.0 version 40; *Crassostrea gigas* v9 version 40; *Lottia gigantea* Lotgi1 version 40; *Strigamia maritima* Smar1 version 26) and from the European Amphioxus Genome webpage (https://amphiencode.github.io/Data/; *Branchiostoma lanceolatum* braLan2). To compare distributions of intron lengths among these species (Fig. 2d), only introns in genes with orthologs across the seven species (as defined by OrthoFinder; see section 8.1) were used. If any of the species had five or more paralogs in the same orthology group, the whole orthogroup was discarded.

Next, to identify *D. gyrociliatus* introns that are conserved with the other species (and thus ancestral) or, on the contrary, that are not shared with any of the other species (and thus likely originated after the split of the *D. gyrociliatus* linage) (Extended Data Figure 4h), we performed the following steps. First, we aligned each *D. gyrociliatus* protein against each annotated protein isoform of each orthologous gene of the abovementioned six species and added the intron positions into the alignments^131^. To identify high-confidence conserved intron positions, we required that a given *D. gyrociliatus* intron position is found at the exact position of the alignment and with the same phase (0, 1 or 2) in at least four out of six other species. To define high-confidence non-conserved (likely novel) introns, we required that a *D. gyrociliatus* intron position did not match an intron position with the same phase within 25 alignment positions in any of the other six species.

Next, we performed a quantitative analysis of intron gain and loss in *D. gyrociliatus* using a database of homologous introns from 28 metazoan genomes (Supplementary Table 7; the annelid *H. elegans* was excluded in this analysis). To do so, we first obtained a list of one-to-one orthologous genes from each species using BUSCO v3^81^ and the OrthoDB v9^132^ dataset of 978 single-copy animal orthologs. BUSCO was run using predicted peptides as input (prot mode) and a maximum expected value of 1e-04 (evalue). For each genome, we then retrieved the list of single-copy genes (excluding duplicates and fragmented genes as defined by BUSCO) and the coordinates of each of their introns in their CDS sequence, obtained from GFF annotation files using the gffread utility (-x flag to extract exons and coordinates). Then, we aligned the predicted peptides using MAFFT v7.310 G-INS-i algorithm^133^. The alignments and CDS intron coordinates were provided to Malin^134^, a software utility that implements a probabilistic model of intron evolution and that we used to identify conserved intron sites across all 28 genomes and infer their conservation status in ancestral nodes. Specifically, we used 18,270 conserved intron sites from all 978 alignments, defined as introns that shared phase, were unambiguously aligned in ≥25% of the orthologs and had at least 5 non-gapped aligned positions at both sides^135^. Using this table of intron presence per extant genome, we estimated the rates of intron gain and loss in each node with Malin’s built-in model maximum-likelihood optimization procedure. The initial model of intron evolution, which assumes equiprobable intron gain and loss rates, was optimized in 1000 optimization iterations (likelihood convergence threshold = 0.001) to obtain node-specific intron gain and loss rates (Supplementary Data File 5). Then, we used this model to estimate the posterior probabilities of intron presence, gain and loss in extant and ancestral nodes.

For each node, we calculated the intron density expressed as introns per kbp of coding sequence (introns/CDS kbp), as follows:

*intron density_i_ = ((num introns_i_ / num genes) / median gene length) * 1000*

where *num introns_i_* is the number of introns present in a given node (extant or ancestral, corrected by missing sites), *num genes* = 978 (number of alignments of one-to-one orthologs) and *median gene length* = 682.5 bp (as obtained from the lengths of the seed proteins curated in the OrthoDB v9 Metazoa data-set; ‘ancestral’ FASTA file). We used the same procedure to obtain the rates of intron gain and loss per node in terms of introns/CDS kbp. To visualize the evolution of intron content, we plotted the intron densities, gains and losses along the animal phylogeny using the *ape* library v5.0^136^ from the R statistical package v3.6 (R Core Team, 2017). In addition, we inferred the uncertainty of the estimated intron gains, losses, and presence values. Specifically, we used Malin to compute 1000 bootstrap iterations with random selection of intron sites from the original intron presence table, with replacement. A complete report of the estimated intron densities (with bootstrapped confidence intervals), gains and losses is available as Supplementary Data File 5.

Finally, to assess the impact of intron length on splicing efficiency on *D. gyrociliatus*, *S. maritima* and *H. sapiens* (Extended Data Figure 4i), we used RNA-seq-based quantifications of intron retention as previous described^137^ and implemented by *vast-tools*^138^. In brief, a sequence library of junctions was generated for all introns, including each exon-intron, intron-exon and exon-exon junction in the genome. RNA-seq data were mapped to these junctions and percent of intron retention (PIR) was calculated for each annotated intron as PIR = 100 * av(#Reads_exon-intron_, #Reads_intron-exon_)/[av(#Reads_exon-intron_, #Reads_intron-exon_) + #Reads_exon-exon_]. For further information about library generation, RNA-seq mapping, read number correction by mappability and imbalance filters, see REF^137^. To calculate the average PIR for each intron, we used 108 RNA-seq samples for *H. sapiens* (collected from VastDB^138^) nine for *S. maritima*^139^ and the nine samples generated in this study for *D. gyrociliatus*. Only introns that had sufficient read coverage (LOW or higher, as defined by *vast-tools*^138^ in at least three samples and with no significant read imbalance between the two exon-intron junctions (as assessed by a binomial test^137^) were used to calculate average PIR.

8. **Gene family evolution analyses**
   8.1. *Construction of orthology groups*

We used OrthoFinder v.2.2.7^140^ with default values to reconstruct gene family assignments between *D. gyrociliatus* non-redundant proteins and 27 other animal non-redundant proteomes (Supplementary Table 6).

8.2. *Gene family gains, losses, and expansions*

OrthoFinder gene families were parsed and mapped onto a reference species phylogeny to infer gene family gains and losses at different nodes using the ETE 3 library^141^. Gene expansions were computed for each species using a hypergeometric test against the median gene number per species for a given family.

8.3. *Evolution of G-coupled protein receptor family GPCRs and neuropeptides*

GPCR sequences were retrieved using HMMER v3.2.1^106,124^ (e-value cutoff < 0.01) with Pfam profiles of class A (PF00001), class B (PF00002), class C (PF00003) and class F (PF01534) GPCRs (according to GRAFS classification). Identical protein sequences from the protein prediction were reduced to a single sequence. Sequences from each class were tested for false positives from other classes (including cAMP slime-mold class E GPCRs, PF05462). The estimated number of different GPCR types (2^nd^ number in Supplementary Table 8) was determined based on BLAST similarities of p > 1e-150, using the Network based cluster search of the CLANS software^142^, with offset values and minimum sequences per cluster set to “1”. The similarity value was chosen after testing the *Drosophila* class A GPCRs to distinguish between the two *D. melanogaster* CCHamide receptor paralogs as well as between the *D. melanogaster* octopamine beta 1, 2 and 3 receptors. For the phylogenetic analysis, sequences were aligned with Clustal X^143^ using standard values, trimmed with trimAl^144^ using -automated1 option, analyzed with FastTree 2^145^ using -lg model and visualized with FigTree. To characterize the GPCR content of other annelids and bilaterians, we used the genome annotations reported in Supplementary Table 7. Neuropeptide candidates were retrieved by a combination of BLAST-searches (e-value cutoff < 0.1) and the use of a customized script to detect cleavage patterns on precursors, as described elsewhere^146^. Neuropeptide precursors are reported in Supplementary Data file 6.

8.4. *Identification of transcription factors and developmental pathways*

We used the functionally annotated gene predictions of *D. gyrociliatus*, and those generated with Trinotate from the genomic annotations of *C. teleta* and *H. robusta* (Supplementary Data files 7 and 8) to identify the repertoires of transcription factors, ligands and receptors present in these three annelids. Searches were conducted by BLAST annotation and PFAM ID, as shown in Extended Data Figure 5g, h. If a gene was not in the annotated genome, and before claiming a gene loss, we performed manual search via BLAST searches on the *de novo* and genome guided transcriptome. For *T. axi* and *D. vorticoides*, gene identification was conducted on the assembled transcriptome via manual BLAST searches.

8.5. *KEGG metabolic pathway mapping*

We used the functionally annotated gene predictions of *D. gyrociliatus*, *C. teleta* and *H. robusta* obtained from Trinotate to extract KEGG IDs, which were thereafter mapped into pathways with KEGG Mapper^147^.

8.6. *Orthology assignment*

Multiple protein alignments were constructed with MAFFT v.7^133^, poorly aligned regions were either removed by hand or with gBlocks^148^. Maximum likelihood trees were constructed with with FastTree 2^145^ with default parameters and visualized with FigTree (https://github.com/rambaut/figtree/).

9. **Gene expression analyses in *D. gyrociliatus***
   9.1. *Gene isolation and riboprobe synthesis*

Sequences of the nine Hox genes were identified in the transcriptome of *D. gyrociliatus* by TBLASTX and compared against those annotated in the genome. We designed gene-specific primers to cover the coding region of each Hox gene with MacVector v.11.0.4 and amplified these fragments from cDNA libraries obtained from mixed embryonic stages, juveniles, and adults. PCR products were cloned into pGEM-T Easy vectors (Promega, USA) and sequence-verified by Sanger sequencing. Before whole mount *in situ* hybridization, antisense riboprobes were synthetized with MEGAscript kit T7 or Sp6 (Ambion) and labelled with digoxigenin-11-UTP (Roche, USA), following manufacturer’s recommendations.

9.2. *In situ hybridisation*

Adults and juveniles were relaxed in 7.8% MgCl_2_ before fixation in 4% paraformaldehyde in MgCl_2_ solution for 1 hour at room temperature (RT). Embryos were collected in their egg clusters and manually dissected. Before fixation, the embryonic eggshell was digested in a solution of 1% sodium thioglycolate (Sigma-Aldrich, T0632,) and 0.05% protease (Sigma-Aldrich, P5147) in sea water, pH 8, for 30 min at RT, followed by relaxation in MgCl_2_ and fixation. All samples were stored in 100% methanol at -20 °C. Whole mount *in situ* hybridization was performed as described elsewhere^149^. Briefly, rehydrated samples were digested with proteinase K for 10 min and hybridised with DIG-labelled riboprobes at a concentration of 1ng/μl, at 62°C for approximately 72 hours. After incubation with anti-DIG-AP antibody at a 1:5000 concentration, the signal was visualized with nitroblue tetrazolium chloride and 5-bromo-4-chloro-3-indolyl phosphate (NBT/BCIP). After noise removal with graded ethanol washes, the samples were stored in 80% glycerol. The nuclear marker DAPI (Sigma-Aldrich) at a concentration of 1:5000 was added to the medium to stain the nuclei and allow the identification of the developmental stages.

9.3. *Imaging*

Images were taken with a Zeiss Axiocam HRc connected to a Zeiss Axioscope Ax10 using bright-field Nomarski optics. Images were then processed with Adobe Photoshop CS6, to adjust the size and the contrast when needed.

10. **Gene expression analyses in *C. teleta***

Adults of *C. teleta*^150^ were cultured in glass finger bowls with artificial sea water (ASW) and fed with sieved mud as described before^151,152^. Embryos and larvae were obtained by dissecting the brooding tubes from females. Fixation and whole-mount *in situ* hybridisation (WMISH) was performed as previously described^153^. *Capitella teleta* orthologs *Ct-fgf8/17/18/24* (Protein ID: 218971), *Ct-pvf1* (Protein ID: 153454) and *Ct-pvf2* (Protein ID: 220370) were mined from the publicly available genome^48^. Gene orthology has been determined previously for *Ct- fgf8/17/18/24*^154^, and *Ct-pfv1* and *Ct-pvf2*^155^. Fragments of the coding sequences of these genes were amplified to use as templates for RNA probes. Gene-specific primers with a T7 adapter on the 5’ end of the reverse primer (underlined) were used as follows: *Ct- fgf8/17/18/24*: 1061 bp containing the FGF domain, Fwd primer 5’-GTAACCTGCCATCCGTTGAC and Rev primer 5’-GCCCCGGCCAGAGAAGGGCGAACTGTTG; *Ct-pvf1*: 1009 bp containing the PDGF domain, Fwd primer 5’-CTTCGCAGTGAGAGAGGCTA and Rev primer 5’-GCCCCGGCGGATCACTTGCTTGGACGC; and *Ct-pvf2*: 738 bp containing the PDGF domain, Fwd primer 5’-TCAGCCCAAAGATGTAACGG and Rev primer 5’-GCCCCGGCGAGAGACGGGTGTTGGGG. Animals were hybridised for 72 h at 65 °C with 1 ng/μL of DIG-labelled anti-sense RNA probes. WMIHS animals were mounted in 70% glycerol in PBS with 1:5000 4’,6-diamidino-2-phenylindole (DAPI, Invitrogen, cat#D3571) for nuclear staining and imaged on a Leica DMRA2 compound microscope coupled with a QIClick camera. Animals stained for F-actin were fixed 30 min at room temperature, incubated with 1:100 BODIPY FL-Phallacidin (Life Technologies, cat #B607; stock concentration 200 Units/mL in methanol), and imaged with an IXplore SpinSR (Olympus). WMISH images were rendered using Helicon Focus (HelSoft). Contrast and brightness were edited with Photoshop CC version 14.0 (Adobe Systems, Inc.) and figure panels were constructed using Illustrator CC version 17.0.0 (Adobe Systems, Inc.).

11. **Macrosynteny analysis**

Single-copy orthologues obtained using the mutual best hit approach (MBH) were used to generate Oxford synteny plots comparing sequentially indexed orthologue positions. Plotting order was determine by hierarchical clustering of the shared orthologue content using the complete linkage method as originally proposed^156^.

12. **Assay for Transposase-Accessible Chromatin using sequencing (ATAC-seq)**
    12.1. *Tagmentation and library preparation*

The Assay of Transposase-Accessible Chromatin with high-throughput sequencing (ATAC-seq) was performed as described elsewhere^157^, using 50,000-70,000 cells (approx. 18 adult females) and home-brewed Tn5 transposase^158^. Cell dissociation and lysis was improved by disaggregating the tissue with a syringe in lysis buffer. Transposed DNA fragments were amplified by 16 cycles of PCR. Two replicate Illumina libraries were sequenced in a NextSeq500, in rapid paired end mode and 75 base pairs of read length.

12.2. *Bioinformatic analysis*

Adapter contamination was removed with CutAdapt v.1.2.1^71^ and the cleaned reads were aligned to the unmasked genome with a script kindly provided by Dr Juan Tena. Peaks were called with MACS2 v.2.1.1.20160309^159^ with the options --nomodel --extsize 70 --shift -30 --call-summits --keep-dup 1. Irreproducible discovery rates (IDR) were calculated with the R scripts of IDRCode^160^. A final set of 10,241 consistent peaks (IDR ≤ 0.05) between the two replicates was used for further analyses (Supplementary Data file 9). *De novo* motif enrichment analysis of the ATAC-defined regulatory sites was performed using HOMER^161^, with default parameters, except -size given (Supplementary Data file 10).

13. **Cap Analysis Gene Expression (CAGE)-seq**
    13.1. *CAGE library preparation*

Total RNA from adult *D. gyrociliatus* was isolated using Trizol followed by RNeasy RNA clean-up protocol (Qiagen). CAGE libraries were prepared for two biological replicates (barcodes ATG and TAC) using the latest nAnT-iCAGE protocol^162^. This protocol allows for the most unbiased CAGE library preparation as it omits the restriction enzyme treatment and PCR amplification. The libraries were sequenced in single-end 50 bp mode (Genomic Facility, MRC LMS).

13.2. *Processing of CAGE tags*

Sequenced libraries were demultiplexed with no mismatches allowed in barcode identification. Demultiplexed CAGE reads (47 bp) were mapped to the *D. gyrociliatus* genome assembly using Bowtie2^163^ with zero mismatches allowed in the 22 nucleotide seed sequence. Mapped reads were sorted, and only uniquely mapped reads were kept for downstream analysis in R graphical and statistical computing environment^103,104^. Bam files were imported into R using the standard CAGEr package (version 1.20.0) workflow^164^. Within CAGEr, 5’G that is commonly added by reverse transcriptase, was removed using the following G-correction workflow: 1) if the first nucleotide is a mismatching G, it is removed; 2) if the first nucleotide is a G and not a mismatch, its removal is decided according to percentage of identified mismatched G’s. Normalization was performed using a referent power-law distribution^165^ (Supplementary Data files 11–14).

CTSSs that passed the threshold of 1 TPM were clustered together using distance-based clustering (CAGEr package) with 20 bp allowed as a maximum distance between neighboring CTSSs (Supplementary Data file 15). We refer to these CTSS clusters as tag clusters and they represent a single functional transcription unit. Broadness of each cluster was calculated as interquantile width spanning 10^th^ and 90^th^ percentile of the signal to exclude extreme outlier CTSSs. Genomic locations of tag clusters were determined using the ChIPseeker package and gene model annotations, where promoters were defined to include 500 bp upstream and 100 bp downstream of the annotated transcript start site. Visualization of motifs, sequence patterns, or reads coverage was performed using Heatmaps and seqPattern Bioconductor packages.

**Supplementary Figures**

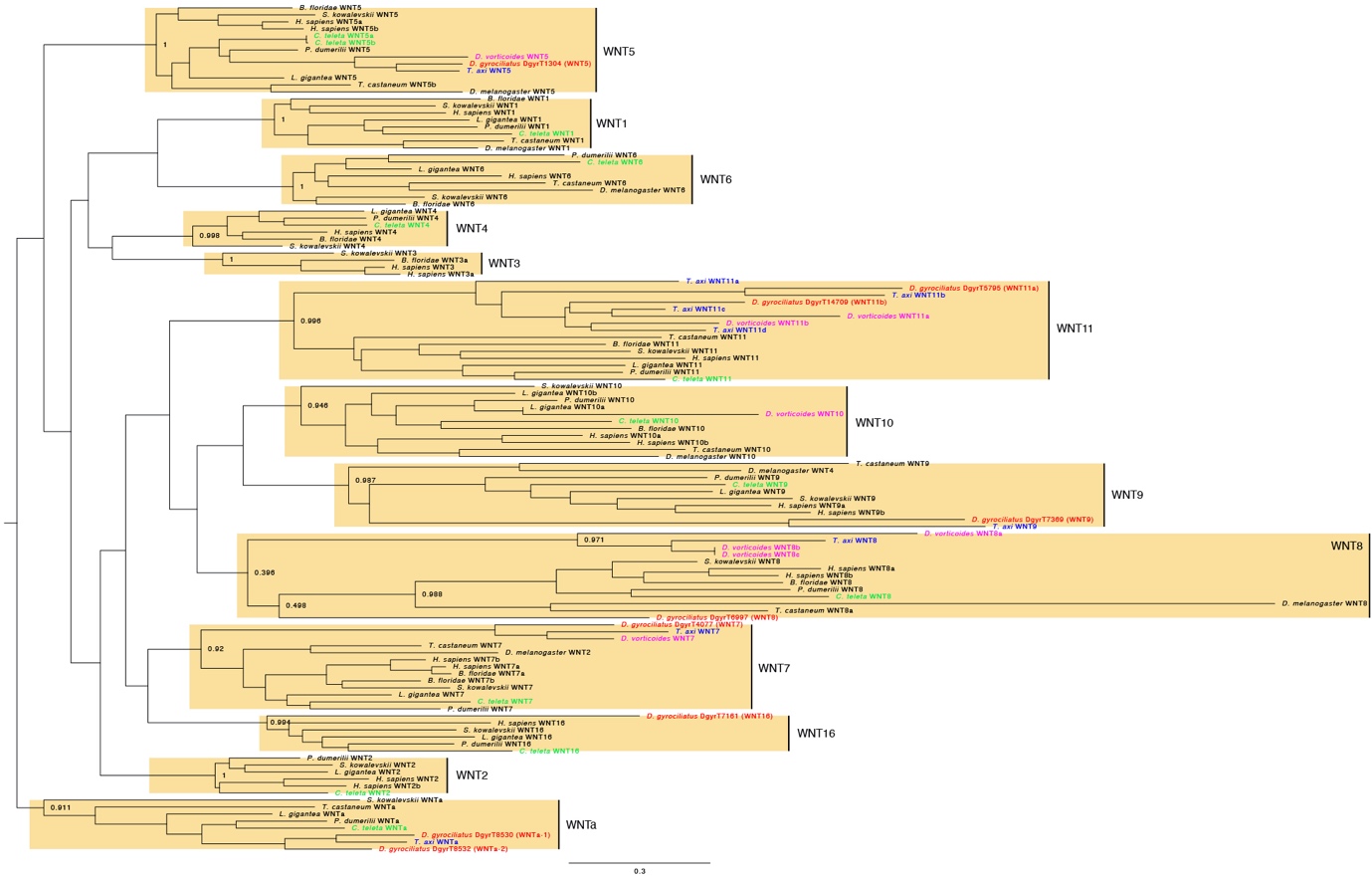

**Supplementary Figure 1 | Wnt ligand orthology assignment.** Maximum likelihood tree of Wnt ligands, including those putative sequences identified in *D. gyrociliatus* (in red), *T. axi* (in blue), *D. vorticoides* (in pink) and *C. teleta* (in green). Each major orthology group is highlighted with a light-yellow box, and only bootstrap values supporting these clades are depicted.

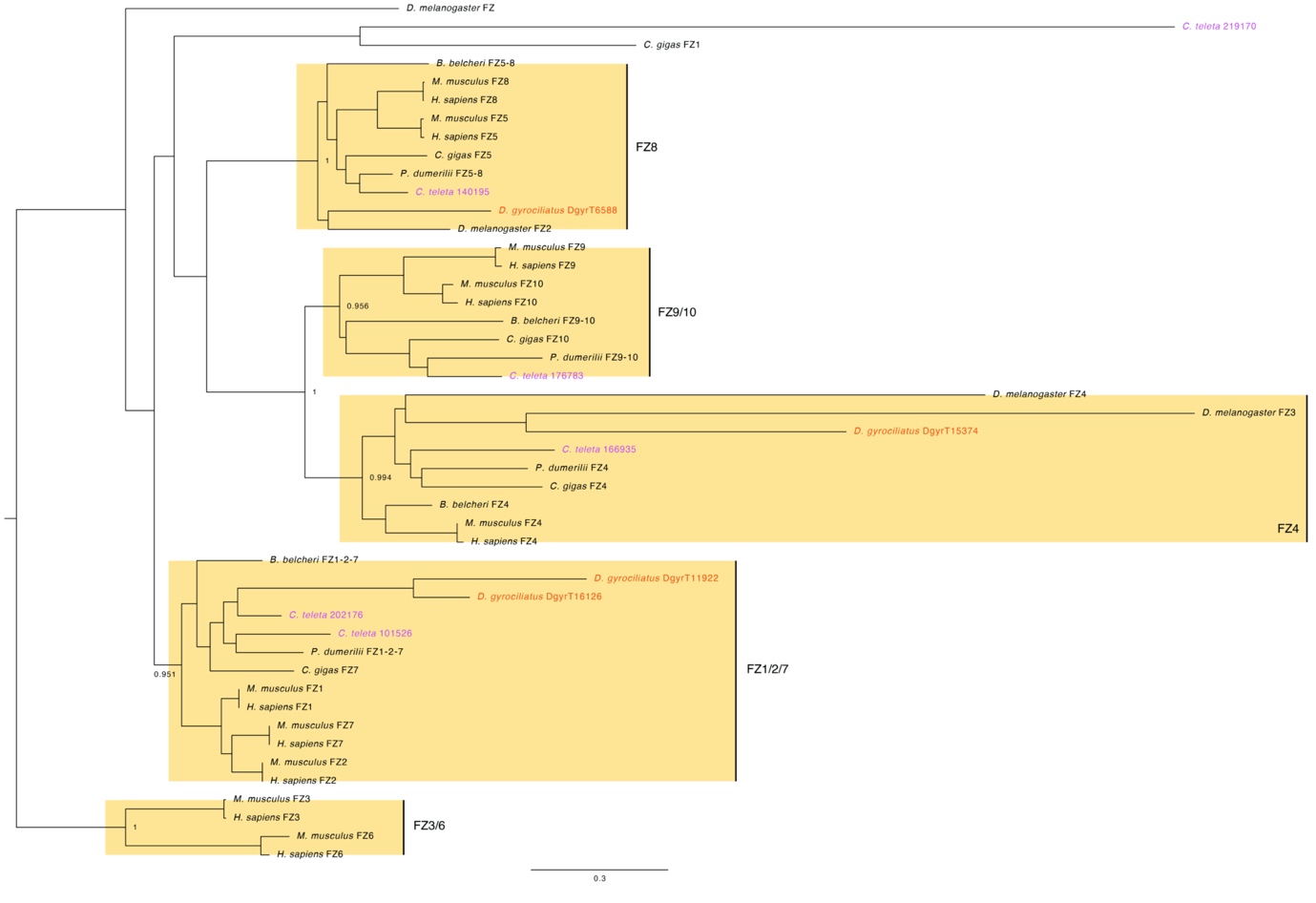

**Supplementary Figure 2 | Frizzled receptor orthology assignment.** Maximum likelihood tree of Frizzled receptors, including those putative sequences identified in *D. gyrociliatus* (in red) and *C. teleta* (in violet). Each major orthology group is highlighted with a light-yellow box, and only bootstrap values supporting these clades are depicted.

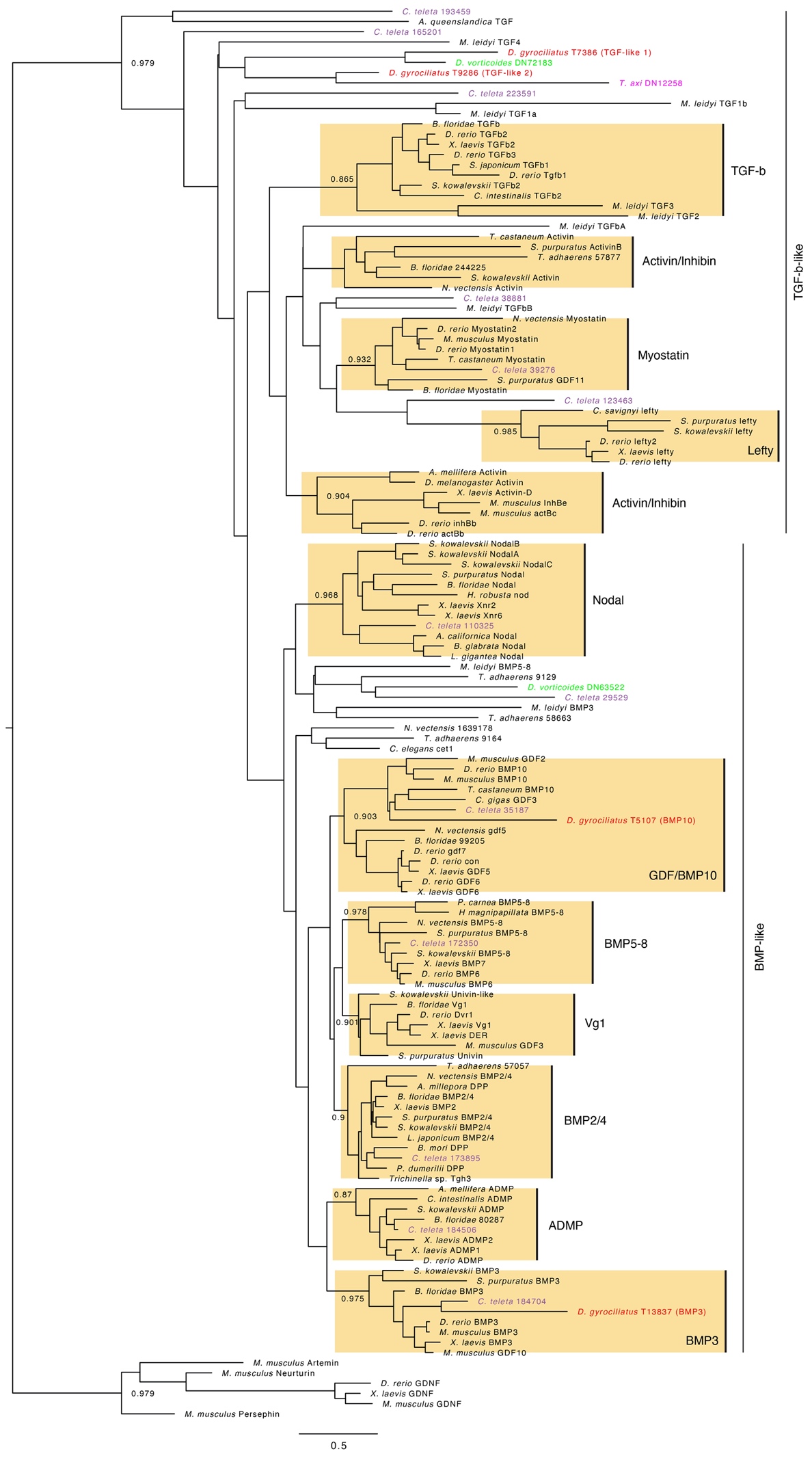

**Supplementary Figure 3 | TGF-β ligand orthology assignment.** Maximum likelihood tree of TGF-β ligands, including those putative sequences identified in *D. gyrociliatus* (in red), *T. axi* (in pink), *D. vorticoides* (in green) and *C. teleta* (in violet). Each major orthology group is highlighted with a light-yellow box, and only bootstrap values supporting these clades are shown.

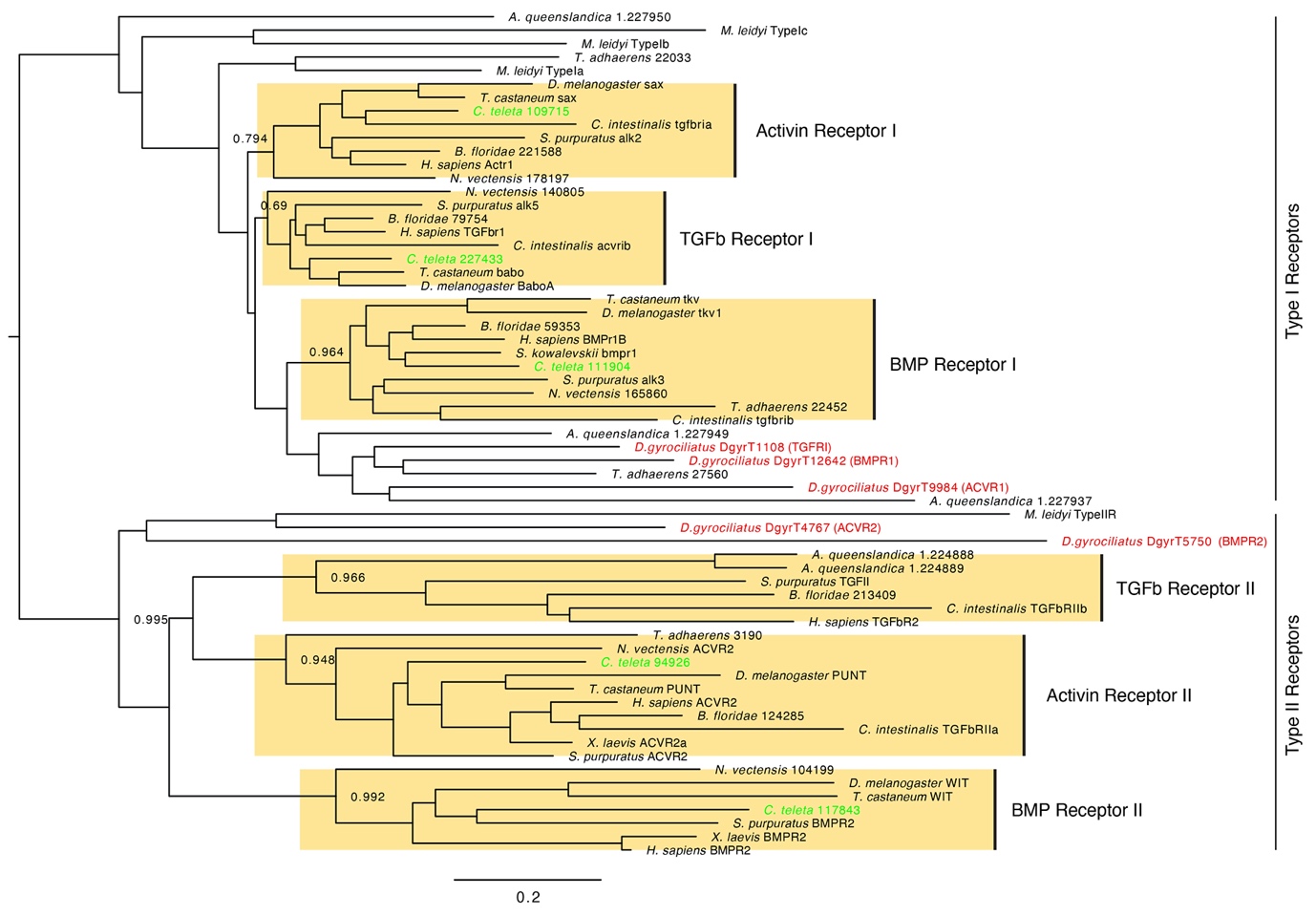
**Supplementary Figure 4 | TGF-β receptor orthology assignment.** Maximum likelihood tree of TGF-β receptors, including those putative sequences identified in *D. gyrociliatus* (in red) and *C. teleta* (in green). Each major orthology group is highlighted with a light-yellow box, and only bootstrap values supporting these clades are shown.

**Supplementary Tables**

**Supplementary Table 1 | Genome assembly statistics.**

|  | *D. gyrociliatus* | *T. axi* | *C. teleta^a^* | *H. robusta*^a^ | *H. elegans^b^* | *E. fetida^c^* | *S. lamarcki^d^* | *L. luymesi^e^* |
| --- | --- | --- | --- | --- | --- | --- | --- | --- |
| Genome size (Mbp) | 77.9 | 120.2 | 333.72 | 235.34 | 1,026.05 | 1,052.63 | 1,501.63 | 687.7 |
| GC (%) | 31.36 | 26.77 | 40.39 | 32.82 | 35.43 | 40.87 | 34.78 | 40.16 |
| # of scaffolds | 349 | 169,751 | 21,042 | 1,993 | 188,407 | 1,659,527 | 2,474,717 | 11,871 |
| # of scaffolds >1 kb | 347 | 28,366 | 21,042 | 1,993 | 101,980 | 268,873 | 355,424 | NA |
| Largest scaffold (Mb) | 5.53 | 0.032 | 1.62 | 13.64 | 0.244 | 0.059 | 0.053 | 2.12 |
| Scaffold N50 (Mb) | 2.24 | 0.002 | 0.19 | 3.06 | 0.018 | 0.003 | 0.002 | 0.373 |
| # N’s per 100 kb | 1,170.41 | 1,117.84 | 16,991.19 | 8,471.67 | 2,551.11 | 34,398.06 | 4,959.94 | NA |

^a^REF^48^; ^b^REF^166^; ^c^REF^167^; ^d^REF^168^; ^e^REF^169^

**Supplementary Table 2 | Transcriptome assembly statistics.**

|  | *de novo* | genome-guided |
| --- | --- | --- |
| Total bases | 200,773,339 | 196,572,502 |
| Transcripts | 87,192 | 74,131 |
| Unique genes | 49,945 | 39,432 |
| Transcripts with ORF | 44,889 | 42,041 |
| Largest | 34,371 | 30,551 |
| Mean transcript length | 2,303 | 2,652 |
| Transcripts over 10kb | 2,301 | 2,302 |
| Transcripts over 1kb | 47,905 | 45,625 |
| N50 | 4,544 | 4,934 |
| E90N50 | 4,017 | 4,229 |
| GC | 0.34 | 0.34 |
| BUSCO (%) | 96.4 | 95.8 |

**Supplementary Table 3 | Automated annotation of repeats and transposable elements in annelid genomes.**

|  | *D. gyrociliatus* | *Capitella teleta^a^* | *Helobdella robusta*^a^ |
| --- | --- | --- | --- |
| Interspersed repeats^b^ | 7.95% (6.19 Mb) | 22.87% (76.39 Mb) | 18.46% (43.38 Mb) |
| Simple repeats | 2.69% (2.09 Mb) | 3.68% (12.3 Mb) | 12.21% (28.69 Mb) |
| Low complexity | 0.55% (0.43 Mb) | 4.24% (14.15 Mb) | 2.58% (6.07 Mb) |
| Total masked | 11.18% (8.71 Mb) | 31.06% (103 Mb) | 33.39% (78 Mb) |

^a^Percentages obtained from REF^48^.

^b^Including unclassified

**Supplementary Table 4 | Manually identified repeats and TEs in *Dimorphilus* genome.**

| Repeat class | Bases | % of assembly |
| --- | --- | --- |
| Gypsy-like | 1298162 | 1,67 |
| Other LTR | 178982 | 0,23 |
| *Total LTR* | *1477144* | *1,90* |
| Mariner | 667512 | 0,85 |
| MULE | 1167144 | 1,50 |
| Helitron | 51842 | 0,07 |
| *Total DNA* | *1886498* | *2,42* |
| Penelope-like | 340281 | 0,44 |
| non-LTR | 29302 | 0,04 |
| *Total non-LTR* | 369583 | 0,47 |
| MITEs | 60382 | 0,08 |
| *Total TEs* | *3793607* | *4,87* |
| rRNA | 39378 | 0,05 |
| snRNA | 5858 | 0,01 |
| tRNA | 26258 | 0,03 |
| Low complexity | 423366 | 0,54 |
| Simple repeats | 2059306 | 2,64 |
| Protein Domain-derived | 1034653 | 1,33 |
| Unknown | 1365378 | 1,75 |

**Supplementary Table 5 | Comparison of TE abundance and diversity in small animal genomes.**

| Genome | Assembly size, Mb | % of annotated TEs | TE superfamilies | Reference |
| --- | --- | --- | --- | --- |
| *Dimorphilus gyrociliatus* | 78 | 4.87 | 7 | This study |
| *Oikopleura dioica* | 72 | 14.10 | 9 | REF^170^ |
| *Fritillaria borealis* | 91 | 14.40 | 9 | REF^170^ |
| *Trichoplax adhaerens* | 98 | 0.13 | 16 | REF^171^ |
| *Polypedilum nubifer* | 107 | 1.26 | 14 | REF^172^ |
| *Polypedilum vanderplanki* | 104 | 0.26 | 6 | REF^172^ |
| *Ramazzottius varieornatus* | 55 | 18.12 | - | REF^35^ |
| *Belgica antarctica* | 99 | 0.12 |  | REF^173^ |
| *Caenorhabditis elegans* | 100 | 12–16 |  | REF^174^ |

**Supplementary Table 6 | List of species and gene annotation files used in this study**

| **Clade** | **Species** | **Source** | **Version** |
| --- | --- | --- | --- |
| Non-bilaterian | *Nematostella vectensis* | EnsemblMetazoa | ASM20922v1 |
|  | *Amphimedon queenslandica* | EnsemblMetazoa | Aqu1 |
|  | *Mnemiopsis leidyi* | EnsemblMetazoa | MneLei_Aug2011 |
| Deuterostomia | *Homo sapiens* | Ensembl | GRCh38 |
|  | *Lepisosteus oculatus* | Ensembl | LepOcu1 |
|  | *Oikopleura dioica* | OikoBase | reference v.1.0 |
|  | *Ciona intestinalis* | Ensembl | GCA_000224145.1 |
|  | *Branchiostoma lanceolatum* | Amphiencode | Bl71 |
|  | *Saccoglossus kowalevskii* | OIST | JGIv3.0 |
| Ecdysozoa | *Ramazzottius varieornatus* | Kumamushi GP | Downloaded 04/09/18 |
|  | *Strigamia maritima* | EnsemblMetazoa | Smar1 |
|  | *Drosophila melanogaster* | EnsemblMetazoa | BDGP6 |
|  | *Tribolium castaneum* | EnsemblMetazoa | Tcas5.2 |
|  | *Caenorhabditis elegans* | EnsemblMetazoa | WBcel235 |
|  | *Pristionchus pacificus* | EnsemblMetazoa | WS220 |
| Spiralia | *Capitella teleta* | EnsemblMetazoa | Capitella_teleta_v1.0 |
|  | *Helobdella robusta* | EnsemblMetazoa | Helro1 |
|  | *Dimorphilus gyrociliatus* | This study | Filtered gene models |
|  | *Hydroides elegans* | NCBI | GCA_001703475.1 |
|  | *Lottia gigantea* | EnsemblMetazoa | Lotgi1 |
|  | *Crassostrea gigas* | EnsemblMetazoa | oyster_v9 |
|  | *Octopus bimaculoides* | EnsemblMetazoa | PRJNA270931 |
|  | *Patinopecten yessoensis* | PyBase | GCF_002113885.1 |
|  | *Lingula anatina* | OIST | v2 |
|  | *Phoronis australis* | OIST | ASM263300v1 |
|  | *Notospermus geniculatus* | OIST | ASM263302v1 |
|  | *Macrostomum lignano* | WormBase | ML2 |
|  | *Schmidtea mediterranea* | smedDB | sexual v4 |
|  | *Adineta vaga* | EnsemblMetazoa | AMS_PRJEB1171_v1 |

**Supplementary Table 7 | Gene family statistics in annelid proteomes**

|  | *D. gyrociliatus* | *C. teleta* | *H. robusta* |
| --- | --- | --- | --- |
| # genes | 14204 | 32173 | 23432 |
| # genes in orthogroups | 12058 | 25271 | 16552 |
| # genes in species-specific orthogroups | 270 | 757 | 353 |
| # shared orthogroups containing species | 7277 | 9586 | 6388 |
| # species-specific orthogroups | 45 | 121 | 96 |
| Average shared orthogroup size | 1.63 | 2.59 | 2.57 |

**Supplementary Table 8 | Estimation of GPCRs in *D. gyrociliatus* and other annelid and bilaterian genomes^a^.**

|  | **Class A**  **(Rhodopsin)** | **Class B**  **(Secretin & Adhesion)** | **Class C**  **(Glutamate)** | **Class F**  **(Frizzled)** |
| --- | --- | --- | --- | --- |
| *D. gyrociliatus* | 211^b^ (184)^c^ [154]^d^ | 16 (12) [12] | 18 (18) [14] | 5 (5) [4] |
| *C. teleta* | 1058 (889) [343] | 40 (34) [22] | 51 (33) [23] | 5 (3) [5] |
| *H. robusta* | 229 (222) [165] | 30 (28) [18] | 11 (9) [10] | 5 (5) [4] |
| *L. anatina* | 492 (384) [197] | 48 (33) [29] | 44 (21) [20] | 6 (5) [5] |
| *L. gigantea* | 309 (296) [196] | 59 (52) [32] | 17 (12) [15] | 5 (3) [5] |
| *R. varieornatus* | 424 (363) [115] | 24 (24) [17] | 22 (22) [12] | 6 (6) [5] |
| *D. melanogaster* | 134 (73) [58] | 33 (14) [8] | 23 (9) [10] | 8 (4) [5] |
| *S. maritima* | 93 (88) [80] | 37 (31) [22] | 14 (11) [13] | 5 (5) [5] |
| *O. dioica* | 21 (21) | 11 (11) | 0 | 2 (2) |

^a^The numbers shown in this table can generally differ between other publications as different thresholds are used to search for GPCRs and different thresholds are used to determine the different types.

^b^Number of GPCR sequences in protein database, including all splice variants.

^c^(#): Number of different GPCR types using a similarity score of p < 1e-150.

^d^[#]: Number of different GPCR types based on monophyletic nodes. (Ignoring species-specific expansions and strongly dependent on taxon sampling)

**Supplementary Table 9 | DNA double-strand break machinery in *D. gyrociliatus***

| **Classical non-homologous end-joining** | | | **Homologous recombination** | |
| --- | --- | --- | --- | --- |
| Mammalian gene | *D. gyrociliatus* ortholog | | Mammalian gene | *D. gyrociliatus* ortholog |
| Ku70 | | DgyrG8781 | Rad50 | DgyrG4628 |
| Ku80 | | DgyrG13008 | Mre11 | DgyrG7523 |
| Artemis | | DgyrG1840 | Nbs1 | DgyrG9305 |
| DNAPKcs | | DgyrG1700 | RPA | DgyrG2509, DgyrG121 |
| Polλ | | DgyrG6710 | Rad51 | DgyrG9448 |
| Polµ | | DgyrG6571 | Rad52 | DgyrG9875 |
| TdT | | – | Rad54 | DgyrG1112, DgyrG6388 |
| Lig4 | | DgyrG11361 | Polδ | DgyrG11456, DgyrG3541 |
| XRCC4 | | DgyrG7932 | BLM | Dgyr_TRINITY_GG_2260_c3_g1 |
| XLF | | DgyrG4118 | Mus81 | DgyrG7297 |
|  | |  | TOP3 | DgyrG5027, DgyrG10546 |
| **Microhomology-mediated end joining** | | | Eme1 | DgyrG6301 |
| FEN1 | | DgyrG4375 | ATM | DgyrG10839 |
| MRE11 | | DgyrG7523 | RBBP8 | DgyrG9775 |
| NBS1 | | DgyrG9305 | BARD1 | DgyrG6393 |
| PARP1 | | DgyrG7212 | BRCA1 | Absent (present in *C. teleta*) |
| XRCC1 | | DgyrG10492 | BRIP1 | DgyrG8613 |
|  | |  | TOPBP1 | DgyrG3868, DgyrG9395 |
|  | |  | Abraxas | DgyrG9679 |
|  | |  | RAP80 | Absent (also in *C. teleta*) |
|  | |  | BABAM1 | DgyrG1474 |
|  | |  | BRE | DgyrG1708 |
|  | |  | BRCC36 | DgyrG7090 |
|  | |  | PALB2 | Absent (also in *C. teleta*) |
|  | |  | BRCA2 | DgyrG9400 |
|  | |  | DSS1 | DgyrG2531 |
|  | |  | SYCP3 | Absent (also in *C. teleta*) |
|  | |  | EXO1 | DgyrG4997 |

**Supplementary Notes**

**Supplementary Note 1: Background information on *Dimorphilus gyrociliatus***

The meiofaunal annelid family Dinophilidae Macalister, 1876, contains 18 species in three genera, which differ in their morphology and life cycle^16^. They are found worldwide in marine intertidal to subtidal waters, living interstitially in coarse sediments as well as epibenthically on biofilm and filamentous algae growing on macroalgae. All dinophilids are microscopic, 1–3 mm long worms, characterised by six body segments and dense midventral locomotory ciliation used for gliding. Their segments are not always externally discernible, and all species lack common annelid traits such as chaetae (chitinous bristles), parapodia (paired lateral segmental outgrows for locomotion), a blood vascular system and a (trochophore) larval stage. Furthermore, they possess protonephridia (not metanephridia) and are devoid of sensory, feeding, and respiratory appendages such as palps, antennae or cirri on both head and trunk^175^.

Whereas the monophyly of the family is highly supported by both morphological and molecular data^16,176,177^ *Dinophilus* O. Schmidt, 1848 was recently recovered paraphyletic with respect to *Trilobodrilus* Remane, 1925, causing the erection of a third monophyletic dinophilid genus *Dimorphilus* Worsaae, Kerbl, Vang & Gonzalez, 2019 (sister to *Dinophilus* *sensu stricto* and *Trilobodrilus*). This genus so far includes two dimorphic species, *D. gyrociliatus* (O. Schmidt, 1857) and *D. kincaidi* (Jones & Ferguson, 1957), which are set apart from other dinophilids by having dwarf males and a short life cycle^16^. *Dimorphilus gyrociliatus* was originally described from tidal zone macroalgae in Naples, Italy, but has since then been reported from public aquaria and tidal habitats around the world, though these records most likely representing several cryptic species^16^. The present study is based on laboratory cultures originally established by B. Åkesson (University of Gothenburg) from collections of *D*. cf. *gyrociliatus*^178^ in the 1980’ies in Xiamen, China and currently kept in the lab of K. Worsaae (University of Copenhagen, since 2006), A. Hejnol (University of Bergen, since 2014) and J. M. Martín-Durán (Queen Mary University of London, since 2018).

*Dimorphilus gyrociliatus* male and female morphology diverges vastly, starting with egg size: male eggs are approximately 40 µm and female eggs 80 µm in diameter^179^. The roughly 1 mm long adult females possess six segments, with well-developed digestive, neural and muscular systems as well as segmentally arranged protonephridia, whereas the males are only 0.05 mm in length and lack a digestive tract. The nervous system is probably the best-studied organ system in both sexes: Females show an annelid-like nervous system with anterodorsal brain (with central neuropil and a surrounding layer of densely packed somata) and ventrolateral nerve cords, which are formed as one pair of ventrolateral nerve cords, two pairs of paramedian nerves and an unpaired ventromedian nerve, all connected by two to three transverse commissures per segment. Despite the male nervous system being reduced to only 68 neurons, it still comprises an anterior brain, two pairs of ventro-lateral nerves extending posteriorly, and a pair of posterior neuron clusters (penis ganglia) connected by a ventro-posterior commissure. No major differences are detected in size and overall shape of somata in male and female brains^46,180^, although the neurons may differ slightly in morphology, e.g., in the branching topology of the axons (Kerbl *et al.*, unpublished). Patterns of genes characterising and demarcating the brain/anterior neural region of adult females reflect the arrangement and expression of orthologous genes in larvae of macroscopic annelids or other Spiralians, though the different stages and lifestyles included in the analyses complicate exact comparisons^45^. Remarkably, the (morphologically apparently similar) neurons differ in the specific neurotransmitters they synthetize, with neurons in the brains of adult females seemingly not producing two to several of the (so far tested) neurotransmitters but being specific to one of them^46^. Surprisingly, there is a much higher overlap of individual neurotransmitter patterns in the nerves around the copulatory organ and penial ganglia of dwarf males^180^, thereby suggesting that neurotransmitter profiles of individual cells might be related to size and complexity of animals.

*Dimorphilus gyrociliatus* has direct development and a fast life cycle with females laying eggs within three weeks of their life span and living only for a few months. The fertilised eggs are deposited in cocoons containing female and male eggs in an approximate ratio of 3:1. Female embryos hatch with five body segments and add the last segment in the days after hatching. Males have a life span of little more than a week, hatching short before their female siblings and fertilising primarily their sisters within the cocoon before this is ruptured. This also infers a putative high degree of inbreeding in *D. gyrociliatus*. The early development of females has been studied in detail^47^, follows the general annelid pattern of spiral cleavage and formation of germ layers, and furthermore proposed homology of the second prostomial ciliary band to the prototroch in trochophore larvae of indirectly developing annelids.

**Supplementary Note 2: Phylogenetic position of *D. gyrociliatus*: background**

Since the discovery of *Dinophilus vorticoides* O. Schmidt, 1848, and thereby members of the family Dinophilidae, its relationship to and within annelids has been debated. Whereas its initial ascription to Platyhelminthes (O. Schmidt 1848) was soon rejected and the relationship to annelids suggested by, among others, their embryonic development^47^, the small and relatively simple body design and ciliated larval appearance of dinophilids have led to several speculations on their origin. Dinophilidae were originally regarded as ancestrally small “Archiannelida”^181-183^, but were later considered to be secondarily miniaturised forms, possibly originating through progenesis (in Gould’s sense^184^, involving accelerated maturation and early offset of somatic growth in a larval/juvenile stage of a macrofaunal ancestor)^50,185,186^. The latter hypothesis, though hard to test, is founded not only by the small size and lack of adult annelid features, but also by the resemblance of adult dinophilids to larval or juvenile stages of macrofaunal annelids (especially within Eunicida). These morphological comparisons focused on the configuration of ciliary patterns, reduced coelomic cavities and protonephridial systems, low segment number as well as the uncondensed architecture of the ventrolateral nerve cords^43,176,186,187^.

Larvae of some eunicidan families such as Dorvilleidae show strong resemblance to dinophilids in the overall shape of prostomium, body segments and pygidium as well as in the presence of segmental transverse ciliary bands. With the discovery of several heavily reduced meiofaunal genera of Dorvilleidae (e.g., *Apodotrocha* Westheide & Riser, 1983), this resemblance was strengthened and Dinophilidae was for a while depicted as highly derived dorvilleid, putatively having originated via a series of reduction events throughout the diversification of Dorvilleidae^188^. However, as also argued in several studies, the severe reduction in dinophilid morphology could also be the outcome of only one or few progenetic events, thereby indicating the family derived from other macrofaunal Eunicida^187,189-191^.

In contrast to these morphological considerations, early molecular phylogenetic studies^177,192^ already defied the close relationship of Dinophilidae to Dorvilleidae and Eunicida, which was also not found when incorporating more data and taxa in later phylogenomic studies^50,51^. Andrade *et al.* (REF^51^) observed Dinophilidae to either nest within Sedentaria or as sister to Pleistoannelida. Since this equivocality seemed to reflect confounding long branch effects, the authors did not regard the family’s position within Annelida as resolved. Struck *et al.* (REF^50^) addressed a different data set and suggested Dinophilidae to be placed with Orbiniida within Sedentaria (as sister group to Nerillidae), but again within a clade containing several long branches. David & Halanych (REF^193^) placed Dinophilidae in Sedentaria based on mitogenomic analyses of seven pleistoannelids, but remarkably, the dinophilid mitochondrial gene order (otherwise regarded to be a highly conserved gene order) differed from that of the other analysed species.

Herein, we find Dinophilidae to group with Lobatocerebridae, the two of them (=Dinophiliformia) constituting the sister group to Pleistoannelida in all analyses, except when using a simpler site-homogenous model (LG4X + R), which is instead inferring a relationship to Orbiniida within Sedentaria. Either position still supports Dinophilidae as secondarily reduced in size and complexity, yet this reduction may have occurred already in the origin of the common ancestor of Dinophiliformia to some degree, since the meiofaunal family Lobatocerebridae also exhibits reduced morphological complexity and size^194^. None of the former analyses of Dinophilidae’s position included Lobatocerebridae, which was only recently phylogenomically established as an annelid family^195^. This sister-family relationship was here found consistently and independent of the inclusion or exclusion of some of the other long branched taxa which Dinophilidae had previously shown affinity to, such as *Osedax*^51^. One may still argue that Lobatocerebridae also constitutes a long branch and their close relationship to Dinophilidae as well as the entire clade’s position next to Pleistoannelida are somehow affected by confounding long branch attraction effects. However, Dinophilidae is consequently placed outside Pleistoannelida throughout all analyses employing site-heterogenous models and with different taxon sampling. Interestingly, Dinophilidae do show some morphological resemblance to Lobatocerebridae besides them both being meiofaunal, e.g., in the common presence of highly separated ventral nerve cords and an unpaired median nerve as well as in a diverse set of mainly epidermal glands^194^. Still, Lobatocerebridae with its elongated, entirely ciliated body, multilobed brain, lack of apparent segmentation and different reproductive system differs starkly from Dinophilidae

103	R: A Language and Environment for Statistical Computing (R Foundation for Statistical Computing, Vienna, Austria, 2019).

104	RStudio: Integrated Development for R (RStudio, Inc, Boston, MA, 2019).

105	Haas, B. J. *et al.* *De novo* transcript sequence reconstruction from RNA-seq using the Trinity platform for reference generation and analysis. *Nat Protoc* **8**, 1494-1512, doi:10.1038/nprot.2013.084 (2013).

106	Eddy, S. R. *HMMER - biosequence analysis using profile hidden Markov models*, 2007).

107	Marletaz, F., Peijnenburg, K., Goto, T., Satoh, N. & Rokhsar, D. S. A New Spiralian Phylogeny Places the Enigmatic Arrow Worms among Gnathiferans. *Curr Biol* **29**, 312-318 e313, doi:10.1016/j.cub.2018.11.042 (2019).

108	Gonzalez-Dominguez, J., Liu, Y., Tourino, J. & Schmidt, B. MSAProbs-MPI: parallel multiple sequence aligner for distributed-memory systems. *Bioinformatics* **32**, 3826-3828, doi:10.1093/bioinformatics/btw558 (2016).

109	Criscuolo, A. & Gribaldo, S. BMGE (Block Mapping and Gathering with Entropy): a new software for selection of phylogenetic informative regions from multiple sequence alignments. *BMC Evol Biol* **10**, 210, doi:10.1186/1471-2148-10-210 (2010).

110	Nguyen, L. T., Schmidt, H. A., von Haeseler, A. & Minh, B. Q. IQ-TREE: a fast and effective stochastic algorithm for estimating maximum-likelihood phylogenies. *Mol Biol Evol* **32**, 268-274, doi:10.1093/molbev/msu300 (2015).

111	Smit, A. F. A. & Hubley, R. RepeatModeler Open-1.0. (2008-2015).

112	Smit, A. F. A., Hubley, R. & Green, P. RepeatMasker Open-4.0. (2013-2015).

113	Yang, G. MITE Digger, an efficient and accurate algorithm for genome wide discovery of miniature inverted repeat transposable elements. *BMC Bioinformatics* **14**, 186, doi:10.1186/1471-2105-14-186 (2013).

114	Lowe, T. M. & Eddy, S. R. tRNAscan-SE: a program for improved detection of transfer RNA genes in genomic sequence. *Nucleic Acids Res* **25**, 955-964, doi:10.1093/nar/25.5.955 (1997).

115	Llorens, C. *et al.* The Gypsy Database (GyDB) of mobile genetic elements: release 2.0. *Nucleic Acids Res* **39**, D70-74, doi:10.1093/nar/gkq1061 (2011).

116	Parra, G., Bradnam, K. & Korf, I. CEGMA: a pipeline to accurately annotate core genes in eukaryotic genomes. *Bioinformatics* **23**, 1061-1067, doi:10.1093/bioinformatics/btm071 (2007).

117	Stanke, M. & Waack, S. Gene prediction with a hidden Markov model and a new intron submodel. *Bioinformatics* **19 Suppl 2**, ii215-225 (2003).

118	Slater, G. S. & Birney, E. Automated generation of heuristics for biological sequence comparison. *BMC Bioinformatics* **6**, 31, doi:10.1186/1471-2105-6-31 (2005).

119	Haas, B. J. *et al.* Improving the *Arabidopsis* genome annotation using maximal transcript alignment assemblies. *Nucleic Acids Res* **31**, 5654-5666 (2003).

120	Wu, T. D. & Watanabe, C. K. GMAP: a genomic mapping and alignment program for mRNA and EST sequences. *Bioinformatics* **21**, 1859-1875, doi:10.1093/bioinformatics/bti310 (2005).

121	Haas, B. J. *et al.* Automated eukaryotic gene structure annotation using EVidenceModeler and the Program to Assemble Spliced Alignments. *Genome Biol* **9**, R7, doi:10.1186/gb-2008-9-1-r7 (2008).

122	ORFik: Open Reading Frames in Genomics v. R package version 1.6.9 (2020).

123	Altschul, S. F., Gish, W., Miller, W., Myers, E. W. & Lipman, D. J. Basic local alignment search tool. *J Mol Biol* **215**, 403-410, doi:10.1016/S0022-2836(05)80360-2 (1990).

124	Finn, R. D., Clements, J. & Eddy, S. R. HMMER web server: interactive sequence similarity searching. *Nucleic Acids Res* **39**, W29-37 (2011).

125	Petersen, T. N., Brunak, S., von Heijne, G. & Nielsen, H. SignalP 4.0: discriminating signal peptides from transmembrane regions. *Nat Methods* **8**, 785-786, doi:10.1038/nmeth.1701 (2011).

126	Krogh, A., Larsson, B., von Heijne, G. & Sonnhammer, E. L. Predicting transmembrane protein topology with a hidden Markov model: application to complete genomes. *J Mol Biol* **305**, 567-580, doi:10.1006/jmbi.2000.4315 (2001).

127	Ashburner, M. *et al.* Gene ontology: tool for the unification of biology. The Gene Ontology Consortium. *Nat Genet* **25**, 25-29, doi:10.1038/75556 (2000).

128	Powell, S. *et al.* eggNOG v3.0: orthologous groups covering 1133 organisms at 41 different taxonomic ranges. *Nucleic Acids Res* **40**, D284-289, doi:10.1093/nar/gkr1060 (2012).

129	Kanehisa, M., Goto, S., Sato, Y., Furumichi, M. & Tanabe, M. KEGG for integration and interpretation of large-scale molecular data sets. *Nucleic Acids Res* **40**, D109-114, doi:10.1093/nar/gkr988 (2012).

130	Danks, G. *et al.* OikoBase: a genomics and developmental transcriptomics resource for the urochordate Oikopleura dioica. *Nucleic Acids Res* **41**, D845-853, doi:10.1093/nar/gks1159 (2013).

131	Irimia, M. & Roy, S. W. Spliceosomal introns as tools for genomic and evolutionary analysis. *Nucleic Acids Res* **36**, 1703-1712, doi:10.1093/nar/gkn012 (2008).

132	Zdobnov, E. M. *et al.* OrthoDB v9.1: cataloging evolutionary and functional annotations for animal, fungal, plant, archaeal, bacterial and viral orthologs. *Nucleic Acids Res* **45**, D744-D749, doi:10.1093/nar/gkw1119 (2017).

133	Katoh, K. & Standley, D. M. MAFFT multiple sequence alignment software version 7: improvements in performance and usability. *Mol Biol Evol* **30**, 772-780, doi:10.1093/molbev/mst010 (2013).

134	Csuros, M. Malin: maximum likelihood analysis of intron evolution in eukaryotes. *Bioinformatics* **24**, 1538-1539, doi:10.1093/bioinformatics/btn226 (2008).

135	Csuros, M., Rogozin, I. B. & Koonin, E. V. A detailed history of intron-rich eukaryotic ancestors inferred from a global survey of 100 complete genomes. *PLoS Comput Biol* **7**, e1002150, doi:10.1371/journal.pcbi.1002150 (2011).

136	Paradis, E. & Schliep, K. ape 5.0: an environment for modern phylogenetics and evolutionary analyses in R. *Bioinformatics* **35**, 526-528, doi:10.1093/bioinformatics/bty633 (2019).

137	Braunschweig, U. *et al.* Widespread intron retention in mammals functionally tunes transcriptomes. *Genome Res* **24**, 1774-1786, doi:10.1101/gr.177790.114 (2014).

138	Tapial, J. *et al.* An atlas of alternative splicing profiles and functional associations reveals new regulatory programs and genes that simultaneously express multiple major isoforms. *Genome Res* **27**, 1759-1768, doi:10.1101/gr.220962.117 (2017).

139	Torres-Mendez, A. *et al.* A novel protein domain in an ancestral splicing factor drove the evolution of neural microexons. *Nat Ecol Evol* **3**, 691-701, doi:10.1038/s41559-019-0813-6 (2019).

140	Emms, D. M. & Kelly, S. OrthoFinder: phylogenetic orthology inference for comparative genomics. *Genome Biol* **20**, 238, doi:10.1186/s13059-019-1832-y (2019).

141	Huerta-Cepas, J., Serra, F. & Bork, P. ETE 3: Reconstruction, Analysis, and Visualization of Phylogenomic Data. *Mol Biol Evol* **33**, 1635-1638, doi:10.1093/molbev/msw046 (2016).

142	Frickey, T. & Lupas, A. CLANS: a Java application for visualizing protein families based on pairwise similarity. *Bioinformatics* **20**, 3702-3704, doi:10.1093/bioinformatics/bth444 (2004).

143	Larkin, M. A. *et al.* Clustal W and Clustal X version 2.0. *Bioinformatics* **23**, 2947-2948, doi:10.1093/bioinformatics/btm404 (2007).

144	Capella-Gutierrez, S., Silla-Martinez, J. M. & Gabaldon, T. trimAl: a tool for automated alignment trimming in large-scale phylogenetic analyses. *Bioinformatics* **25**, 1972-1973, doi:10.1093/bioinformatics/btp348 (2009).

145	Price, M. N., Dehal, P. S. & Arkin, A. P. FastTree 2--approximately maximum-likelihood trees for large alignments. *PLoS One* **5**, e9490, doi:10.1371/journal.pone.0009490 (2010).

146	Thiel, D., Franz-Wachtel, M., Aguilera, F. & Hejnol, A. Xenacoelomorph Neuropeptidomes Reveal a Major Expansion of Neuropeptide Systems during Early Bilaterian Evolution. *Molecular Biology and Evolution* **35**, 2528-2543, doi:10.1093/molbev/msy160 (2018).

147	Kanehisa, M. & Sato, Y. KEGG Mapper for inferring cellular functions from protein sequences. *Protein Sci* **29**, 28-35, doi:10.1002/pro.3711 (2020).

148	Castresana, J. Selection of conserved blocks from multiple alignments for their use in phylogenetic analysis. *Mol Biol Evol* **17**, 540-552, doi:10.1093/oxfordjournals.molbev.a026334 (2000).

149	Kerbl, A., Martin-Duran, J. M., Worsaae, K. & Hejnol, A. Molecular regionalization in the compact brain of the meiofaunal annelid *Dinophilus gyrociliatus* (Dinophilidae). *Evodevo* **7**, 20 (2016).

150	Blake, J. A., Grassle, J. P. & Eckelbarger, K. J. *Capitella teleta*, a new species designation for the opportunistic and experimental *Capitella* sp. I, with a review of the literature for confirmed records. *2009* **2**, 29, doi:10.11646/zoosymposia.2.1.6 (2009).

151	Grassle, J. & Grassle, J. F. Sibling species in the marine pollution indicator *Capitella* (polychaeta). *Science* **192**, 567-569, doi:10.1126/science.1257794 (1976).

152	Seaver, E. C., Thamm, K. & Hill, S. D. Growth patterns during segmentation in the two polychaete annelids, *Capitella* sp. I and *Hydroides elegans*: comparisons at distinct life history stages. *Evol Dev* **7**, 312-326, doi:10.1111/j.1525-142X.2005.05037.x (2005).

153	Meyer, N. P., Carrillo-Baltodano, A., Moore, R. E. & Seaver, E. C. Nervous system development in lecithotrophic larval and juvenile stages of the annelid *Capitella teleta*. *Front Zool* **12**, 15, doi:10.1186/s12983-015-0108-y (2015).

154	Oulion, S., Bertrand, S. & Escriva, H. Evolution of the FGF Gene Family. *Int J Evol Biol* **2012**, 298147, doi:10.1155/2012/298147 (2012).

155	Setiamarga, D. H. *et al.* An in-silico genomic survey to annotate genes coding for early development-relevant signaling molecules in the pearl oyster, *Pinctada fucata*. *Zoolog Sci* **30**, 877-888, doi:10.2108/zsj.30.877 (2013).

156	Putnam, N. H. *et al.* Sea anemone genome reveals ancestral eumetazoan gene repertoire and genomic organization. *Science* **317**, 86-94, doi:10.1126/science.1139158 (2007).

157	Buenrostro, J. D., Giresi, P. G., Zaba, L. C., Chang, H. Y. & Greenleaf, W. J. Transposition of native chromatin for fast and sensitive epigenomic profiling of open chromatin, DNA-binding proteins and nucleosome position. *Nat Methods* **10**, 1213-1218, doi:10.1038/nmeth.2688 (2013).

158	Picelli, S. *et al.* Tn5 transposase and tagmentation procedures for massively scaled sequencing projects. *Genome Res* **24**, 2033-2040, doi:10.1101/gr.177881.114 (2014).

159	Zhang, Y. *et al.* Model-based analysis of ChIP-Seq (MACS). *Genome Biol* **9**, R137, doi:10.1186/gb-2008-9-9-r137 (2008).

160	Li, Q., Brown, J. B., Huang, H. & Bickel, P. J. Measuring reproducibility of high-throughput experiments. *Ann. Appl. Stat.* **5**, 1752-1779 (2011).

161	Heinz, S. *et al.* Simple combinations of lineage-determining transcription factors prime cis-regulatory elements required for macrophage and B cell identities. *Mol Cell* **38**, 576-589, doi:10.1016/j.molcel.2010.05.004 (2010).

162	Murata, M. *et al.* Detecting expressed genes using CAGE. *Methods Mol Biol* **1164**, 67-85, doi:10.1007/978-1-4939-0805-9_7 (2014).

163	Langmead, B. & Salzberg, S. L. Fast gapped-read alignment with Bowtie 2. *Nature Methods* **9**, 357, doi:10.1038/nmeth.1923

<https://www.nature.com/articles/nmeth.1923#supplementary-information> (2012).

164	Haberle, V., Forrest, A. R., Hayashizaki, Y., Carninci, P. & Lenhard, B. CAGEr: precise TSS data retrieval and high-resolution promoterome mining for integrative analyses. *Nucleic acids research* **43**, e51, doi:10.1093/nar/gkv054 (2015).

165	Balwierz, P. J. *et al.* Methods for analyzing deep sequencing expression data: constructing the human and mouse promoterome with deepCAGE data. *Genome Biol* **10**, R79, doi:10.1186/gb-2009-10-7-r79 (2009).

166	Shikuma, N. J., Antoshechkin, I., Medeiros, J. M., Pilhofer, M. & Newman, D. K. Stepwise metamorphosis of the tubeworm *Hydroides elegans* is mediated by a bacterial inducer and MAPK signaling. *Proc Natl Acad Sci U S A* **113**, 10097-10102, doi:10.1073/pnas.1603142113 (2016).

167	Zwarycz, A. S., Nossa, C. W., Putnam, N. H. & Ryan, J. F. Timing and Scope of Genomic Expansion within Annelida: Evidence from Homeoboxes in the Genome of the Earthworm Eisenia fetida. *Genome Biology and Evolution* **8**, 271-281, doi:10.1093/gbe/evv243 (2015).

168	Kenny, N. J., Namigai, E. K., Marletaz, F., Hui, J. H. & Shimeld, S. M. Draft genome assemblies and predicted microRNA complements of the intertidal lophotrochozoans *Patella vulgata* (Mollusca, Patellogastropoda) and *Spirobranchus* (*Pomatoceros*) *lamarcki* (Annelida, Serpulida). *Mar Genomics* **24 Pt 2**, 139-146, doi:10.1016/j.margen.2015.07.004 (2015).

169	Li, Y. *et al.* Genomic adaptations to chemosymbiosis in the deep-sea seep-dwelling tubeworm *Lamellibrachia luymesi*. *BMC Biol* **17**, 91, doi:10.1186/s12915-019-0713-x (2019).

170	Naville, M. *et al.* Massive Changes of Genome Size Driven by Expansions of Non-autonomous Transposable Elements. *Curr Biol* **29**, 1161-1168 e1166 (2019).

171	Wang, S., Zhang, L., Meyer, E. & Bao, Z. Genome-wide analysis of transposable elements and tandem repeats in the compact placozoan genome. *Biol Direct* **5**, 18, doi:10.1186/1745-6150-5-18 (2010).

172	Gusev, O. *et al.* Comparative genome sequencing reveals genomic signature of extreme desiccation tolerance in the anhydrobiotic midge. *Nat Commun* **5**, 4784, doi:10.1038/ncomms5784 (2014).

173	Kelley, J. L. *et al.* Compact genome of the Antarctic midge is likely an adaptation to an extreme environment. *Nat Commun* **5**, 4611, doi:10.1038/ncomms5611 (2014).

174	Laricchia, K. M., Zdraljevic, S., Cook, D. E. & Andersen, E. C. Natural Variation in the Distribution and Abundance of Transposable Elements Across the *Caenorhabditis elegans* Species. *Mol Biol Evol* **34**, 2187-2202, doi:10.1093/molbev/msx155 (2017).

175	Westheide, W. & Riser, N. W. Morphology and phylogenetic relationships of the neotenic interstitial polychaete *Apodotrocha progenerans* n.gen., n.sp. (Annelida). *Zoomorphology* **103**, 67-87, doi:10.1007/BF00312239 (1983).

176	Müller, M. C. M. & Westheide, W. Comparative analysis of the nervous systems in presumptive progenetic dinophilid and dorvilleid polychaetes (Annelida) by immunohistochemistry and cLSM. *Acta Zoologica* **83**, 33-48, doi:10.1046/j.1463-6395.2002.00096.x (2002).

177	Struck, T. H., Westheide, W. & Purschke, G. Progenesis in Eunicida ("Polychaeta," Annelida)--separate evolutionary events? Evidence from molecular data. *Mol Phylogenet Evol* **25**, 190-199, doi:10.1016/s1055-7903(02)00231-2 (2002).

178	Wu, B. L. & Chen, M. On archiannelids of the Yellow Sea II: Dinophilidae and Nerillidae. *Oceanol Sin* **2**, 90-97 (1980).

179	Traut, W. Eine Mutante mit vergrößerten Männchen-Eiern bei Dinophilus gyrociliatus (Archiannelida). *Experientia* **22**, 237-238 (1966).

180	Kerbl, A., Winther Tolstrup, E. & Worsaae, K. Nerves innervating copulatory organs show common FMRFamide, FVRIamide, MIP and serotonin immunoreactivity patterns across Dinophilidae (Annelida) indicating their conserved role in copulatory behaviour. *BMC Zoology* **4**, 8, doi:10.1186/s40850-019-0045-x (2019).

181	Hatschek, B. Studie über die Entwicklungsgeschichte der Anneliden. Ein Beitrag zur Morphologie der Bilaterien. *Arb Zool Inst Univ Wien Zool Stat Triest* **1**, 277-404 (1878).

182	Hatschek, B. System der Anneliden, ein vorläufiger Bericht. *Lotos* **13**, 123-126 (1893).

183	Bubko, O. V. On systematic position of Oweniidae and Archiannelida (Annelida). *Zool Zhur* **52**, 1286-1296 (1973).

184	Gould, S. J. *Ontogeny and phylogeny*. (The Belknap Press of Harvard University Press, 1977).

185	Westheide, W. Progenesis as a principle in meiofauna evolution. *J. Nat. Hist.* **21**, 843-854 (1987).

186	Worsaae, K. & Kristensen, R. M. Evolution of interstitial Polychaeta (Annelida). *Hydrobiol* **535-536**, 319-340 (2005).

187	Westheide, W. in *The origin and relationships of lower invertebrates*	(eds S. Conway Morris, J. D. George, R. Gibson, & H. M. Platt) 310-326 (Oxford University Press, 1985).

188	Eibye-Jacobsen, D. & Kristensen, R. M. A new genus and species of Dorvilleidae (Annelida, Polychaeta) from Bermuda, with a phylogenetic analysis of Dorvilleidae, Iphitimidae and Dinophilidae. *Zoologica Scripta* **23**, 107-131, doi:10.1111/j.1463-6409.1994.tb00379.x (1994).

189	Westheide, W. *Ikosipodus carolensis* gen. et sp. n., an interstitial neotenic polychaete from North Carolina, U.S.A., and its phylogenetic relationship within Dorvilleidae. *Zool Scripta* **11**, 117-126 (1982).

190	Struck, T. H. Progenetic species in polychaetes (Annelida) and problems assessing their phylogenetic affiliation. *Integr Comp Biol* **46**, 558-568 (2006).

191	Purschke, G. in *Reproductive Biology and Phylogeny of Annelida*	(eds G. Rouse & F. Pleijel) 639-667 (Science Publishers, 2006).

192	Struck, T. H., Halanych, K. M. & Purschke, G. Dinophilidae (Annelida) is most likely not a progenetic Eunicida: evidence from 18S and 28S rDNA. *Mol Phylogenet Evol* **37**, 619-623, doi:10.1016/j.ympev.2005.07.010 (2005).

193	David, K. T. & Halanych, K. M. Mitochondrial genome of *Dinophilus gyrociliatus* (Annelida: Dinophilidae). *Mitochondrial DNA Part B* **2**, 831-832, doi:10.1080/23802359.2017.1407704 (2017).

194	Kerbl, A., Bekkouche, N., Sterrer, W. & Worsaae, K. Detailed reconstruction of the nervous and muscular system of Lobatocerebridae with an evaluation of its annelid affinity. *BMC Evol Biol* **15**, 277, doi:10.1186/s12862-015-0531-x (2015).

195	Laumer, C. E. *et al.* Spiralian phylogeny informs the evolution of microscopic lineages. *Curr Biol* **25**, 2000-2006, doi:10.1016/j.cub.2015.06.068 (2015).
